# Carbohydrate-active enzymes from *Akkermansia muciniphila* breakdown mucin O-glycans to completion

**DOI:** 10.1101/2024.03.27.586211

**Authors:** Cassie R. Bakshani, Taiwo O. Ojuri, Bo Pilgaard, Jesper Holck, Ross McInnes, Radoslaw P Kozak, Maria Zakhour, Sara Çakaj, Manon Kerouedan, Emily Newton, David N. Bolam, Lucy I. Crouch

**Affiliations:** Institute of Microbiology and Infection, College of Medical and Dental Sciences, University of Birmingham, Birmingham, B15 2TT; Protein Chemistry and Enzyme Technology Section, DTU Bioengineering, Department of Biotechnology and Biomedicine, Technical University of Denmark, 2800 Kgs, Lyngby, Denmark; Ludger Ltd, Culham Campus, Abingdon, OX14 3EB, UK; Biosciences Institute, Medical School, Newcastle University, Newcastle upon Tyne, NE2 4HH, UK

## Abstract

*Akkermansia muciniphila* is a human microbial symbiont residing in the mucosal layer of the large intestine. Its main carbon source is the highly heterogeneous mucin glycoprotein and *A. muciniphila* uses an array of Carbohydrate-active enzymes and sulfatases to access this complex energy source. Here we describe the biochemical characterisation of fifty-four glycoside hydrolases, eleven sulfatases, and one polysaccharide lyase from *A. muciniphila* to provide a holistic understanding of the carbohydrate-degrading activities. The results provide an extensive insight into the sequence of O-glycan degradation and how *A. muciniphila* can access this structurally variable substrate. One of the most outstanding elements of this work was the demonstration that these enzymes can act synergistically to degrade the O-glycans on the mucin polypeptide to completion, down to the core GalNAc. Additionally, human breast milk oligosaccharide, ganglioside, and globoside glycan structures were included in the study to understand the full degradative capability of *A. muciniphila*.

## Introduction

The mucosal surface of the human large intestine is predominantly comprised of gel-forming secreted mucins and approximately 80 % by dry weight of this glycoprotein is O-glycan^1, 2^. Mucin O-glycans include only five different monosaccharides, thus their considerable structural heterogeneity is attributed to linkage diversity between these monosaccharides and different sulfation patterns (Figure 1). The mucosal layer is the barrier between the host epithelial cells and the dense community of microorganisms residing in the colon^3^. Mucin-degrading activities are a key aspect of the complex interactions between host and microbe and under healthy conditions the production and breakdown of mucin is balanced^4^. However, disease states of the colon, which are on the rise, are typified by the mucosal layer being disrupted and changes in the proportion of mucinolytic bacteria^4, 5^. For *Akkermansia muciniphila*, multiple studies have shown its prevalence is inversely proportional to disease and inflammation markers^6–9^, but for the mucinophile *Ruminococcus gnavus* the opposite is observed and the population increases in disease^5^. It is estimated that 60 % of the human gut microbiota can access mucin as a nutrient source^10, 11^, yet a comprehensive understanding of this process and its links to disease has yet to be determined.

**Figure 1.**
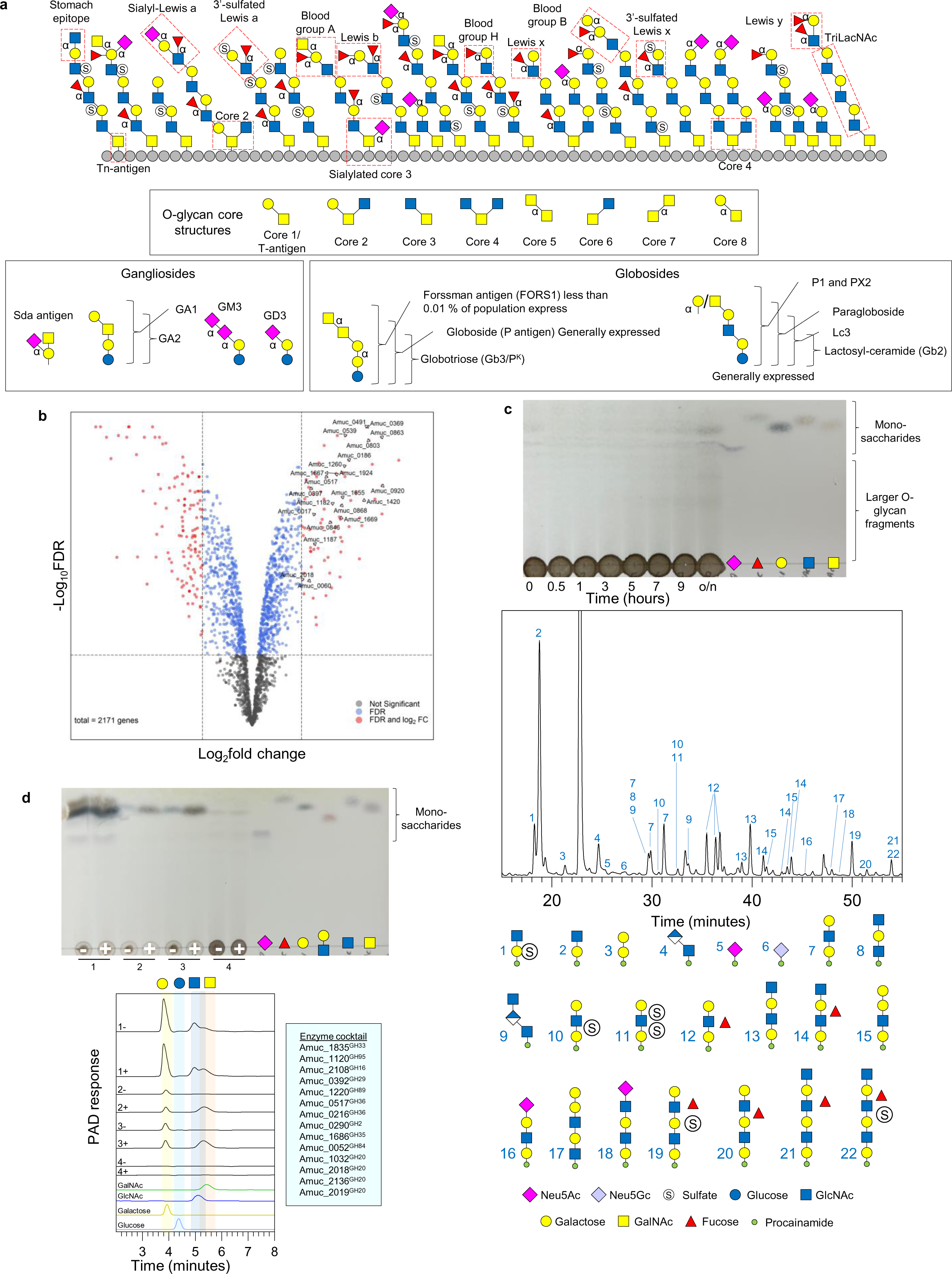
Activity of the recombinantly-expressed glycoside hydrolases from AM against α-linked monosaccharides. **a,** Structural features that are expected in natural secreted mucin glycoproteins with epitopes highlighted. Only α-linkages are labelled apart from the core GalNAc monosaccharides which are also alpha linkages. The structure of an O-glycan chain is generally accepted to be categorised into three sections: 1) the core, consisting of an α-linked GalNAc attached to a serine or threonine, 2) the polyLacNAc extensions linked to the core GalNAc, and 3) the terminal ‘capping’ epitopes. The polyLacNAc extensions are generally LacNAc disaccharides linked through β1,3-bonds, but variable sulfation, fucosylation, and branching adds to the complexity and heterogeneity along these chains. Key ganglioside and globoside structures are also included. **b,** Volcano plot highlighting the differential gene expression of AM when grown with PGMIII compared to glucose. Genes which did not pass the threshold of significance (FDR (Benjamini-Hochberg) < 0.05) are coloured grey. Genes which passed the threshold of significance (FDR < 0.05) but had a Log2 fold change between -1.5 and 1.5 are coloured blue. Genes which were significantly differentially expressed (FDR < 0.05) and had a Log2 fold change < -1.5 or > 1.5 are coloured red. Enzymes from this study that were significantly differentially expressed are labelled. The total number of genes in the analysis was 2171. **c,** Thin layer chromatography of whole cell assay of PGMIII-grown AM against fresh PGMIII. A smear can be seen increasing in concentration over time. The glycans in these samples were then labelled with procainamide and analysed by LC-FLD-ESI-MS. The chromatogram for the 9-hour sample is shown and the different glycan peaks are labelled. The data for all the time points is in Supplementary Figure 6. The data allows the reconstruction of the order of monosaccharides in an oligosaccharide but does not provide information about linkages. It is also not always possible to tell where a fucose or sulfate group are along the chain. **d,** A cocktail of enzymes from *A. muciniphila* BAA-835 can completely degrade the O-glycans from PGMIII down to the core GalNAc. Top and bottom panel are a TLC and HPAEC-PAD of the results, respectively. PGMIII was incubated with a cocktail of enzymes (in pale blue box) and the reaction stopped by boiling (1). The result of this reaction was a large amount of monosaccharides that can be seen on the TLC and by HPAEC-PAD. This reaction was then dialysed to remove all free monosaccharides and glycans (2) and concentrated (3). Untreated PGMIII was also included as a control (4). The application of Amuc_1008^GH31^ is indicated by plus signs and in samples 3 and 4, GalNAc can be seen to be released. Standards have also been included. Enzyme assays were carried out at pH 7, 37 °C, overnight, and with 1 μM enzymes.

It is the 20-year anniversary of the discovery of *Akkermansia muciniphila* from the relatively understudied Planctomycetota-Verrucomicrobiota-Chlamydiota (PVC) superphylum^12^. *A. muciniphila* is detectable in most people, colonises early on in life, and typically constitutes 1-3 % of the total microbiota^13–15^. *Akkermansia*-like sequences have also been detected in a wide variety of vertebrates which suggests a long evolutionary history between the mucosal surface of the gastrointestinal tract in vertebrates and *Akkermansia* species^13^. A biochemical characterisation of some of the Carbohydrate-Active enZymes (CAZymes) from *A. muciniphila* ATCC BAA-835 (AM) has been carried out in detail to provide insights into how AM tackles this complicated structure^16–20^. Furthermore, characterisation of glycopeptidases from AM have also illuminated how mucins are broken down^21–23^. However, a systematic approach to understanding mucin degradation has not previously been undertaken for any bacterial species and a significant number of enzymes from AM remain uncharacterised. Here, we provide a comprehensive picture elucidating how AM sequentially degrades mucin O-glycans, related structures, and other host carbohydrates.

## Results

### In vivo studies of AM with mucin

AM is highly restricted in terms of the substrates it can access, with its main carbon source being mucin. These observations were reproduced here (Supplementary Figure 1). RNA-Seq was performed on porcine gastric mucin III (PGMIII) to investigate the enzymes that AM uses to breakdown this nutrient source (Figure 1, Supplementary Data 1). We found that twenty CAZymes (from glycoside hydrolase (GH) families 2, 16, 20, 27, 29, 36, 89, 95, 97, 105, 109, and 123) and four sulfatases encoded in the AM genome were upregulated. These results were complementary to similar studies published previously (Supplementary Figure 2). A pangenome analysis was also undertaken to examine the prevalence of different CAZymes throughout strains and species (Supplementary Figure 3 & Supplementary Table 1). Strikingly, most enzymes were highly conserved between strains and close homologues were identified in other species.

Whole cell assays were used to assess enzyme activities on the surface of AM (Figure 1, Supplementary Figures 4-7). Results were initially assessed using thin layer chromatography (TLC), and a smear, with increasing in concentration over time, could be identified for PGMIII. This corresponded to the typical migration pattern for glycan fragments, rather than monosaccharides (Figure 1 and Supplementary Figure 4&5). Released glycan fragments were labelled with the fluorophore procainamide and detailed characterisation was performed using with liquid chromatography-fluorescence detection-electrospray-mass spectrometry (LC-FLD-ESI-MS). The data reveal that a range of mucin O-glycan fragments are produced by the enzymes on the surface of AM (Figure 1, Supplementary Figure 6). Galactose is typically present at the reducing end, indicative of GH16 endo-O-glycanase activity, and the degree of sulfation and fucosylation is variable^18^. We also used PGMIII-grown cells to carry out whole cell assays against several defined oligosaccharides (Supplementary Table 2). The most prominent activities were against TriLacNAc, human milk oligosaccharides (HMOs), Forssman antigen, and Lacto-N-biose. Whole cell assays of PGMIII-grown cells against bovine submaxillary mucin (BSM), which has sialylated Core 1 decorations^24^, showed only the release of sialic acids and this was confirmed by high-performance anion exchange chromatography with pulsed amperometric detection (HPAEC-PAD; Supplementary Figure 7).

### AM CAZymes can get down to the core GalNAc in PGMIII

Two putative GH31 enzymes are encoded in the AM genome. Amuc_1008^GH31^ is from subfamily 18 and other members of this subfamily, from *Bacteroides caccae*, *Phocaeicola plebius, Enterococcus faecalis, Clostridium perfringens, and Bombix mori* (domestic silk moth), have specificity for removing the α-GalNAc from peptide^25–27^. A phylogenetic tree of characterised bacterial GH31 enzymes shows clustering of activities and Amuc_1008^GH31^ clusters with the other characterised enzymes from GH31_18 (Supplementary Figure 8). We also found that Amuc_1008^GH31^ had specificity for the core α-GalNAc linked to peptide (BSM) with no activity against any other substrates (Supplementary Figure 9-13). Quantification of GalNAc release using HPAEC-PAD showed that Amuc_1008^GH31^ removed the highest concentration of all the enzymes tested here (Supplementary Figure 9). Notably, Amuc_1008^GH31^ could not hydrolyse Tn antigen, so therefore requires more than one amino acid for activity. For the more complex substrate PGMIII, Amuc_1008^GH31^ could not liberate GalNAc when tested in isolation, however when a cocktail of AM enzymes was used, GalNAc could then be released (Figure 1). This observation demonstrates that enzymes characterised from AM are capable of complete O-glycan degradation down to the polypeptide, which to our best knowledge has not been demonstrated before. This enzyme is predicted to be periplasmic and there was no obvious removal of GalNAc from BSM during whole cells assays, supporting this prediction (Supplementary Figure 7). A comparison of Amuc_1008^GH31^ model and the *E. faecalis* GH31_18 is provided in the Supplementary Discussion and Supplementary Figure 14.

### Hydrolysis of α-linked galactose from the non-reducing ends of PGMIII O-glycans

Galactose, GalNAc, GlcNAc, fucose and sialic acid all cap mucin O-glycans via α-linkages at the non-reducing end and prevalence of these monosaccharides will vary according to mucin type and genetics, for instance. Here, we determined which enzymes AM uses to tackle these different capping monosaccharides by using a panel of different substrates of varying complexity in overnight end-point assays.

There are two putative GH110 enzymes (Amuc_0480^GH110^ and Amuc_1463^GH110^) encoded by the AM genome, which share 28 % identity. These displayed activity towards blood group B (BGB) Types I and II and Galα1,3-Gal/GalNAc (Supplementary Figures 9-13). BGB is slightly less prevalent than BGA (10-40 %) in the human population, ranging between 0-30 %, depending on geographical location^28^. These are the only enzymes that are active against BGB from AM, which predicate the removal of fucose by the fucosidases. The structure of Amuc_1463^GH110^ has recently been solved^20^.

One putative GH27 (Amuc_1187^GH27^) is encoded in the AM genome and was active against α-linked galactose from the non-reducing end of most of the defined substrates tested here, except BG structures. Amuc_1187^GH27^ can hydrolyse Galα1,3Gal/GalNAc and globotriose, but P1 antigen could not be completely broken down in an end-point assay. This substrate specificity is discussed in the context of a model of Amuc_1187^GH27^ compared to solved structures of enzymes from *H. sapien* (Supplementary Discussion and Supplementary Figure 15).

AM also has three putative GH36 enzymes encoded in the AM genome, which have low sequence homology (Supplementary Table 3) and all three cluster in different locations on a phylogenetic tree when compared to characterised GH36 enzymes (Supplementary Figure 16). Amuc_0855^GH36^ has specificity for Galili antigen and globotriose in the defined substrate screen. Finally, AM also has one putative GH97 enzyme (Amuc_1420^GH97^), which displayed a preference for α-linked galactose in the defined substrate screen, except in the context of BGB. A model of Amuc_1420^GH97^ is discussed in the context of its activity (Supplementary Discussion and Supplementary Figure 17). All these enzymes were able to remove galactose from PGMIII, albeit in relatively small amounts, but this is likely due to the type of mucin used. The identity of the galactose was confirmed by HPEAC (Figure 2 and Supplementary Figure 18)

**Figure 2.**
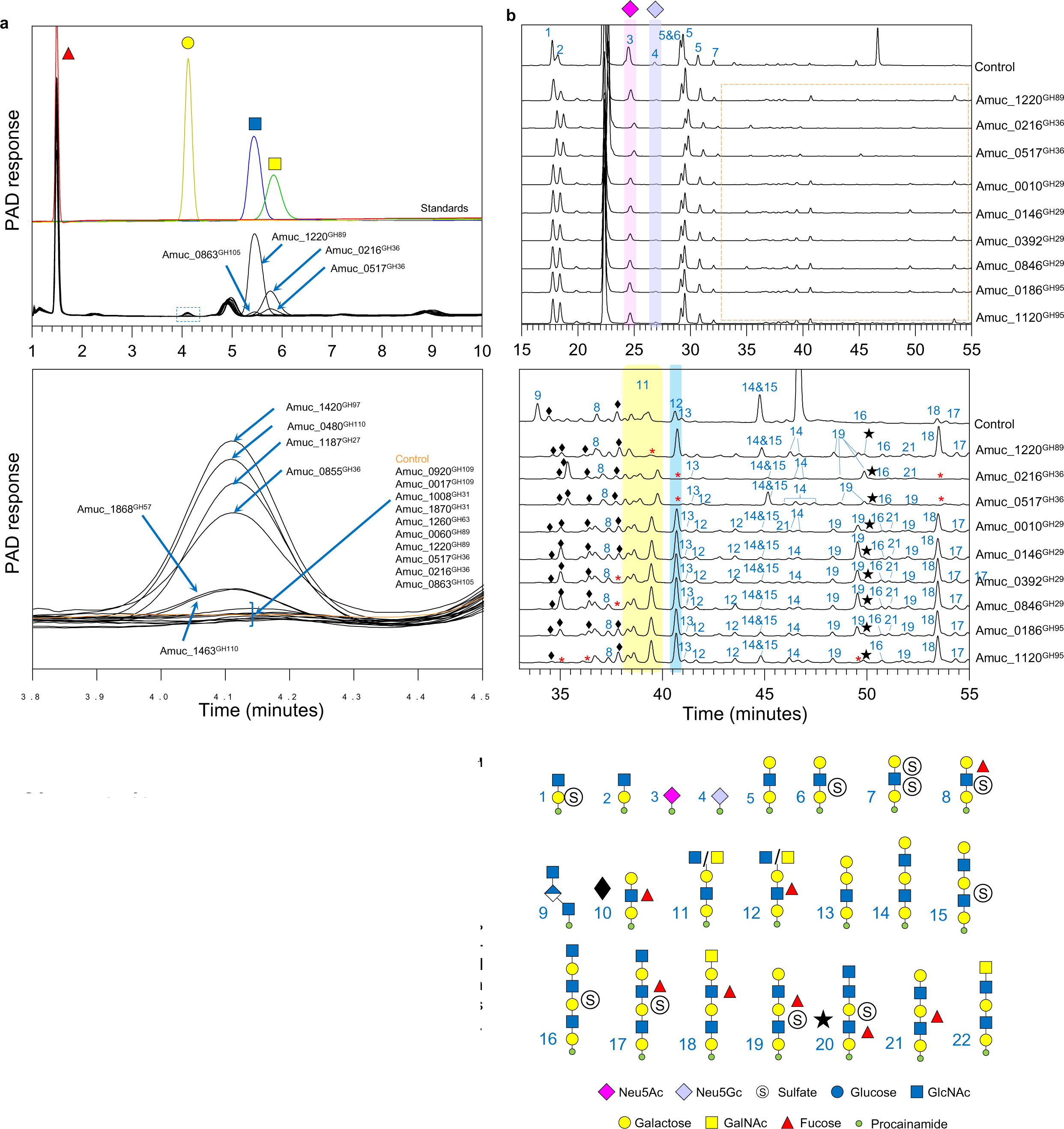
Activity of the recombinantly-expressed glycoside hydrolases from AM against α-linked monosaccharides capping mucin. **a,** Activity of enzymes against PGMIII (pre-treated with Amuc_1835^GH33^ and Amuc_1120^GH95^) and analysed using HPAEC-PAD. Standards were also run to identify monosaccharide products. Top panel: relevant area of chromatograms. Bottom panel: zoom in of where galactose elutes to show detail (dashed blue box in top panel). **b,** Activity of enzymes releasing α-GlcNAc, α-GalNAc, and α-fucose from GH16-released O-glycans from PGMIII. Top panel: full chromatograms with the two types of sialic acid highlighted in pink and purple. Bottom panel: zoom in of the smaller peaks (highlighted by the dashed orange box in the first panel). Black diamonds and black stars indicate glycan #10 and #20 peaks, respectively. Highlighted in yellow and blue are #11 and #12 peaks. The red asterisk indicated where a glycan is absent.

### Hydrolysis of α-linked GalNAc from the non-reducing ends of PGMIII O-glycans

The AM enzymes with activity against α-GalNAc capping structures are from families GH36 and GH109. All four enzymes were active against BGA structures, and they must remove this sugar before the fucosidases can act. Amuc_0216^GH36^ was able to act on all the substrates with α-GalNAc decorations we could test (Supplementary Figure 10-13). Notably, this was the only enzyme we found in AM to degrade Tn antigen. A phylogenetic tree of currently characterised bacterial GH36 enzymes, including those-from the AM genome described here, showed that Amuc_0216^GH36^ clustered with the other two examples of α-GalNAc’ases in the family (Supplementary Figure 16). One of these previously characterised enzymes also has activity on BGA and has been shown to convert BGA whole blood to universal type blood for transfusion applications^29^. Building on this application, recent work has also demonstrated that some of the AM CAZymes can be used to generate universal blood^20^. Amuc_0517^GH36^ was much narrower in its strict specificity for BGA only. There are two putative GH109 enzymes (Amuc_0017^GH109^ and Amuc_0920^GH109^) encoded by the AM genome, which have 66 % identity between them and display specificity towards α-GalNAc substrates apart from Tn antigen. When tested against PGMIII both GH36 enzymes could release relatively large amounts of GalNAc, with or without sialidase and fucosidase pre-treatment, and this was confirmed using HPAEC-PAD (Figure 2 and Supplementary Figure 18).

### Hydrolysis of α-linked GlcNAc from the non-reducing ends of PGMIII O-glycans

There are two putative GH89 enzymes (Amuc_0060^GH89^ and Amuc_1220^GH89^) encoded for by the AM genome and both only showed activity against GlcNAcα1,4-Gal disaccharide (stomach epitope), however Amuc_0060^GH89^ could not hydrolyse all the GlcNAcα1,4-Gal overnight, which suggests this is not the preferred substrate (Supplementary Figure 10-13). Amuc_1220^GH89^ was able to remove relatively large amounts of GlcNAc from PGMIII and this was confirmed using HPAEC-PAD (Figure 2 and Supplementary Figure 18). Further discussion on models of these enzymes in comparison to solved structures is presented in the Supplementary Discussion and Supplementary Figure 19.

### Hydrolysis of α-linked sialic acid and fucose from the non-reducing ends of PGMIII O-glycans

AM has three sialidases and six fucosidases that together can tackle a wide range of substrates and their specificities have been characterised in depth previously^16, 17^. The specificities observed for these enzymes presented in this report are comparable to previous characterisations, but we were able to identify notable further observations (Supplementary Figures 20-22). In terms of the sialidases, we found that the two GH33 enzymes from AM can act on the relatively complex GD1a and GT1b ganglioside structures and Sda antigen, which are common features of the human glycome^30, 31^. In terms of the fucosidases, Amuc_0010^GH29^ exhibits specificity towards type II BG structures (not type I) and the lacto-N-*neo*tetraose HMO series (not the lacto-N-tetraose), whereas Amuc_1120^GH95^ can hydrolyse fucose from both type I and II structures. This specificity facilitates the probing of mucin glycan structures in more detail. For instance, only Amuc_1120^GH95^ can hydrolyse fucose from untreated PGMIII, indicating that this substrate only has type I structures immediately available to the fucosidases at the non-reducing ends of the O-glycans (Supplementary Figure 18). A full discussion of the characterisation of these enzymes is described in the Supplementary discussion.

### Combining an AM GH16 endo-O-glycanase with CAZymes hydrolysing the α-capping monosacchairdes

To further explore the specificities of the CAZymes removing α-linked monosaccharides from PGMIII, we used a series of sequential reactions, which were then labelled with procainamide and analysed by LC-FLC-ESI-MS (Figure 2 and Supplementary Figure 23). Complementary to the relatively high monosaccharide release seen by HPAEC-PAD, the LC-FLC-ESI-MS data showed that Amuc_1220^GH89^, Amuc_0216^GH36^, and Amuc_0517^GH36^ could act on GH16-derived O-glycan fragments (Figure 2). These assays, therefore, provide the ability to characterise the fragments produced by the GH16. For example, glycan #11, that is absent in the Amuc_1220^GH89^ assay, has a GlcNAcα1,4-Gal capping structure. This also confirms that the GH16 can accommodate this epitope in its active site. Amuc_0216^GH36^ and Amuc_0517^GH36^ both acted on glycans #12 and 18, confirming that they have α-GalNAc caps. Furthermore, we could also observe four of the fucosidases acting on GH16-derived #10 O-glycan fragments. The different peaks corresponding #10 glycan compositions will be different combinations of linkages and positioning of the fucose. For instance, the structures hydrolysed by Amuc_1120^GH95^ are not hydrolysed by Amuc_0010^GH29^, thus confirming them as type 1 α1,2-fucose structures. In addition, Amuc_0392^GH29^ and Amuc_0846^GH29^ both hydrolyse a different glycan #10 (eluting at a different time to the glycan #10 acted on by Amuc_1120^GH95^) complementary to their comparable activities seen in the defined substrate screen.

### Investigating β-galactosidase activity

The combination of GH16 endo-O-glycanase activity and removal of α-capping monosaccharides from the non-reducing ends leads us to how the remaining O-glycan fragments will be broken down. This will require contribution from β-galactosidases, β-HexNAc’ases, fucosidases, and sulfatases. There are nine putative CAZymes encoded in the AM genome that were highlighted as possible β-galactosidases from families GH2, GH35, and GH43 subfamily 24 (Supplementary Table 4). Recombinant enzymes were screened against a variety of substrates to determine their specificity (Figure 3; Supplementary Figure 24-27). The results revealed a range of specificities, but between them, the β-galactosidases could breakdown all the defined oligosaccharides tested. There are examples of very broad-acting (Amuc_0290^GH2^) and highly specific β-galactosidases (Amuc_0539^GH2^ was found to be specific to β1,4-linked Gal with either a Gal or GalNAc in the +1 position). None of these enzymes showed activity towards 3-FL, Lewis A, or Lewis X based structures, which confirms that there is a strict order of degradation, with fucosidases acting on these substrates first, followed by galactosidases.

**Figure 3.**
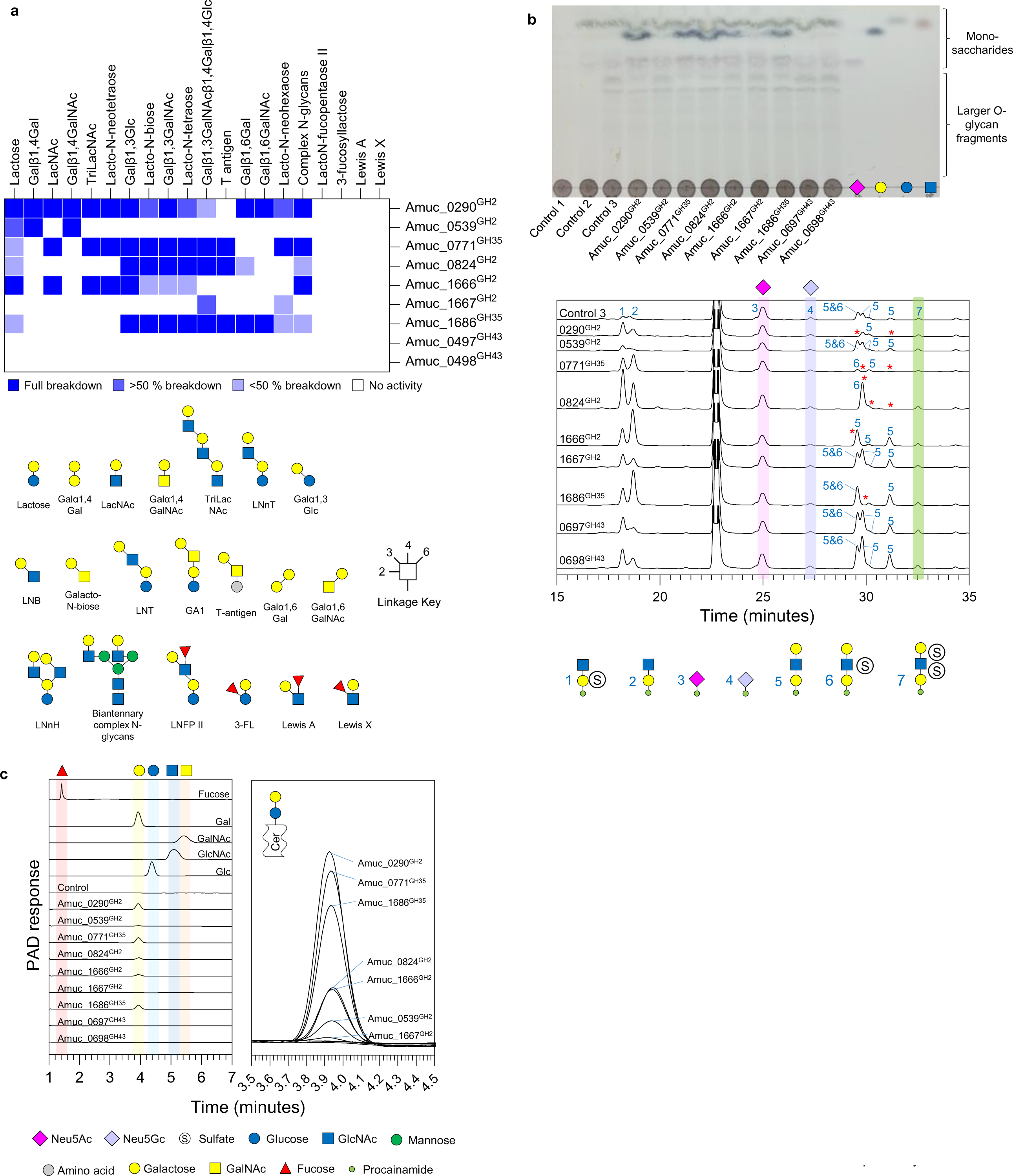
Activity of the glycoside hydrolases from families 2, 35, and 43 from AM against β-linked galactose substrates. **a,** Heat map of recombinant enzyme activities against defined oligosaccharides. The dark blue and white indicate full and no activity, respectively, and partial activities are represented by the lighter blues. Partial activity is when all the substrate has not been broken down in an end-point assay. **b**, The activity of the panel of β-galactosidases against PGMIII that had been sequentially degraded. Control 1 – no enzymes added, Control 2 - Amuc_1835^GH33^ and Amuc_1120^GH95^, Control 3 – Control 2 + Amuc_2108^GH16^. The top and bottom panels are the TLC and the LC-FLD-ESI-MS results, respectively. These assays were performed in stages, so the reactions were boiled in between steps. Enzyme assays were carried out at pH 7, 37 °C, overnight, and with 1 μM enzymes. Monosaccharide standards are shown in the right-hand lanes. **d**, Activity of the GH2/35/43 enzymes against Lactosylceramide. The assays were analysed using HPAEC-PAD and the different controls confirm the release of galactose. Left panel – the different controls and samples are stacked. Right panel – The assay chromatograms are overlaid for comparison so the galactose peaks can be observed in more detail. The substrate could not be resolved using this method, so whether the reaction had gone to completion could not be determined and thin layer chromatography of the samples was also inconclusive. Enzyme assays were carried out at pH 7, 37 °C, overnight, and with 1 μM enzymes.

We tested this panel of β-galactosidases against PGMIII that had been sequentially treated with different AM CAZymes. It should be noted that, although with the addition of only fucosidase, sialidase, and a GH16 a range sizes and compositions of O-glycan fragments were produced, the majority of these were di- and tri-saccharides of alternating galactose and GlcNAc, ascertained by the intensity of the fluorescence signals being relatively high for these structures (Figure 3 and Supplementary Figure 28). When testing the panel of β-galactosidases, the sample was pre-treated with the three enzymes that remove α-linked GlcNAc and GalNAc (Amuc_1220^GH89^, Amuc_0216^GH36^, and Amuc_0517^GH36^) and a fucosidase (Amuc_0392^GH29^) to maximise the O-glycans with β-galactose at the non-reducing ends. A TLC of these assays clearly shows galactose being released for five of the enzymes (Figure 3, blue stained bands). These samples were labelled with procainamide and analysed by LC-FLD-ESI-MS (Figure 3 and Supplementary Figure 28). Activities for four of the β-galactosidases was observed against the trisaccharides. Amuc_0539^GH2^ showed no activity and this supports the observation of only being active when a Gal or GalNAc is in the +1 subsite. Notably, the doubly-sulfated trisaccharide glycan #7 was not acted on by any β-galactosidase. The GH2 and GH35 enzymes also showed activity against lactosylceramide (Ganglioside and Globoside core structure), with Amuc_0290^GH2^, Amuc_0771^GH35^, and Amuc_1686^GH35^ releasing the most galactose (Figure 3). Further discussion on the activities of the β-galactosidases is in the Supplementary Discussion and Supplementary Figures 29-30.

### Investigating β-HexNAc activities

The GH20 family is the largest represented family in AM, with eleven putative enzymes encoded with generally low identity between sequences (Supplementary Figure 31 and Supplementary Table 5). The activity predominantly observed in this family is exo-acting β-GlcNAc’ases, but activity on GalNAc has also been observed. We also included Amuc_0052^GH84^, Amuc_0803^GH123^, and Amuc_2109^GH3^, with the latter having been shown to have relevant activities (pNPGlcNAc>GalNAc)^32^ and clusters with other β-HexNAc’ases on a phylogenetic tree when characterised enzymes were compared (Supplementary Figure 32). In summary, we found a range of different β-HexNAc’ase specificities encoded for by the AM genome and the specificities of these enzymes are generally towards the glycan structures found in mucins rather than complex N-glycans or chitin. Between them, these β-HexNAc’ases could access all the defined substrates provided (Figure 4 and Supplementary Figures 33-35). The panel of β-HexNAc’ases were then tested against PGMIII that had been sequentially treated (controls 1-4; Supplementary Figure 23). The samples were then labelled with procainamide and analysed by LC-FLD-ESI-MS. The predominant glycans in ‘Control 4’ were two disaccharides, one sulfated on the galactose and one not (Figure 4). Four of the β-HexNAc’ases were able to breakdown the sulfated disaccharide, providing further insight into the different specificities of these enzymes. To further explore the capacity of β-HexNAc’ases against host glycan structures, we used the AM CAZymes to prepare Sda antigen from the ganglioside GD1a. Intriguingly, out of the five β-HexNAc’ases that could act on GA2, only Amuc_1924^GH20^ was able to accommodate the branching sialic acid to access the GalNAc present in Sda antigen in an end-point reaction (Figure 4). In addition to the sequential degradation of mucin O-glycans, the activity of the AM CAZymes against HMOs was also explored and is detailed in the Supplementary Discussion and Supplementary Figures 36-37.

**Figure 4.**
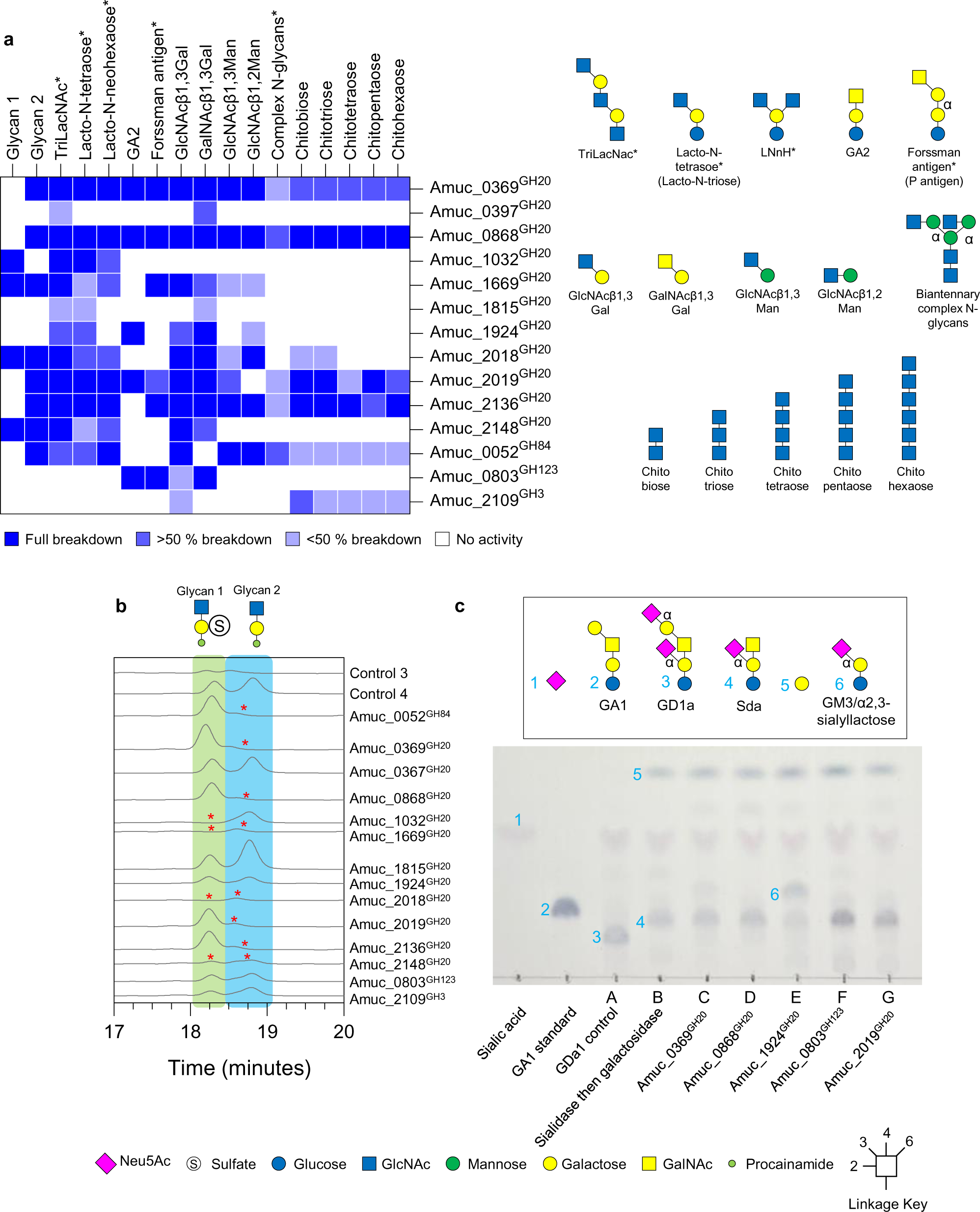
Activity of β-HexNAc’ases from glycoside hydrolases families 20, 84, 123 and 4 from AM. **a,** Heat map of recombinant enzyme activities against defined oligosaccharides. The dark blue and white indicate full and no activity, respectively, and partial activities are represented by the lighter blues. Asterisk represents those substrates that were generated by pre-treating with other CAZymes. **b**, The activity of the panel of β-HexNAc’ases against PGMIII that had been sequentially degraded. Control 1: no enzymes added, Control 2: Amuc_1835^GH33^ and Amuc_1120^GH95^, Control 3: Control 2+Amuc_2108^GH16^, and Control 4: Control 3+Amuc_0771^GH35^. The relevant area of the chromatograms of the LC-FLD-ESI-MS data is shown and the two glycans are highlighted. Red asterisk indicate where a glycan is not present. These assays were performed in stages, so the reactions were boiled in between steps. Enzyme assays were carried out at pH 7, 37 °C, overnight, and with 1 μM enzymes. **c**, TLC results of sequential assays against ganglioside GD1a. GDa1 was sequentially treated with Amuc_1835^GH33^ and then Amuc_0771^GH35^ to (Lane B). The sample was then boiled and a panel of β-HexNAc’ases added.

### Tackling sulfated substrates

We highlighted twelve potential sulfatases from the AM genome using upregulation data^4,33–35^ and SulfAtlas database^36^. These enzymes could be assigned to seven different families and one unknown, which ultimately turned out not to be a sulfatase (Supplementary Table 1). The different subfamilies cluster on a phylogenetic tree and the activities (mainly explored in *Bacteroides thetaiotaomicron*) within a subfamily correspond those already described (Supplementary Figure 37)^37, 38^. The sequence identity between the sulfatases is low, with the highest being ∼60 % for pairs in the same subfamily (Supplementary Table 8). The defined sulfated substrates tested were sulfated monosaccharides and Lewis structures (Supplementary Figure 38). We first screened the sulfatases against the sulfated monosaccharides and found Amuc_1655^Sulf16^ and Amuc_1755^Sulf16^ displayed activity against 4S-Gal, Amuc_1755^Sulf16^ also displayed activity against 4S-GalNAc. The subfamily 11 enzymes Amuc_1033^Sulf11^ and Amuc_1074^Sulf11^ displayed activity against 6S-GlcNAc and Amuc_0121^Sulf15^ displayed partial activities against 6S-Gal and 6S-GalNAc. These specificities align with those previously observed for these subfamilies^37^.

We then used combinations of the sulfatases, α-fucosidases and β-galactosidase encoded in the AM genome to determine how the different sulfated Lewis structures are broken down (Supplementary Figures 39-40). For the 3’S and 6S-Lewis A and X glycans, Amuc_0392^GH29^ removes the fucose to produce 3’S and 6S-LacNAc and LNB. From here, the 3’S disaccharides are then acted on by sulfatases from subfamily 20. In contrast, the 6S disaccharides are acted on by the β-galactosidases first. In this study, we could not find a way to breakdown the 6’S-Lewis structures. Previous activities against these substrates have been observed for BT1624 from subfamily 15^37^, so we had predicted that Amuc_0121^Sulf15^ may be able to do this but only partial activity against sulfated galactose could be detected.

### The unexpected and unsolved

There are two observations from this study worthy of note, concerning the ability of AM to colonise the mucosal surface and interact with other members of the microbiota. Firstly, AM did not grow directly on any of the glycosaminoglycans (GAGs) tested (Supplementary Figure 1), however, its genome encodes Amuc_0778^PL38^ and Amuc_0863^GH105^ and activity against all GAGs except heparin sulfate was observed for these enzymes. Secondly, AM grew on a high-mannose N-glycoprotein and, by analysing substrates remaining in the broth, we deciphered that AM was using the protein component and leaving the glycan. A full discussion on why AM has these features is in the Supplementary Discussion and Supplementary Figures 42-46.

In total, we were unable to find activities for eleven of the putative CAZymes and three of the sulfatases during this work (Supplementary Table 1 and Supplementary Discussion). Some of these CAZymes are from the GH13 and 77 families, which are currently solely associated with breaking down α-glucose polymers (Supplementary Figure 47).

## Discussion

The human gut microbiota plays an intrinsic role in health and disease, with impacts now recognised to extend to many other parts of the body. One of the key processes occurring at the interface between microbes and host is the breakdown of mucins and this holistic investigation of AM CAZymes described here is an important step forward in understanding this complex relationship. Previously, we characterised the GH16 endo-O-glycanases from AM (Amuc_0724^GH16^, Amuc_0875^GH16^ Amuc_2136^GH20^) that hydrolyse within the polyLacNAc chains of mucin to produce fragments of O-glycans^18^. Here, we expand on this work significantly by taking a systematic approach to assessing the full enzymology of this organism. The results have allowed an order of enzyme activities on different substrates to be assembled (Figure 5), sequential breakdown of complex substrates to be demonstrated, and determination that a cocktail of AM CAZymes could be applied to reach the core GalNAcs of a complex mucin substrate. A model of how AM degrades mucin can now be proposed (Figure 6).

**Figure 5.**
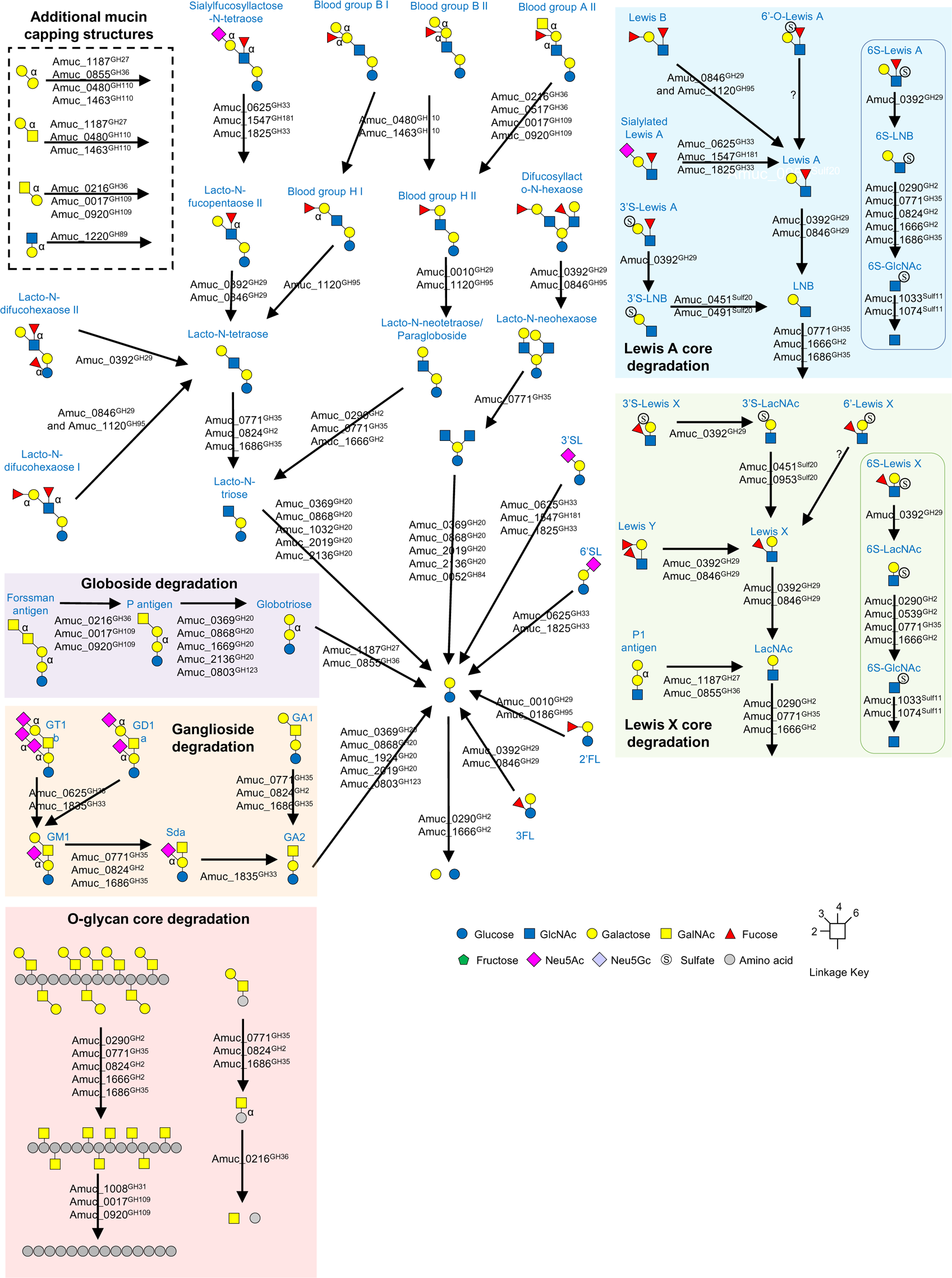
Illustration of the CAZymes characterised in this report against host glycans by AM. The different type of substrates are grouped where possible. The enzymes listed for each reaction are the ones that will work alone, but for the Lewis B structures the “and” signifies that both enzymes are required to remove the fucose. For the SFLNT, the possible fucosidase activities were not included, but Amuc_0392^GH29^ can act on this substrate also. Only alpha linkages are labelled apart from the core GalNAc monosaccharides which are also alpha linkages.

**Figure 6.**
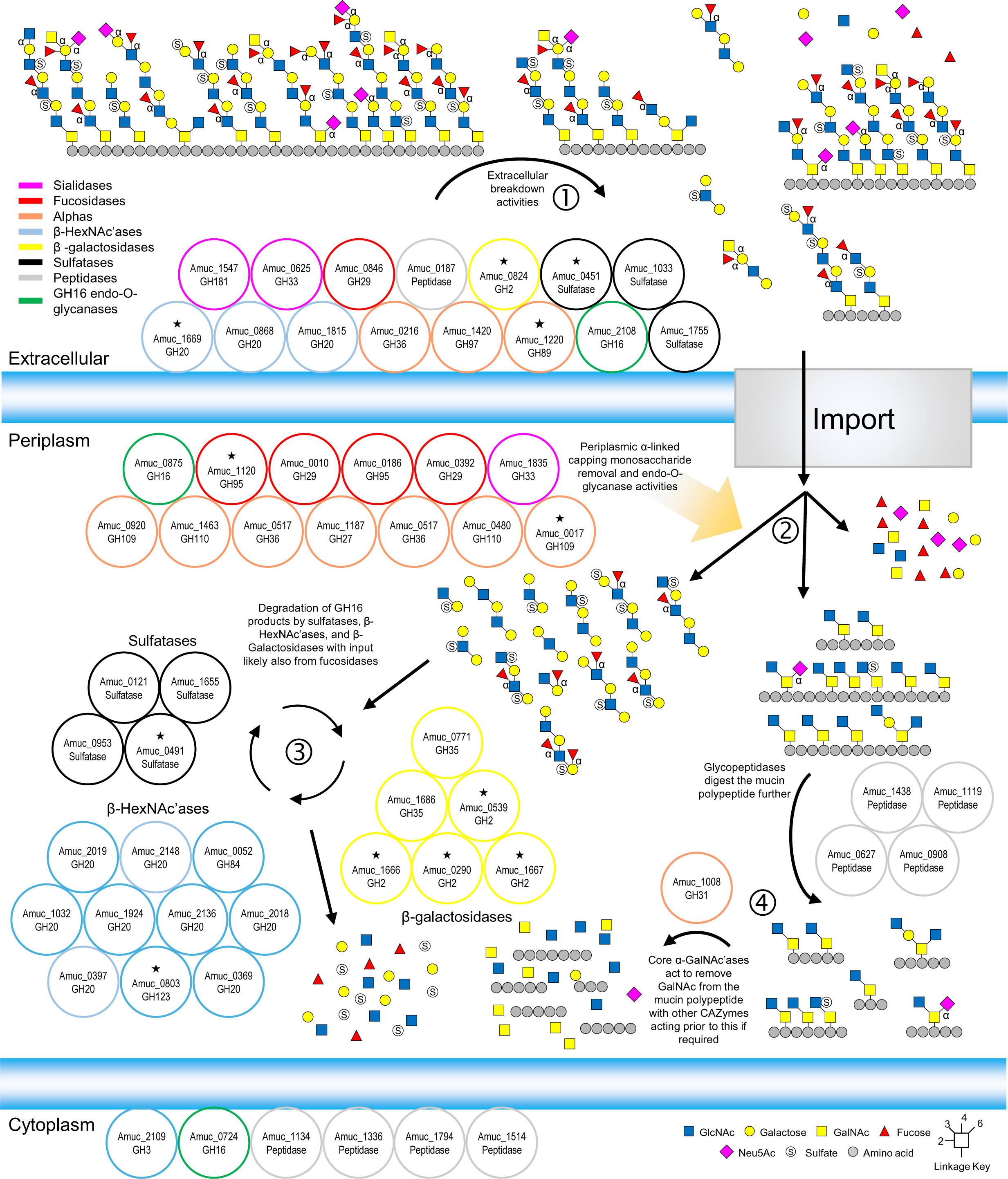
A model for the degradation of mucins by *Akkermansia muciniphila* based on current understanding. The GH enzymes included are only where activity has been observed and the colour indicated the type of activity (see key). The localisation of all the enzymes is based on the SignalP 6.0 prediction, apart from Amuc_2108^GH16^ due to it being observed in outer membrane analysis when AM was grown on mucin (Ottman *et al.* 2016) and our whole cell assays support a GH16 being localised to the outside of the cell. The peptidases included are both those that have been characterised and those that have been highlighted as upregulated on mucin. The signal sequences of a peptidase and sulfatases potentially localise them to the outside of the cell also. The numbers indicate the order that mucin is degraded. 1:There is some processing of mucin on the surface, with both exo- and endo-GH activities. Large sections of mucin are then imported into the cell for further breakdown. 2:Initially, there will be further exo- and endo-GH activities to produce fragments of O-glycans and mucin polypeptide with only core glycan decoration remaining. 3:The O-glycan fragments will then be broken down to monosaccharides through the alternating action of sulfatases, fucosidases, β-galactosidases, and β-HexNAc’ases. 4:The remaining glycopeptides are known targets for characterised *A. muciniphila* glycopeptidases and Amuc_1008^GH31^ will remove core GalNAc from the polypeptide. The stars indicate Mul1A association.

The glycomes of different humans vary and the capacity of the AM CAZymes to breakdown most of the substrates tested reveals how AM has adapted to accessing mammalian-derived glycans. Blood groups are a recognisable example, which varies globally^39^, but Galα1,4Gal epitopes of P1 and P^k^ antigens are also generally present in the mucosal surface of the human large intestine. These glycans, and others like them, are receptors for a variety of pathogens and their toxins, such as Shiga toxin^40^. Forssman antigen is normally present in only 0.01 % of the human population but in that relatively small population this antigen would be present in the GI tract^41^. It can change susceptibility to some diseases and is also expressed in some cancers, but its true prevalence and impact on different diseases is understudied^42^. Characterising enzymes with specificity towards these structures enables research into these epitopes much more broadly. One of the major impacts of this work will be the ability to characterise different mucins in much more detail and in a higher throughput manner. We hope that these enzymes provide new glycotools for the community and have far-reaching applications, such as monitoring the change in mucin O-glycans between healthy and diseased tissue, or the detection of disease-associated epitopes.

## Materials and Methods

### Substrates

A list of substrates and their sources are listed in Supplementary Table 2. Porcine gastric mucin III (Sigma) was prepared by dissolving overnight in sterile DI water. This is too turbid for bacterial growths, so the precipitate is removed by centrifugation at 1,500 x g for 5 minutes in sterile 15 mL falcon tubes. Polyglucuronic acid was prepared as described^43^.

### Recombinant protein expression and purification

Recombinant plasmids were transformed into Tuner cells (Novagen) in Luria-Bertani (LB) broth containing with 50 or 100 μg/mL kanamycin or ampicillin, respectively. The plates were used to inoculate 1 L flasks (also including relevant antibiotics) and grown at 37 °C with shaking at 180 rpm to mid-exponential, this was then cooled to 16 °C, and isopropyl β-D-thiogalactopyranoside (IPTG) was added to a final concentration of 0.2 mM. Recombinant His-tagged protein was purified from cell-free extracts using immobilized metal affinity chromatography (IMAC using Talon resin; Takara Bio). The buffer used during the purification process was 20 mM Tris-HCl, 100 mM NaCl, pH 8. Cell-free extracts (∼30 mL) were passed through a 5 mL bed volume of resin. This was then washed with 50 mL of buffer and the protein eluted with 10 mM and then 100 mM imidazole. The purification is assessed by sodium dodecyl sulfate–polyacrylamide gel electrophoresis (SDS-PAGE; precast 4 to 20 % gradient from Bio-Rad and stained with Coomassie Brilliant Blue), but proteins typically elute in the 100 mM fraction. The proteins were then concentrated using centrifugal concentrators (Vivaspin®, 10 kDa MWCO), the absorbance determined at 280 nm (NanoDrop™ One, Thermo Scientific), and concentrations calculated using extinction coefficients taken from ProtParam.

### Recombinant enzyme assays

The concentrations of defined oligosaccharide substrates used in the assays are presented in Supplementary Table 2. The activities of the recombinant enzymes were typically assessed in 20 mM 4-morpholinepropanesulfonic acid (MOPS, pH 7) at 37 °C, and a final enzyme concentration of 1 μM. The volume varies according to how much was required for subsequent analysis. For the sulfatase assays, 5 mM of CaCl was included. PNGaseL (LZ-PNGaseL-50-KIT) was used where required.

### Thin layer chromatography

For defined oligosaccharides, 3 μL of an assay containing 1 mM substrate was spotted onto silica plates (Sigma Z740230). For assays against glycoproteins, GAGs and starch substrates, this was increased to 9, 6, and 9 μL, respectivley. The plates were resolved in running buffer containing butanol/acetic acid/water. Different ratios were used for different substrates - 2:1:1 (typically resolved once) was used for most of the defined oligosaccharides and 1:1:1 (typically resolved twice) was typically used for substrates where large products were expected (e.g., PGMIII). All TLCs were stained using a diphenylamine-aniline-phosphoric acid (DPA) stain^44^.

### Growth of AM on a variety of substrate

AM was grown on chopped meat broth (CMB) as described in^18^. Overnight starter cultures were inoculated from a glycerol stock into 5 mL of chopped meat broth (CMB) and grown in an anaerobic cabinet (Whitley A35 Workstation, Don Whitley Scientific). Growth curves were collected in 96-well plates using Cerillo Stratus plate readers. 300 μL of 2x CMB (minus the monosaccharides) was mixed with 300 μL substrate and 200 μL was pipetted into three wells to provide replicates. Growth curves were also repeated with different starter cultures and on different days.

### RNA-seq of AM

AM was grown overnight in CMB at 37°C under anaerobic conditions. The culture was diluted 1:20 into CMB minus monosaccharide with the addition of either glucose or mucin and grown to an OD of 0.5 at 600 nm. Cells were pelleted by centrifugation at 16,000 x g for 30 seconds and flash frozen using a dry ice and ethanol bath. Sample processing, library construction, and sequencing were performed by Azenta. Sequencing adapter removal and read quality trimming were performed with fastp v.0.23.2 using default parameters^45^. Reads were mapped to the AM (GCF_000020225.1) and enumerated using kallisto v.0.46.2^46^. Differential gene expression analyses were performed with Voom/limma v.3.40.6^47^ implemented in Degust v.4.2-dev (https://zenodo.org/records/3501067). The volcano plot was created using the package EnhancedVolcano v.1.22.0 (https://github.com/kevinblighe/EnhancedVolcano) in R v.4.4.0.

### Whole cell assays

Overnight starter cultures were inoculated from a glycerol stock into 5 mL of CMB and grown under anaerobic conditions. This was then used to inoculate a 5 mL culture of AM in CMB (minus the monosaccharides) with substrate and cells were pooled, harvested at mid-exponential, washed twice with PBS, and 250 μL of 2x PBS was used per 5 ml culture to resuspend cells. The assays were then mixed 50:50 cells to substrate, where substrate concentrations were the same as used in the recombinant enzyme assays. PGMIII, BSM, hyaluronic acid, chondroitin sulfate, RNaseB, and *Sc*mannan assays were 500 μL total and 50 μL samples were taken for each time point. Assays with defined oligosaccharides and HMOs were 200 μL and 20 μL samples were taken for each time point. Assays were carried out at 37 °C and reactions were terminated by boiling samples.

### High-performance anion exchange chromatography with pulsed amperometric detection

To analyse monosaccharide release, sugars were separated using a CARBOPAC PA-1 anion exchange column with a PA-1 guard using a Dionex ICS-6000 (ThermoFisher) and detected using pulsed amperometric detection (PAD). Flow was 1 mL/min and elution conditions were 0-25 min 5 mM NaOH and then 25-40 min, 5-100 mM NaOH. The software was Chromeleon Chromatography Data System. Monosaccharide standards were 0.1 mM, all data was obtained by diluting assays 1/10 before injection, apart from the ceramide samples, which were diluted 15 μL in 100 μL.

### Steady-state kinetics of AmPL38

Initial velocities of purified recombinant Amuc_0778^PL38^ were quantified on HA, CSA, CSC and DS substrates, with concentrations ranging from 0.25 to 12 g·l⁻¹ at 37 °C, 100 mM NaCl, and UB4 buffer at pH 7^48^. The average initial velocities, quantified in milli-absorbance units (mAU) at A235nm per second, were converted to µM of product generated by measuring the amount of Δ4,5 bonds formed per second, using the experimentally confirmed extinction coefficient for unsaturated glucuronic acid of 6150 M-1cm-1^43^. Kinetic parameters were determined by plotting initial velocities against substrate concentrations and fitting the Michaelis-Menten model using GraphPad Prism.

### Initial rates of AmGH105

The steady-state reactions of Amuc_0778^PL38^ were sealed to prevent evaporation and incubated overnight at 37°C. Subsequently, Amuc_0863^GH105^ was added to a final concentration of 1 µM, and the decrease in A235nm was monitored for a minimum of two hours at 37°C. The substrate concentration was calculated by converting the initial absorbance of the Amuc_0778^PL38^ steady-state reactions, minus absorbance backgrounds, to µM double bonds. The initial rates of Amuc_0863^GH105^ were then calculated as the loss of double bonds in µM per minute in absolute values.

### Liquid Chromatography-Mass Spectrometry (LC-MS) of GAG substrates

Duplicate time-course reactions for Amuc_0778^PL38^ were prepared under the same conditions as for the kinetics at 2 g·l⁻¹ substrate concentrations. Reactions were terminated by heating the samples at 95°C for 5 min. Amuc_0863^GH105^ was added to the 20-hour reaction to a final concentration of 1 µM and left to run for two hours before heat inactivation. The final sample preparation and LC-MS analysis were carried out using an Iontrap coupled with GlycanPac chromatography as previously described^49^. Compounds were observed as single or double charge m/z, primarily as deprotonated adducts. The compounds were identified by MS and MS^2^ fragmentation, if possible. Extracted ion chromatograms of identified compounds were prepared, and the areas of the peaks were used to quantify the products (Supplementary Figure 41). Fragments follow the nomenclature of Domon and Costello^50^.

### Glycan labelling

Released O-glycans were fluorescently labelled by reductive amination with procainamide as described previously using LudgerTagTM Procainamide Glycan Labelling Kit (LT-KPROC-96)^51^. Briefly, samples in 10 μL of pure water were incubated for 60 min at 65°C with procainamide labelling solution. Residual chemicals were removed from the procainamide labelled samples using LudgerClean S-cartridges (LC-S-A6). The purified procainamide labelled O-glycans were eluted with pure water (1 mL). The samples were dried by vacuum centrifugation and resuspended in pure water (50 μL) for further analysis.

### LC-FLR-ESI-MS

Procainamide labelled samples were analysed by LC-FLD-ESI-MS using an ACQUITY UPLC® BEH-Glycan 1.7 µm, 2.1 x 150 mm column at 40 °C on a Thermo Scientific UltiMate 3000 UPLC instrument with a fluorescence detector (λex = 310 nm, λem = 370) controlled by HyStar software version 3.2. Gradient conditions were: 0 to 10 min 15% A at a flow rate of 0.4 mL/min; 10 to 95 min, 15 to 43% A at a flow rate of 0.4 mL/min, 95 to 98 min, 43 to 90% A at 0.4 to 0.2 mL/ml; 98 to 99 min, 90% A at 0.2 mL/min; 99 to 100 min, 90 to 15% A at 0.2 mL/min; 100 to 103 min, 15% A at 0.2 mL/min; 103 to 115 min, 15% A from 0.2 to 0.4 mL/min. Solvent A was 50 mM ammonium formate pH 4.4 made from Ludger Stock Buffer (LS-N-BUFFX40); Solvent B was acetonitrile (Acetonitrile 190 far UV/gradient quality; Romil #H049). Samples were injected in 15% aqueous/85% acetonitrile; injection volume 20 µL. The UPLC system was coupled on-line to an AmaZon Speed ETD electrospray mass spectrometer (Bruker Daltonics, Bremen, Germany) with the following settings: source temperature 180°C, gas flow 10 L/min; Capillary voltage 4500 V; ICC target 200,000; maximum accumulation time 50 ms; rolling average 2; number of precursors ions selected 3, release after 0.2 min; Positive ion mode; Scan mode: enhanced resolution; mass range scanned, 400-1700. A glucose homopolymer ladder labelled with procainamide (Ludger; CPROC-GHP-30) was used as a system suitability standard as well as an external calibration standard for GU allocation. ESI-MS and MS/MS data analysis was performed using Bruker Compass DataAnalysis V4.1 software and GlycoWorkbench software^52^.

### Bioinformatics

Putative signal sequences were identified using SignalP 6.0^53^. Identities between different sequences were determined using Clustal Omega using full sequences^54^. The CAZy database (http://www.cazy.org) was used as the main reference for CAZymes^55^. Phylogenetic trees were completed in SeaView^56^ and final trees were produced in the Interactive Tree of Life (iTOL)^57^.

#### Pangenome analysis

To analyse the CAZymes across different species and strains, CAZyme sequences were downloaded from UNIPROT using the get.seq function from the Bio3d package in RStudio^58^. Sequence selection was based on the accession numbers of the CAZymes from the most up to date version of the CAZy repository at the time. Sequence txt files were converted to a FASTA format using the TabulartoFasta function from the following GitHub repository: https://github.com/lrjoshi/FastaTabular. All multiple sequence alignment and phylogenetic analysis was performed using Clustal Omega^54^. Percentage identity matrices were downloaded directly from the programme and merged with the original file from CAZy according to shared accession number. The presence or absence of different enzymes and how well conserved they were across strains and species was curated by hand.

#### Structural comparisons

Where possible, protein models were retrieved from the AlphaFold Protein Structure Database^59, 60^. Where the models had not been built, this was completed using AlphaFold2 Colab^61^. Previously published protein structures were retrieved from the RCSB Protein Data Bank. Different structures or models were superimposed using COOT^62^ and PyMOL was used to look at structures/models and generate figures. Glycan models were built using the Carbohydrate Builder on http://glycam.org.

## Acknowledgements

Many thanks to Professor William Willat’s group (Newcastle University, UK) for providing some of the substrates for us to test. Thank you to Carl Morland (Newcastle University, UK) for technical support. Many thanks to Professor Anne S. Meyer and Professor Willem van Schaik for their insightful comments about the manuscript. We would also like to thank Dr Stefan Janecek from the Slovak Academy of Sciences for generously sharing his insights regarding GH57 enzymes. The work was funded by The Academy of Medical Sciences (SBF0061175), the Wellcome Trust/Royal Society (224240/Z/21/Z), and T.O.Ojuri is funded by the BBSRC Midlands Integrative Biosciences Training Partnership (MIBTP) with his studentship in collaboration with industrial partners Ludger (Oxford, UK). Finally, thank you to fantastic parents/grandparents, Clare and Nick Crouch, who facilitated some of our experiments by babysitting late into the night.

## Data Availability

The full RNA-seq data are provided in Supplementary Data 1 and submitted to https://www.ebi.ac.uk/ena/browser/home with accession number PRJEB76658. The data that support the findings presented in this manuscript are available upon request from the corresponding authors.

## Supplementary Information

**Supplementary Table 1.** Summary of enzymes from *A. muciniphila* BAA-835 included in this paper. This is a summary of the enzymes explored in this report, how they were divided into different groups, their overall result, and where they are predicted to be localised in the cell according to Signal P 6.0 (grey shading highlights those possibly on the outside of the cell). The enzymes highlighted in pink are the ones that were highlighted by proteomics when *A. muciniphila* BAA-835 was grown on human breast milk (Kostopoulos *et al.* 2020).

|  | Enzyme | Headline | Predicted localisation | Importance judged by TnSeq (LogFC - 1.5 and lower) | Interaction with Mul1A | Highly conserved across different *Akkermansia* species | Cloning |
| --- | --- | --- | --- | --- | --- | --- | --- |
| Sialidases | Amuc_0625<sup>GH33</sup> | Broad-acting | 90 % SPI, 7 % SPII |  |  | ✓ | pET28b<br>Cl Kan |
|  | Amuc_1547<sup>GH181</sup> | α1,3 only | 51 % SPII, 48 % SPI |  |  | ✓ | pET28b<br>Cl Kan |
|  | Amuc_1835<sup>GH33</sup> | Broad-acting | 99 % SPI | ✓ |  |  | pET28b<br>Cl Kan |
| Fucosidases | Amuc_0010<sup>GH29</sup> | Specific for BGH type II structures | 99 % SPI |  |  |  | pET28b<br>GS Kan |
|  | Amuc_0146<sup>GH29</sup> | ★Puzzle★ | 71 % SPI, 27 % other | ✓ |  |  | pET28b<br>Cl Kan |
|  | Amuc_0186<sup>GH95</sup> | 2FL only | 99 % SPI |  |  |  | pET21b<br>Cl Amp |
|  | Amuc_0392<sup>GH29</sup> | Very broad for α1,3/4 | 99 % SPI |  |  | ✓ | pET28b<br>Cl Kan |
|  | Amuc_0846<sup>GH29</sup> | Very broad for α1,3/4 (apart from SFLNT) | 94 % SPI, 6 % SPII |  |  |  | pET28b<br>Cl Kan |
|  | Amuc_1120<sup>GH95</sup> | Broad α 1,2 – only one that can handle BGH I | 33 % SPI, 65 % other | ✓ | ✓ |  | pET28b<br>Cl Kan |
| α-linked exo activities | Amuc_1187<sup>GH27</sup> | Broad α-galactosidase | 99 % SPI |  |  |  | pET28b<br>Cl Kan |
|  | Amuc_1008<sup>GH31</sup> | Specificity for the core α-GalNAc | 99 % SPI |  |  |  | pET21b<br>Cl Amp |
|  | Amuc_1870<sup>GH31</sup> | ★Puzzle★ pNP-α-Glc | Other |  |  |  | pET21b<br>Cl Amp |
|  | Amuc_0216<sup>GH36</sup> | Broad α-GalNAc'ase. Only enzyme to degrade Tn and Forssman antigens. | 74 % SPII, 26 % SPI |  |  |  | pET28b<br>Cl Kan |
|  | Amuc_0517<sup>GH36</sup> | Specificity for BGA | 99 % SPI |  |  |  | pET28b<br>Cl Kan |
|  | Amuc_0855<sup>GH36</sup> | Specificity for Galα1,3Gal | Other |  |  |  | pET21b<br>Cl Amp |
|  | Amuc_1868<sup>GH57</sup> | ★Puzzle★ pNP-α-Glc | Other | ✓ |  | ✓ | pET28b<br>Kan |
|  | Amuc_1260<sup>GH63</sup> | ★Puzzle★ | 99 % SPI |  |  |  | pET28b<br>Cl Kan |
|  | Amuc_0060<sup>GH89</sup> | α-GlcNAc'ase but true substrate unclear | 99 % SPI | ✓ |  |  | pET28b<br>Kan |
|  | Amuc_1220<sup>GH89</sup> | GlcNAcα1,4-Gal disaccharide | 63 % SPI, 37 % SPII | ✓ | ✓ |  | pET28b<br>Cl Kan |
|  | Amuc_1420<sup>GH97</sup> | α-galactosidase, likely requires +2 subsite | 60 % SPI, 39 % SPII |  |  |  | pET28b<br>Cl Kan |
|  | Amuc_0017<sup>GH109</sup> | α-GalNAc'ase (not BGA or Tn) | TAT SPI |  |  | ✓ | pET28b<br>Cl Kan |
|  | Amuc_0920<sup>GH109</sup> | α-GalNAc'ase (not BGA or Tn) | TAT SPI |  |  |  | pET28b<br>Cl Kan |
|  | Amuc_0480<sup>GH110</sup> | α-galactosidase (not globotriose or P1 antigen) | 23 % SPI, 75 % other |  |  |  | pET28b<br>Cl Kan |
|  | Amuc_1463<sup>GH110</sup> | Broad α-galactosidase | 99 % SPI |  |  |  | pET28b<br>GS Kan |
| β-galactosidas | Amuc_0290<sup>GH2</sup> | Broad β-galactosidase | 99 % SPI | ✓ | ✓ | ✓ | pET21b<br>Cl Amp |
|  | Amuc_0539<sup>GH2</sup> | Galβ1,3Gal/GalNAc | 99 % SPI | ✓ | ✓ | ✓ | pET28b<br>Cl Kan |
|  | Amuc_0771<sup>GH35</sup> | Requires a Glc/GlcNAc in the +1 position for | 98 % SPI |  |  |  | pET21b<br>Cl Amp |
| | | β1,4-linked. Not β1,6 substrates. | | | | | |
| | Amuc_0824<sup>GH2</sup> | β1,3-galactosidase | 55 % SPII, 44 % SPI | | ✓ | ✓ | pET21b Cl Amp |
| | Amuc_1666<sup>GH2</sup> | β1,4-galactosidase with a Glc sugar in the +1 subsite | 99 % SPI | ✓ | | ✓ | pET28b Kan |
|  | Amuc_1667<sup>GH2</sup> | True substrate not identified, partial against GA2 | 99 % SPI | ✓ | ✓ |  | pET28b Kan |
| | Amuc_1686<sup>GH35</sup> | β1,3/6-galactosidase | 99 % SPI | | | ✓ | pET21b Cl Amp |
|  | Amuc_0497<sup>GH43</sup> | ★Puzzle★ | 99 % SPI | ✓ |  |  | pET28b Kan |
| | Amuc_0498<sup>GH43</sup> | ★Puzzle★ pNP-β-Gal | 94 % SPII, 6 % SPI | | | | pET29b Kan |
| β-HexNAc'ases | Amuc_0369<sup>GH20</sup> | Broad β-HexNAc'ase | 99 % SPI | | | | pET28b Kan |
| | Amuc_0397<sup>GH20</sup> | β1,3-GalNAc'ase, true substrate not identified | 99 % SPI | ✓ | | | pET28b Kan |
| | Amuc_0868<sup>GH20</sup> | Broad β-HexNAc'ase | 55 % SPI, 43 % SPII | | ✓ | | pET28b Kan |
| | Amuc_1032<sup>GH20</sup> | GlcNAcβ1,3Gal specificity, requires +2 sugar | 99 % SPI | | | ✓ | pET21b Cl Amp |
| | Amuc_1669<sup>GH20</sup> | β1,3-HexNAc preference (host substrates only) | 90 % SPI, 10 % SPII | ✓ | ✓ | | pET28b Kan |
| | Amuc_1815<sup>GH20</sup> | β1,3-GalNAc'ase, true substrate not identified | 61 % SPI, 39 % SPII | | | | pET28b Kan |
| | Amuc_1924<sup>GH20</sup> | β-GalNAc'ase, not P antigen | 99 % SPI | | | | pET21b Cl Amp |
| | Amuc_2018<sup>GH20</sup> | β1,3-HexNAc preference, not P antigen | 99 % SPI | | | | pET28b Kan |
| | Amuc_2019<sup>GH20</sup> | Broad β-HexNAc'ase | 69 % SPI, 29 % SPII | | | | pET21b Cl Amp |
| | Amuc_2136<sup>GH20</sup> | Broad β-HexNAc'ase, not GA2 | 99 % SPI | ✓ | | | pET21b Cl Amp |
| | Amuc_2148<sup>GH20</sup> | β1,3-GlcNAc'ase preference | 99 % SPI | ✓ | | | pET28b Kan |
| | Amuc_0052<sup>GH84</sup> | Broad β-GlcNAc'ase (host substrates only) | 99 % SPI | ✓ | | | pET21b Cl Amp |
| | Amuc_0803<sup>GH123</sup> | Broad β-GalNAc'ase | 99 % SPI | | ✓ | ✓ | pET21b Amp |
| | Amuc_2109<sup>GH3</sup> | β-GlcNAc'ase, true substrate not identified | Other | | | | pET28b Kan |
| GAGs | Amuc_0778<sup>PL38</sup> | HA, CSA, CSC, and DS | 99 % SPI |  |  |  | pET28b Kan |
|  | Amuc_0863<sup>GH105</sup> | Products of PL38 | 93 % SPI |  |  |  | pET28b Kan |
| Sulfatases | Amuc_0451<sup>Sulf20</sup> | 3'S-LNB and LacNAc | 51 % SPII, 48 % SPI |  | ✓ | Not tested | pET28b GS Kan |
|  | Amuc_0491<sup>Sulf20</sup> | 3'S-LNB | 99 % SPI | ✓ | ✓ | Not tested | pET29b GS Kan |
|  | Amuc_0565<sup>Sulf4</sup> |  | 99 % SPI |  |  | Not tested | pET28b GS Kan |
|  | Amuc_0864<sup>Sulf26</sup> |  | Other |  |  | Not tested | pET28b GS Kan |
|  | Amuc_0953<sup>Sulf20</sup> | 3'S-LNB and LacNAc | 99 % SPI |  |  | Not tested | pET28b GS Kan |
|  | Amuc_1033<sup>Sulf11</sup> | 6S-GlcNAc | 87 % SPI, 12 % SPII |  |  | Not tested | pET29b GS Kan |
|  | Amuc_1182<sup>Sulf19</sup> |  | 99 % SPI | ✓ |  | Not tested | pET28b GS Kan |
|  | Amuc_1290<sup>Sulf</sup> |  | Other | ✓ |  | Not tested | pET28b GS Kan |
|  | Amuc_1655<sup>Sulf16</sup> | 4S-Gal | 99 % SPI |  |  | Not tested | pET29b GS Kan |
|  | Amuc_1755<sup>Sulf16</sup> | 4S-Gal/GalNAc | 70 % SPII, 30 % SPI |  |  | Not tested | pET29b GS Kan |
|  | Amuc_1074<sup>Sulf11</sup> | 6S-GlcNAc | 99 % SPI |  |  | Not tested | pET29b GS Kan |
|  | Amuc_0121<sup>Sulf15</sup> | Partial 6S-Gal/GalNAc | 99 % SPI |  |  | Not tested | pET29b<br>GS Kan |
| ? | Amuc_1216<sup>GH177</sup> | ★Puzzle★ | TAT SPI |  |  | ✓ | pET28b<br>Cl Kan |
|  | Amuc_0623<sup>GHnc</sup> | ★Puzzle★ | 99 % SPI |  |  |  | pET28b<br>Cl Kan |
| Starch | Amuc_1621<sup>GH77</sup> | Ladder of products<br>from amylose | Other |  |  |  | pET29b<br>GS Kan |
|  | Amuc_1637<sup>GH13</sup> | ★Puzzle★ | Other |  |  |  | pET29b<br>GS Kan |
|  | Amuc_1751<sup>GH13</sup> | ★Puzzle★ | Other | ✓ |  |  | pET29b<br>GS Kan |
|  | Amuc_1812<sup>GH13</sup> | ★Puzzle★ | Other | ✓ |  |  | pET28b<br>Cl Kan |

GS – gene synthesis. Cl – cloned.

**Supplementary Table 2.**
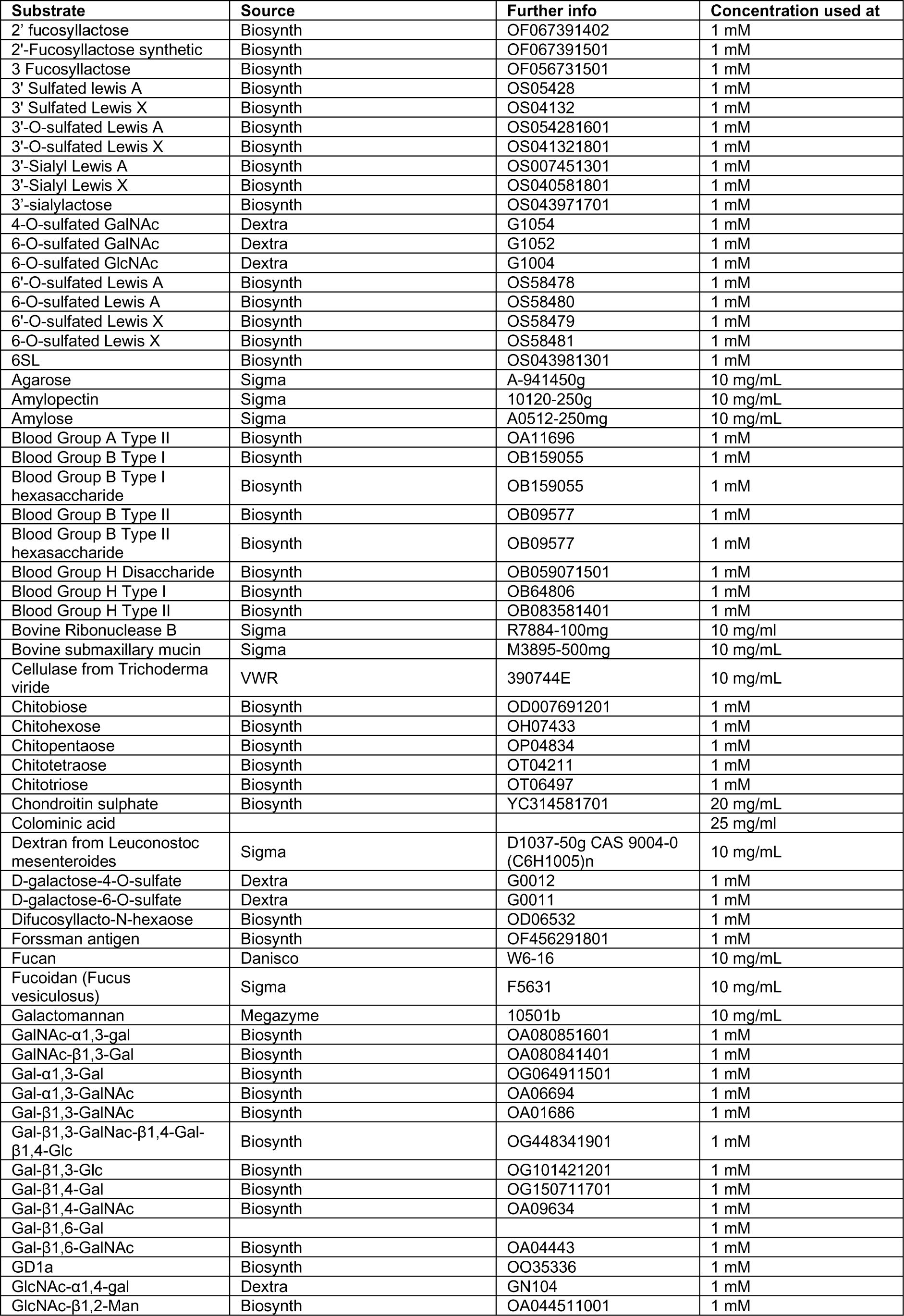
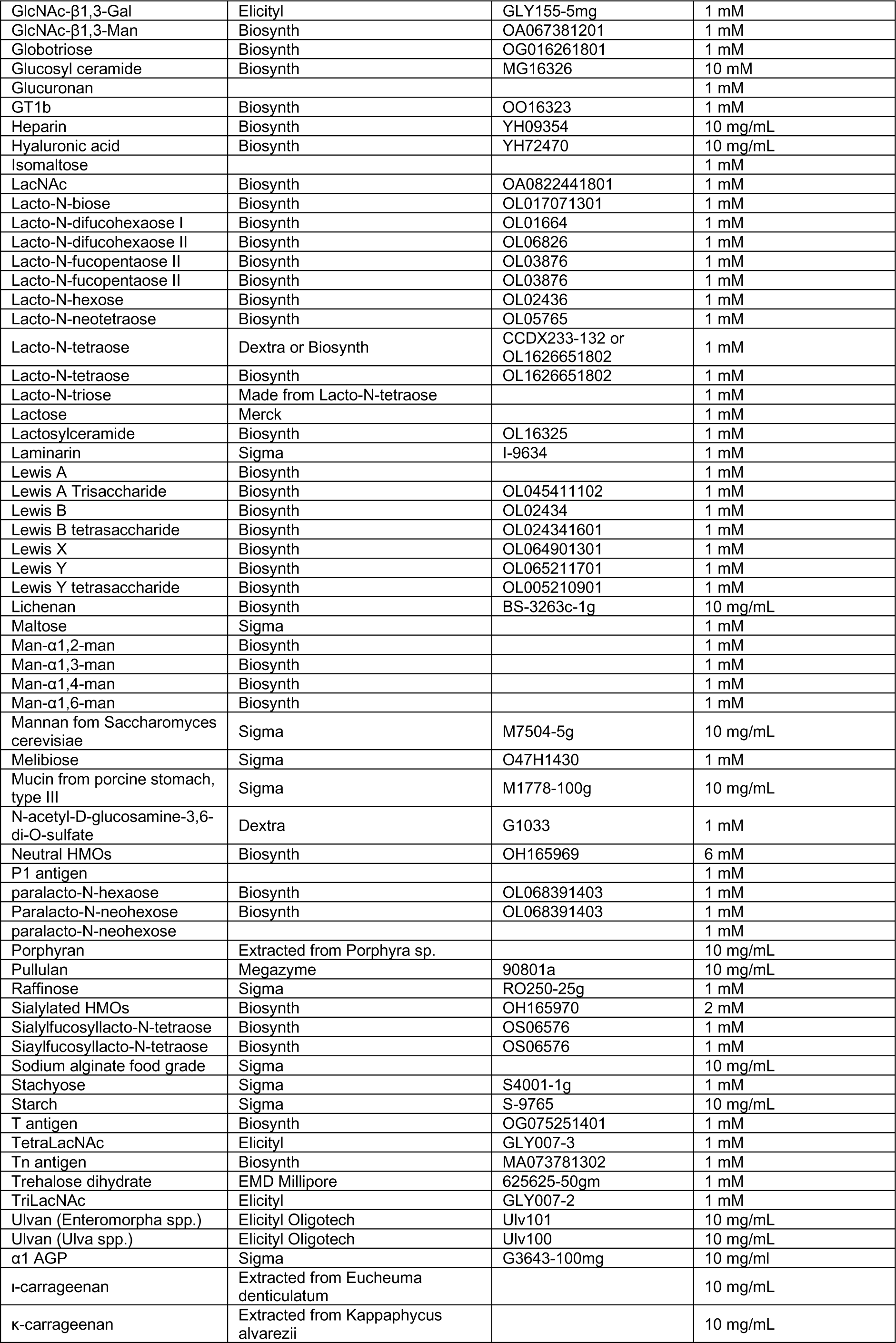
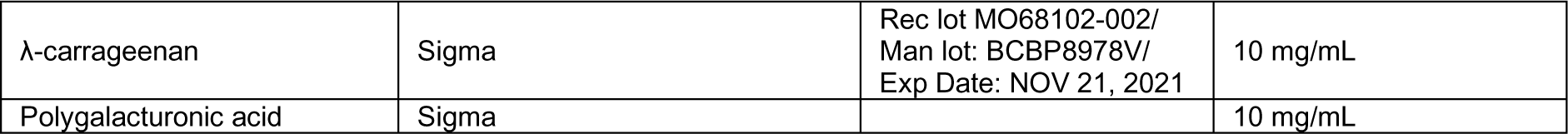
Substrates. Details about the substrates used in this work and the concentration they were used at.

**Supplementary Table 3.**
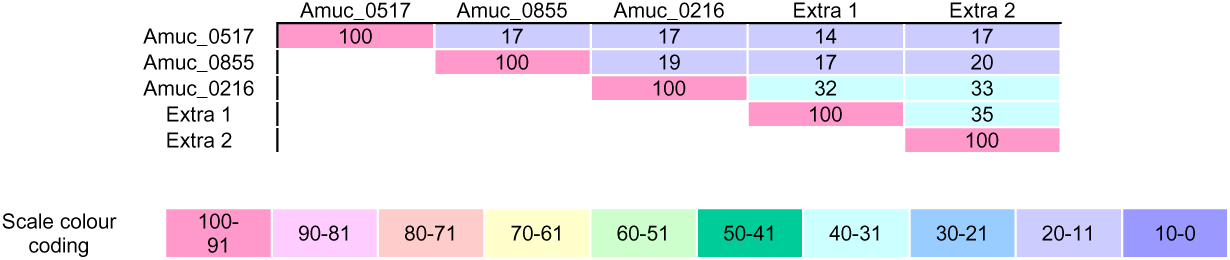
Prevalence of GH36 enzymes in *Akkermansia* strains. For this analysis, we took the GH36 sequences Extra 1 and 2 from *A. biwaensis*.

**Supplementary Table 4.**
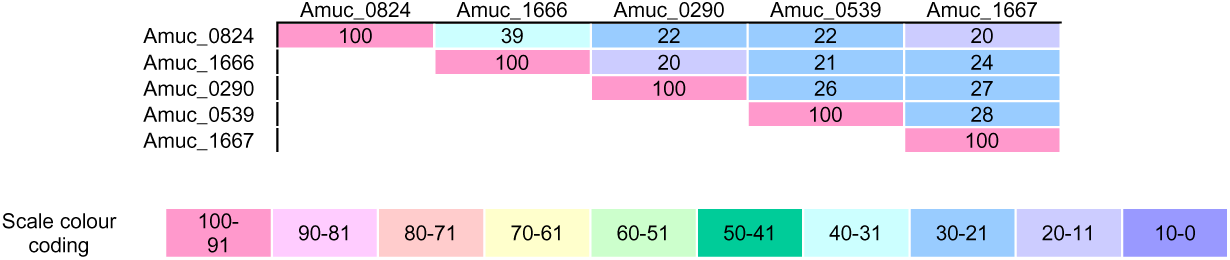
Percentage identity between GH2 family members from *A. muciniphila* BAA-835.

**Supplementary Table 5.**
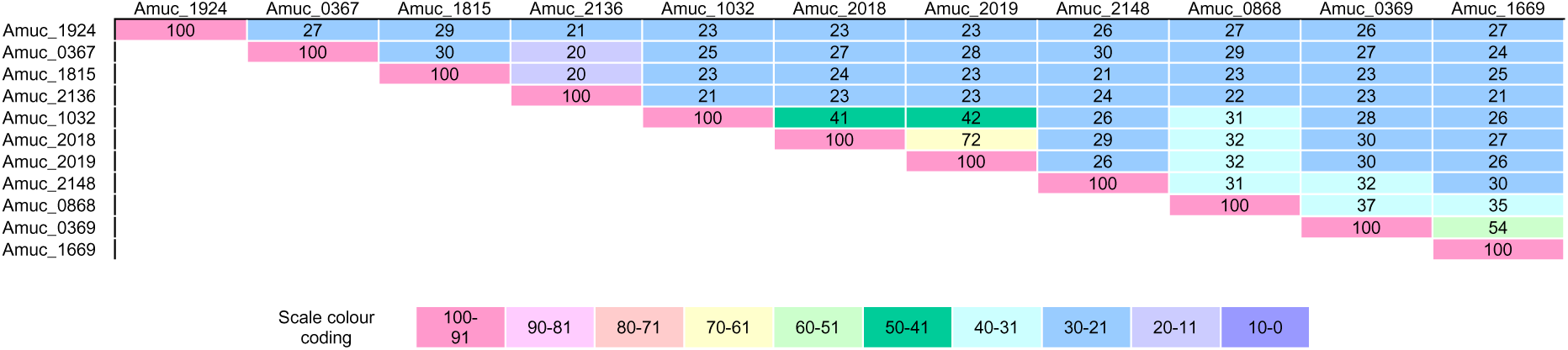
Percentage identity between GH20 family members from *A. muciniphila* BAA-835.

**Supplementary Table 6.**
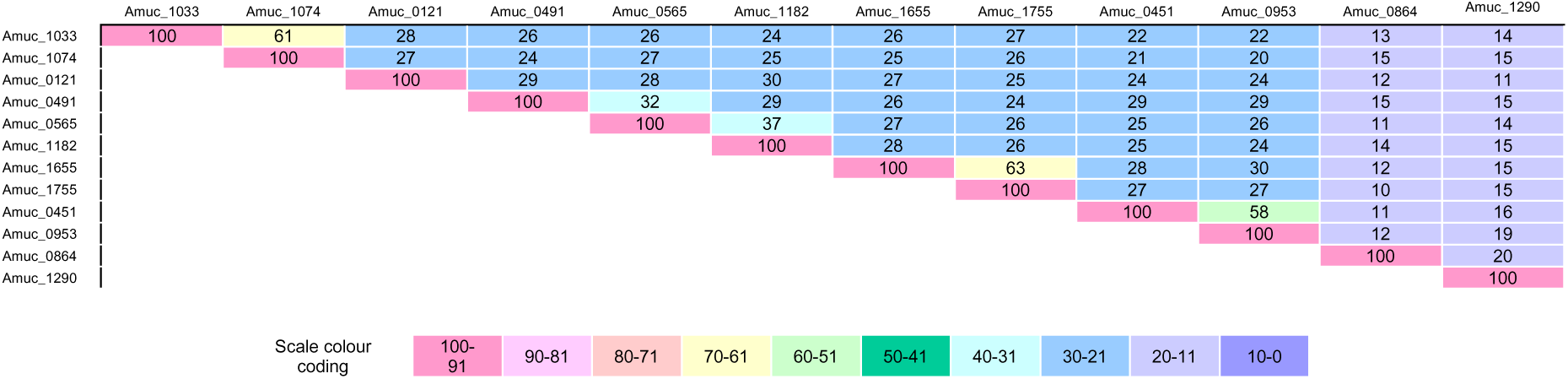
Percentage identity between putative sulfatases from *A. muciniphila* BAA-835.

**Supplementary Table 7.** Specific activities calculated from the highest rates observed in the kinetic analysis experiments of Amuc_0778^PL38^ and Amuc_0863^GH105^. One Unit was defined as μmol double bond produced/converted per minute. Initial substrate concentrations (µM double bonds) for Amuc_0863^GH105^ was calculated based on the absorbance (A235) of the samples after terminated overnight reactions of Amuc_0778^PL38^.

| Substrates | Amuc_0778<sup>PL38</sup> |  |  | Amuc_0863<sup>GH105</sup> |  |  |
| --- | --- | --- | --- | --- | --- | --- |
|  | Specific activity (U · mg<sup>-1</sup>) | Substrate concentration (g · l<sup>-1</sup>) | Relative activity (%) | Specific activity (U · mg<sup>-1</sup>) | Substrate concentration (uM) | Relative activity (%) |
| Hyaluronic acid | 29.373 ± 0.710 | 12 | 100 | 0.029 ± 0.002 | 1036 | 42 |
| Chondroitin sulfate A | 29.393 ± 0.401 | 12 | 100 | 0.069 ± 0.012 | 1016 | 100 |
| Chondroitin sulfate C | 4.426 ± 0.080 | 12 | 15 | 0.025 ± 0.002 | 1172 | 36 |
| Dermatan sulfate | 2.274 ± 0.258 | 8 | 8 | 0.026 ± 0.001 | 540 | 38 |
| Heparin | nd. | - | - | nd. | - | - |
| Polymannuronic acid | nd. | - | - | nd. | - | - |
| Polyguluronic acid | nd. | - | - | nd. | - | - |
| Polyglucuronic acid | nd. | - | - | nd. | - | - |
| Polygalacturonic acid | nd. | - | - | nd. | - | - |

**Supplementary Table 8.** Kinetic parameters for Amuc_0778^PL38^ on four different substrates. Parameters were obtained by fitting the Michaelis-Menten model to the initial rates for increasing substrate concentrations. ± represents standard deviations from triplicate experiments.

| | $k_{\text{cat}}$ (s<sup>-1</sup>) | $K_{\text{M}}$ (g · l<sup>-1</sup>) | $k_{\text{cat}} / K_{\text{M}}$ (s<sup>-1</sup> g · l<sup>-1</sup>) |
| --- | --- | --- | --- |
| Hyaluronic acid | 23.13 ± 0.81 | 2.98 ± 0.27 | 7.79 ± 0.45 |
| Chondroitin sulfate A | 29.24 ± 2.00 | 6.38 ± 0.93 | 4.62 ± 0.38 |
| Chondroitin sulfate C | 3.06 ± 0.11 | 3.06 ± 0.41 | 1.01 ± 0.11 |
| Dermatan sulfate | 0.77 ± 0.12 | 1.51 ± 0.50 | 0.53 ± 0.10 |

### RNA-seq

The transcriptional response of AM varied greatly when grown with the addition of mucin compared to glucose. A total of 201 genes (9.3%) were differentially expressed with 76 significantly upregulated (Log2 Fold Change > 1.5 & FDR < 0.05) and 125 significantly downregulated (Log2 Fold Change < -1.5 & FDR < 0.05). Of the 67 enzymes in this study, 22 were significantly upregulated and 0 were significantly downregulated.

### Pangenome analysis

One way to assess the importance of the AM CAZymes to the species in terms of mucin degradation is to review how prevalent and conserved the enzymes are over multiple *A. muciniphila* isolates (Supplementary Figure 3). In this analysis we also included *Akkermansia biwaensis* WON2089, twelve *Akkermansia massiliensis* strains, and five *Candidatus Akkermansia timonensis* strains. The latter two species have recently been reclassified as their own species but were previously cluster II and III of *A. muciniphila*^1^. The isolates included in this analysis were from a range of human faecal sources, including different collection efforts, geographical location, adults, and children.

The striking aspect of this analysis was how highly most of these enzymes were conserved between *A. muciniphila* strains (>90 %). There were a few examples of enzymes not being present in many strains (e.g. Amuc_0875^GH16^ and Amuc_0146^GH29^), but generally enzymes were present. As another generalisation, the other species assessed did have close homologues of the CAZymes in *A. muciniphila* BAA-835, with *A. massiliensis* having higher identity than *A. biwaensis* and *Candidatus A. timonensis*.

Additional enzymes were also identified during this analysis:

- A GH2 in some *A. muciniphila* strains.
- A GH2 in *A. biwaensis* and *Candidatus A. timonensis*.
- A GH16 in *A. biwaensis* and *Candidatus A. timonensis*.
- A GH27 in *A. massiliensis*.
- A GH27 in *A. biwaensis* and *Candidatus A. timonensis*.
- A GH29 in many of the *A. muciniphila* strains
- A GH29 in most *A. massiliensis*.
- A GH29 in *A. biwaensis* and *Candidatus A. timonensis*.
- A GH36 in *A. biwaensis* and *Candidatus A. timonensis*.
- A GH36 in *A. massiliensis, A. biwaensis* and *Candidatus A. timonensis*.
- A GH84 in most of all strains and species assessed.
- A GH95 in *A. massiliensis*.
- A GH95 in *A. biwaensis* and *Candidatus A. timonensis*.

### Further description of enzyme specificities and analysis of structural models

#### α-galactosidases

##### GH27

The characterised activities from GH27 family members so far include α-galactosidases, α-GalNAc’ases, β-L-arabinopyranosidases, and isomaltodextrinases. Bacterial GH27 family members with activity against α-Gal have been characterised in the context of plant polysaccharides, pectic galactan and galactomannan, for example, and there are no structures of these enzymes^2, 3^. Therefore, Amuc_1187^GH27^ is the first example of an α-galactosidase GH27 family member of prokaryotic origin with specificity for animal-type substrates rather than those of plant. The structural and mutational studies of two GH27 enzymes from humans has previously been insightful in terms of understanding their differing specificities for α-galactose or α-GalNAc^4^. In humans, two GH27 enzymes, one α-galactosidase and one α-GalNAc’ase (48 % sequence identity), are involved in the recycling of glycolipids and glycoproteins. Deficiencies in these enzymes in humans is linked to several lysosomal storage diseases due to the accumulation of glycoconjugates. In contrast to Amuc_1187^GH27^, the human enzymes can accommodate the fucose in the blood group structures to hydrolyse either α-Gal or α-GalNAc^5^. Between these two human enzymes, eleven out of thirteen amino acids coordinating the sugar in the -1 subsite were identical. Those that differed were key to the accommodation of either an α-Gal or α-GalNAc. In the α-GalNAc’ase, there were relatively small residues (serine and alanine) coordinating at the N-acetyl group of the GalNAc, but in the α-galactosidase these residues were larger (glutamate and leucine) thus prohibiting the binding of GalNAc (Supplementary Figure 15). Mutation of the key residues in both enzymes to that of the other, allowed the swapping of specificities^4^. Amuc_1187^GH27^ has approximately 30 % sequence identity to the human enzymes, but in terms of the active site, the residues coordinating the -1 sugar in the two human enzymes are generally conserved in Amuc_1187^GH27^ and it has a serine and asparagine at the critical locations (S320 and N321), aligning with the observed activity of α-galactosidase (Supplementary Figure 15).

##### GH97

Amuc_1420^GH97^ was able to hydrolyse galactose from PGMIII from samples with or without sialidase and fucosidase pre-treatment, comparable to that of the other AM α-galactosidases (Figure 1 and Supplementary Figure 18). It was also able to hydrolyse α-galactose capping substrates (not blood groups) and had a preference for the trisaccharide globotriose. Intrigued by this contrast in activities, we analysed an Alphafold model of Amuc_1420^GH97^ alongside available GH97 structures. The only GH97 α-galactosidase crystal structure with a ligand is that of BT1871 from *Bacteroides thetaiotaomicron*, which has been crystallised with a lactose that also had an α-linked galactose to the Glc through a 1,1-linkage^6^. This ligand was overlaid into the Alphafold model for Amuc_1420^GH97^ and suggests there would be significant interactions with the sugar in the +2 subsite through a tryptophan (W461) (Supplementary Figure 17). This likely explains why poorer activity is detected against disaccharide host-type substrates compared to trisaccharides and complex substrates.

##### GH36

There are three putative GH36 enzymes encoded in the AM genome, and these have relatively low sequence identity between them (Supplementary Table 3). The family members characterised so far display exo-acting activities towards α-Gal or α-GalNAc, with attention predominantly on the former in terms of raffinose-type oligosaccharides and galactomannan. Amuc_0855^GH36^ was able to liberate galactose from Galili antigen, globotriose, and PGMIII with or without sialidase and fucosidase treatment (Figure 2 and Supplementary Figure 18). Galili antigen is expressed by most mammals and some other animals, but not humans, apes, and Old World monkeys^7^. Therefore, gut microbes would only have access to this structure through animal products in the diet of humans. On the phylogenetic tree, this enzyme clusters with α-galactosidases characterised to be active against both galactomannan and raffinose-type oligosaccharides (Supplementary Figure 16). Amuc_0855^GH36^ also had trace activity against these substrates, but no growth was observed on these substrates for AM, so this is unlikely to be physiologically relevant (Supplementary Figure 1).

The whole cell assays showed no activity against the defined substrates with α-galactose at the non-reducing end, suggesting that none of the AM CAZymes acting on these epitopes are present on the outside of the cell (Supplementary Figures 4&5).

#### α-GalNAc’ases

##### GH31

A model of Amuc_1008^GH31^ was compared to the structure of the *E. faecalis* enzyme, which is the only structure from this subfamily. An overlay indicates that the structure of Amuc_1008^GH31^ has extra C-terminal domains compared to *Ef*GH31 (Supplementary Figure 14). One of these extra domains is classified as a family 32 carbohydrate binding domain (CBM32), which have been to show a variety of different sugars and glycans. The GH31 from *Clostridium perfringens* with the same activity as Amuc_1008^GH31^ was also has a CBM32, which has been characterised to bind GalNAc^8^. Tn antigen that was crystallised in the active site of *Ef*GH31 overlays well into the active site of Amuc_1008^GH31^ (Supplementary Figure 39). The residues interacting with the Tn-antigen in the *Ef*GH31 crystal structure are all conserved in Amuc_1008^GH31^.

#### α-GlcNAc’ases

There are currently five characterised GH89 family members all with specificity for this α-linked GlcNAc^9^. The first enzyme identified to have specificity for stomach epitope was one from *C. perfringens*^10^ and was subsequently crystallised with this disaccharide in the active site (4A4A; Supplementary Figure 22)^11^. Other members of the GH89 family are active against heparan sulfate and there is a crystal structure of this from *H. sapien* (4XWH)^12^. Mutations of this GH89 gene leads to Sanfilippo syndrome or mucopolysaccharoidosis III, where recycling of heparan sulfate is impaired, and there is currently no cure for this or treatments to slow disease progression. Detailed characterisations of glycobiology (or absence of) leading to disease may highlight potential ideas for treatments in the future.

Models of the two GH89 enzymes from AM and these two crystal structures were superimposed and compared (Supplementary Figure 19). The active site of Amuc_1220^GH89^ has a similar conformation to the pocket from *C. perfringens*, whereas the substrate pocket of Amuc_0060^GH89^ has a similar conformation to the pocket from *H. sapien* (Supplementary Figure 22). This implies that the substrates for Amuc_1220^GH89^ and Amuc_0060^GH89^ are GlcNAcα1,4-Gal stomach epitope and heparan sulfate or other GlcNAc-containing glycosaminoglycans, respectively. The data obtained in this report show that Amuc_1220^GH89^ is removing GlcNAc from PGMII, but we did not observe any activity for Amuc_0060^GH89^ against GAGs incubated alone or in combination with other AM CAZymes (See GAGs section below).

Amuc_1220^GH89^ is predicted to be localised to the periplasm (Supplementary Table 1) and whole cell assays using GlcNAcα1,4-Gal suggest that this epitope is not broken down on the outside of the cell (Supplementary Figure 4&5).

#### α-sialidases

The two GH33 family members (Amuc_0625^GH33^ and Amuc_1835^GH33^) are broad-acting sialidases in terms of linkage, whereas the Amuc_1547^GH181^ is α2,3-specific^13^. In this work, we explored whether these enzymes act upon ganglioside structures, which have structural similarities to O-glycans (Supplementary Figure 20). Gangliosides are important components of eukaryotic plasma membranes and are linked to several genetic diseases and pathogen interactions^14^. Gangliosides (GM3 and GD3; Figure 1) are also found in the membranes of milk fat globules in breast milk, so it is likely that gut microbes in the infant gut would encounter these^15^. Therefore, gangliosides would be found in the large intestine at all life stages from both host and dietary sources. We found that the two GH33 enzymes from AM can act on the relatively complex GD1a and GT1b ganglioside structures, with one or two sialic acids being removed. The inability to remove one of the sialic acids is likely due to this being the branching sialic acid and therefore relatively difficult to access. However, if we removed the capping β1,3-linked galactose with β-galactosidases from AM, then it was possible for Amuc_1835^GH33^ to remove all sialic acids (Supplementary Figure 20). Importantly, the removal of this galactose produces Sda antigen, which is present in 91 % of humans and is expressed in mucosal surfaces and other tissues. Therefore, is it valuable to understand that sialidases from AM can act on Sda antigen and gangliosides. Amuc_1835^GH33^ was recently highlighted as important for growth on mucin using a Tn-seq mutant library^16^, and Amuc_0625^GH33^ and Amuc_1547^GH181^ were highly conserved (>90 % sequence identity) across all *Akkermansia* species assessed (Supplementary Figure 5).

#### α-fucosidases

Between them, the GH29 and GH95 enzymes from AM can degrade a wide variety of fucose-rich substrates (Supplementary Figure 21&22)^13, 17^. These enzymes cannot access the fucose in blood group A and B (BGA and B) structures, so these glycan epitopes are first partially broken down by other CAZymes encoded in the AM genome to produce blood group H structures. Amuc_1120^GH95^ is a broad-acting α1,2-fucosidase, whereas Amuc_0010^GH29^ had specificity to α1,2-linked fucose on lactose/LacNAc (β1,4), but not Lacto-N-biose (β1,3). This means that Amuc_0010^GH29^ exhibits specificity towards type II blood group structures (not type I) and the lacto-N-*neo*tetraose HMO series (not the lacto-N-tetraose). In contrast, Amuc_0846^GH29^ and Amuc_0392^GH29^ have comparable activities with activities against α1,3 and 4-linked fucose, but Amuc_0392^GH29^ did exhibit some slightly broader activity by also being able to liberate galactose from some sulfated Lewis structures and sialylfucosyllacto-N-tetraose. For Amuc_0186^GH95^, we were only able to find activity against 2’-fucosyllactose and for Amuc_0146^GH29^ we could find no activities, including with pNP-monosaccharides.

Amuc_0146^GH29^ and Amuc_1120^GH95^ were recently highlighted as important for growth on mucin using a TnSeq mutant library^16^ and Amuc_0392^GH29^ is highly conserved across different *Akkermansia* species (Supplementary Figure 3). Furthermore, five out of nine of the sialidases and fucosidases were highlighted by proteomics when AM was grown on human milk (Supplementary Table 1)^18^.

#### β-galactosidases

The five GH2 enzymes show low sequence identity, with the highest being 39 % between Amuc_0824 and Amuc_1666 (Supplementary Table 4). The two GH35 enzymes share 37 % identity and the two GH43_24 enzymes have 87 % identity. Amuc_0539^GH2^ was found to be specific to β1,4-linked Gal with either a Gal or GalNAc in the +1 position, with some activity against lactose (Supplementary Figures 24-26). Conversely, Amuc_0771^GH35^ and Amuc_1666^GH2^ could not hydrolyse these substrates and instead required a Glc/GlcNAc in the +1 position for β1,4-linked Gal substrates. Amuc_1666^GH2^ was also able to act on TriLacNAc and Lacto-N-neotetraose, but only Galβ1,3Glc out of those substrates with β1,3-linked Gal. Amuc_0771^GH35^ was much broader with the capacity to also hydrolyse β1,3-linked Gal with Glc, GalNAc, and Gal in the +1 position. This is also the only enzyme that could remove both Gal sugars from the branching lacto-N-neohexaose. These branching structures, formed through a GlcNAcβ1,6Gal linkage, are important due to their prevalence in humans (99 % of population) and are referred to as “I antigens” (no branching is referred to as “i antigen”). Mucins across most tissues will have this branching^14^. Complementary to the screening carried out in this report, Amuc_1666^GH2^ has previously been shown to have a preferences for LacNAc^19^ and Amuc_0771^GH35^ has also previously been shown to have activity against mucin core 1 and core 2 structures, which both have β1,3 linkages^20^.

Amuc_0824^GH2^ is highly specific towards β1,3-linked Gal with no preference observed for what occupies the +1 positions and no activity against β1,4-linked Gal substrates. In line with our findings, Amuc_0824^GH2^ has previously been shown to have preferences for Galacto-N-biose^19^. Amuc_1686^GH35^ showed very similar specificity to Amuc_0824^GH2^ except that it could also accommodate β1,6-linked Gal with Gal/GalNAc in the +1 (no β1,6-linked substrates with a Glc-type sugar were available to test) and has previously been shown to have a preference for Galβ1,3GalNAc^21^. Amuc_0290^GH2^ can accommodate β1,4 and β1,6-linked Gal substrates with less preference for β1,3-linked Gal substrates. Finally, Amuc_1667^GH2^ was inactive against most substrates, with only some activity towards the ganglioside core structure GA1. Concerning biantennary complex N-glycans, all three enzymes that can accommodate other Galβ1,4GlcNAc structures, could remove Gal from N-glycan that had been released from the protein and de-sialylated. This is unlikely to be physiologically relevant, however, as AM will not grow on this glycoprotein substrate despite similarities to O-glycans (Supplementary Figure 2).

Regarding larger more complex substrates, Amuc_0771^GH35^ was able to release the largest concentration of galactose from BSM (core 1; Supplementary Figure 9). Amuc_0824^GH2^ was also able to release galactose, which complements the understanding that this substrate constitutes core 1 structures, but Amuc_0771^GH35^ can release a higher concentration of galactose indicating that it can accommodate the mucin polypeptide more easily than Amuc_0824^GH2^.

Of these enzymes, Amuc_0824^GH2^ is predicted to localise to the cell surface, while the rest are predicted to reside in the periplasm (Supplementary Table 1). Notably, the mucin-grown whole cells showed rapid degradation of Lacto-N-biose, one of the substrates for this enzyme, and this was reproducible over two different experiments (Supplementary Figure 4&5). This was the fastest degradation of a substrate, with complete hydrolysis recorded in 30 minutes. Lactose and Galβ1,4GalNAc are not substrates for this enzymes and the whole cell assays did not show degradation of these substrates (Supplementary Figure 5). Amuc_0824^GH2^ cannot be directly linked to this activity, but techniques that have been developed for the genetic manipulation of AM may lend themselves to these types of experiments in the future.

Of the fourteen enzymes identified as having >90 % sequence identity conservation across the different *Akkermansia* species we assessed, all the GH2 and GH35 family members, apart from Amuc_0771^GH35^ and Amuc_1667^GH2^, were in this group (Supplementary Figure 3) and Amuc_0290^GH2^, Amuc_0539^GH2^, Amuc_1666^GH2^ and Amuc_1667^GH2^ were highlighted as important for growth on mucin^16^. This emphasises the important role these β-galactosidases play in nutrient acquisition for AM (Supplementary Tabel 1). No activities for the GH43_24 enzymes were observed against the substrates tested here, but Amuc_0698^GH43^ was active against pNP-Gal (Supplementary Figure 27). Interestingly, Amuc_0697^GH43^ was also highlighted as important for growth on mucin^16^, conversely however, Amuc_0697^GH43^ was not well conserved between *A. muciniphila* strains (Supplementary Figure 3).

One of the striking aspects of CAZymes from AM is the length of the putative enzymes, which structural modelling predicts to be a high number of accessory modules accompanying the catalytic module. This is exemplified by the β-galactosidases (Supplementary Figure 29). How these accessory modules contribute to function is still yet to be determined. The active sites of the different GH2 and GH35 enzyme models were compared to relevant existing structures to provide insight into the different specificities being observed. Amuc_0539^GH2^ has specificity to β1,4-linked Gal with either a Gal or GalNAc in the +1 position, not Glc/GlcNAc. Galactose and glucose type sugars have the hydroxyl in the axial and equatorial positions at C4, respectively. The difference means that β1,4-linked sugars have very different overall conformations, and this is reflected in some of the specificities observed for the β-galactosidases. Models of different disaccharides are included to contextualise these possible conformational differences (Supplementary Figure 30). Current structures do not include any examples with substrates with Gal/GalNAc in the +1 subsite, but LacNAc and lactose examples are available. When we overlay LacNAc from *Streptococcus pneumoniae* GH2 (4CUC) into the active site of Amuc_0539^GH2^ a tryptophan is highlighted as being very close to the +1 GlcNAc. This suggests that this amino acid may contribute significantly to excluding substrates with Glc-type sugars in the +1 subsite. Similarly, Amuc_0290^GH2^, which has a much broader activity, has W686 in the same region, but rotated approximately 180 ° along its plane. This may act to contribute to binding glycans rather than having a selectivity role. Apart from these observations, the active sites of these two enzymes are predicted to be generally quite open.

In contrast to the GH2 and GH35 enzymes, the GH43 proteins only constitute a catalytic module and they are very similar in sequence. The only difference is a short section of sequence that corresponds to an extended loop in Amuc_0697^GH43^ that protrudes towards the active site (Supplementary Figure 27).

#### β-HexNAc’ases

The identity between the GH20 sequences from AM is generally low ∼20-30 % with the exceptions of 72 % between Amuc_2018^GH20^ and Amuc_2019^GH20^. These enzyme sequences were put into phylogenetic trees alongside other characterised members of their families (Supplementary Figures 31). These showed that the enzyme sequences from the AM genome clustered with previously characterised enzymes with β-HexNAc’ase activities. The first step in this analysis was to screen a panel of defined oligosaccharides (Supplementary Figures 33-35). The enzyme with the broadest activity was Amuc_0868^GH20^ that could degrade most of the substrates provided, apart from GlcNAcβ1,3Man and β1,2-linked GlcNAc on biantennary N-glycans, to which it was only partially active. This enzyme was previously shown to prefer GlcNAc over GalNAc^22^. Amuc_2019^GH20^ showed a similar pattern but exhibited only partial activity against the P antigen. Again, Amuc_2136^GH20^ showed a similar pattern to Amuc_0868^GH20^, but with no activity against the β1,4-linked GalNAc on GA2. Amuc_2136^GH20^ has also previously been shown to prefer GlcNAc over GalNAc^23^. Amuc_0369^GH20^ was also relatively broad-acting, but with a different pattern; it was able to access most of the host-type glycans tested and demonstrated relatively less activity against chitooligosaccharides. Amuc_2018^GH20^ showed a similar pattern to these three broad-acting GH20 enzymes, but with negligible activity against chitooligosaccharides and no activity against β1,4-linked GalNAc and P antigen. Amuc_2018^GH20^ has previously been shown to prefer GlcNAc over GalNAc and have some activity against N-glycan structures^22^. Amuc_0052^GH84^ showed specificity towards GlcNAc over GalNAc substrates and prefers smaller substrates (disaccharides). From the substrate screen, Amuc_1669^GH20^ is selective for host glycans with a preference for GlcNAcβ1,3Gal substrates. It can also hydrolyse GalNAc in the context of globoside P antigen, but not ganglioside GA2. Amuc_2148^GH20^ shows very similar pattern of activity to Amuc_1669^GH20^ but is even more specific as it will not act on P antigen. Amuc_1924^GH20^ has a preference for GalNAc in the -1 subsite but was unable to accommodate P antigen and this also may be due to overall glycan conformation. Amuc_1032^GH20^, acted only upon GlcNAcβ1,3Gal in the context of TriLacNAc and lacto-N-triose, but not GlcNAcβ1,3Gal alone, indicating that this enzyme prefers substrates that are at least three sugars long (i.e. a sugar occupying the +2 subsite). Finally, Amuc_0803^GH123^ had strict specificity towards the GalNAc substrates, including GA2. Exact specificities could not be determined for Amuc_0397^GH20^ and Amuc_1815^GH20^. Both only had trace activity against GalNAcβ1,3Gal, but it may be that they are active on longer versions of this substrate. Specificities could also not be found for Amuc_2109^GH3^, with only trace activity against chitooligosaccharides and GaNcAcβ1,3Gal.

In terms of the redundancy seen for the β-galactosidases and β-HexNAc’ases, the expression and localisation of these different enzymes in AM may require this redundancy in specificity for such a complicated substrate to maximise AMs access to the sugars in mucin. The conservation of most of the β-galactosidases and β-HexNAc’ases over species and strains supports this hypothesis.

### Activity of AM CAZymes against human milk oligosaccharides

The growth of AM on human milk has previously been demonstrated and proteomics highlighted a high proportion of the AM CAZymes ^18^(Supplementary Table 1). The structures of human milk oligosaccharides have many similarities to mucin O-glycans, so we performed sequential reactions on two commercially available products. We first tested the AM CAZymes that remove the different α-linked capping monosaccharides (Supplementary Figures 36&37). Interestingly, we found no release of sialic acid, despite one of the substrates being labelled as such. In terms of fucosidases, most of the AM enzymes could act on one of the products. Discrepancies were observed between the activities of Amuc_0010^GH29^ and Amuc_1120^GH95^ against HMOs, with the latter releasing more fucose and, therefore, these HMOs have predominantly lacto-N-tetraose structures. This observation corresponds to the understanding of type II structures being present on red blood cells and type I structures are associated with glycan decorations on secretions^14^. Notably, Amuc_0010^GH29^ clusters on a phylogenetic tree with ‘Mfuc5’ from a soil metagenome and has comparable activities^24^. For the AM CAZymes with specificity for removing α-linked GlcNAc, GalNAc, or galactose, no obvious activity was observed.

The panel of β-galactosidases were then tested against the HMOs which had also been pre-treated with two fucosidases (Supplementary Figure 37). The addition of Amuc_0290^GH2^, which had the broadest specificity in the substrate screen, had the most activity (shifting all the bands up the most). Much less galactose could be released without the addition of fucosidases (Supplementary Figure 26). Finally, the panel of β-HexNAc’ases was then tested against the HMOs that had been sequentially treated with fucosidases and galactosidases. The specificities observed for the defined substrate screen facilitates the probing of more complicated structures in more detail and exemplifies how these enzymes may be useful in other contexts. For example, Amuc_0803^GH123^ showed no obvious activity towards the HMO substrates, implying that no β-linked GalNAc is present. Furthermore, Amuc_1032^GH20^ is providing the same pattern as the broad-acting β-HexNAc’ases, so the glycans in this assay were at least 3 sugars long. Finally, Amuc_0052^GH84^ did not breakdown the glycans as much as the broad-acting β-HexNAc’ases, emphasizing its preference for smaller substrates.

### Interaction with GAGs

The PL38 enzymes characterised to date have specificities for glucuronan and alginate^25, 26^, but screens of these substrates did not reveal any activity (Supplementary Figure 42). Characterised GH105 enzymes have been found to be active against the unsaturated products of PLs ^27^. We found no activities for GH105 against any of the other substrates used in this work. Therefore, we tested host GAGs that are known substrates for some PL families and are commonly available to the microbiota in the large intestine. Amuc_0778^PL38^ was found to be active against hyaluronic acid (HA), chondroitin sulfate A (CSA), chondroitin sulfate C (CSC) and dermatan sulfate (DS) (Supplementary Figure 42-44). Amuc_0863^GH105^ had little or no activity when it was incubated alone with these substrates but was able to act on the products of the PL38 from all four substrates.

The Michaelis-Menten kinetic analysis of Amuc_0778^PL38^ against the different GAGs revealed the highest catalytic efficiency (*k*_cat_/*K*_M_) towards HA, followed closely by CSA and four- and eight-fold lower towards CSC and DS, respectively (Supplementary Table 7, Supplementary Figure 43). An attempt to resolve the kinetic parameters of Amuc_0863^GH105^ was performed by preparing substrate using the Amuc_0778^PL38^ and relevant GAGs. Amuc_0863^GH105^ was then added and the loss of the double bond of the unsaturated uronic acid was monitored at

235 nm. While linear decreasing initial rates were observed for the lower substrate concentrations, the higher concentration reactions were not reliable because of the absorbance ceiling of the spectrophotometer. Attempts to downscale or dilute the reactions did not provide data for which a reliable kinetic model could be fitted (Supplementary Figure 43). However, by isolating the highest observed initial rate for each substrate, a clear selectivity of Amuc_0863^GH105^ was observed towards CSA, followed by the three other GAGs at 36-40% relative activity (Supplementary Table 6). These results indicates that this enzyme prefers an unsaturated GlcA in the -1 subsite and a 4SGalNAc in the +1 subsite.

The products of Amuc_0778^PL38^ were then characterised in more detail using LC-MS/MS with UV detection (Supplementary Figure 42). Time-course reactions were performed to assess the product formations. For HA, Amuc_0778^PL38^ produced dimers, tetramers and hexamers. For CSA, the sulfated disaccharide and double sulfated tetrasaccharide were produced at equal rates initially, but as the disaccharide continued to increase in concentration, the tetrasaccharide decreased, indicating degradation to disaccharides also. A different pattern was observed for CSC, where the activity was slower and the tetrasaccharide was the dominant product initially, indicating that the differing sulfation pattern in this substrate reduces activity. The activity was even slower for DS and the double sulfated tetrasaccharide was the dominant product until much later in the time-course. The only difference between CSA and DS is the presence of GlcA and IdoA, respectively, therefore based on this data, Amuc_0778^PL38^ appears to prefer GlcA over IdoA containing substrates.

Finally, we included the sulfatases with the Amuc_0778^PL38^ (with and without Amuc_0863^GH105^) in assays with CSC and HS and no obvious activity could be seen by TLC (Supplementary Figure 45). The two GH89 enzymes were also tested against GAGs both with and without Amuc_0778^PL38^ and Amuc_0863^GH105^, but no activity was observed (Supplementary Figure 43).

Why have CAZymes if not to use them for nutrient acquisition? Our current hypothesis is that these enzymes give AM a colonisation advantage. GAGs are an important part of the glycocalyx and would be constant presence in the lumen of the colon. Another prominent member of the human gut microbiota, *Bacteroides thetaiotaomicron*, prioritises GAGs as a nutrient source, which demonstrates its significance in the colonic environment^28^. In AM, this enzyme may be one way that AM can burrow into the mucosal layer. Interestingly, a hyaluronidase from *Streptococcus agalactiae* has been shown to dampen down the immune system and increase invasion in the context of the female reproductive system and pre-term labour^29^. This system may be comparable to the role of the GAG-active CAZymes in AM.

### Growth of AM on high-mannose N-glycoproteins

When testing the growth of AM against a variety of substrates to support the biochemical work, we included glycoproteins that are decorated with mammalian complex N-glycans, as the sialidases, galactosidases, and GlcNAc’ases were observed to breakdown these substrates. The detected enzyme activities were unsurprising, as mammalian complex N-glycans have very similar glycan structures to O-glycans. For completeness, a glycoprotein with high-mannose N-glycans was also included. Unexpectedly, AM was observed to grow on the high-mannose N-glycoprotein and not the two glycoproteins with complex N-glycans (Supplementary Figure 1). Furthermore, we also observed growth on *Saccharomyces cerevisiae* mannan (*Sc*mannan), which is an N-glycan core structure with large antennae composed of α-linked mannose and a recognised nutrient source for other gut microbes^30^.

Whole cell assays using cells that had been grown on high-mannose N-glycoprotein revealed a smear on the TLC that appeared overnight (Supplementary Figure 46). This smear did not run as fast as the free high-mannose N-glycan control (suggesting a higher molecular weight), but when the sample was treated with PNGaseL, its TLC migration pattern was the same. This strongly suggests that AM is using the protein as a nutrient source and the by-product of this is high-mannose N-glycan that is still attached to a small peptide. The molecular mechanism underpinning why AM can degrade this substrate (and not other proteins or glycoproteins included in this work) is beyond the scope of this study, but it is likely attributable to a particular glycopeptidase or peptidase. However, this observation indicates a possible important cross feeding relationship AM may have with other human gut microbes. N-glycans are prevalent decorations on secreted proteins and high-mannose N-glycans are universal to eukaryotes, so microbes in the human gut would encounter this substrate from the host (secreted proteins and sloughed off dead cells) and any type of diet (animal- or plant-derived).

The whole cell assays for *Sc*mannan did not show any detectable glycobiology by TLC. All the CAZymes associated with the hydrolysis of α-linkages were tested against a panel of α-mannose glycans during the process of deciphering this growth, but there were no positive results (Supplementary Figure 13). SDS-PAGE with Coomassie staining of *Sc*mannan also showed no obvious protein in the sample (Supplementary Figure 46). To rule out very small free peptides being used as a nutrient source, the *Sc*mannan was dialysed to try and remove this, but growth was comparable to the original substrate (Supplementary Figure 1). It may be that there is a small amount of peptide attached to the *Sc*mannan that AM can access.

### Starch-degrading activities

Encoded in the AM genome are two GH families that are currently characterised to have sole specificity towards starch-type polysaccharides – one GH77 and three GH13 enzymes. These enzymes were tested against several different substrates composed of α-linked glucose, however activity was only found for Amuc_1621^GH77^, primarily against amylose and produced oligosaccharides of one to five glucose units (Supplementary Figure 47). There was no activity against dextran or amylopectin, but potentially trace activity against pullulan. There was no growth on these substrates (Supplementary Figure 1). Interestingly, two of these enzymes were highlighted as important for growth on mucin using a TnSeq library^16^.

Concerning GAG and starch utilisation, why the AM genome would encode enzymes specific for these substrates, when it does not use them as a nutrient source directly, remains unresolved at present. One possibility we explored is that the large mucin structures being taken up by AM also commonly contain GAG and starch contaminants. The PL38, GH105, sulfatases, GH13 and GH77 enzymes would be present internally to break these down, either for use as a nutrient source or to expel them. We attempted to test the idea that GAGs areused as a nutrient source through growths with different proportions of GAG-to-mucin, but the decrease in growth with increasing GAG concentration suggests that this theory is incorrect (Supplementary Figure 1).

### The extraordinary import system of AM

An intriguing physiological phenomenon of AM is its unusual substrate import mechanisms. In a paper describing the first genetic manipulation of AM, Davey *et al.* highlighted a possible import system that implicated pili in the import of mucin^16^. They found that mucin was accumulating in intracellular compartments, dubbed “mucinosomes”. Nine CAZymes were associated with this likely import system (Mul1A) (Supplementary Table 1). Interestingly, these CAZymes were also either very highly conserved across *Akkermansia* species, predicted to be localised to the outside of the cell, and/or highlighted as important for growth on mucin. It is likely that some of these different observations are connected. Complementing the observation of mucinosomes, was the types of enzyme activity detected on the surface of AM in this report. Prominent activities included GH16 and sialidase activities, and fucosidase activities were also observed by Shuoker *et al*.^13^, but other glycobiology was much lower. This suggests that most of the degradation of mucin is taking place inside the cell (Figure 6). Finally, growth of AM on human milk has also been observed ^18^, so there must also be a way to import human milk oligosaccharides and structurally similar GH16-derived O-glycan fragments produced on the surface of the cell. Future work on characterising this import system will be paradigm shifting in the microbiology field.

### CAZymes where no activities could be identified

In this report we have acknowledged several puzzles where activity could not be elucidated (Supplementary Table 1). Below is a discussion on these.

#### GH31

The second GH31, Amuc_1870^GH31^ is from subfamily 1, which has 84 characterised members and all bacterial members exhibit α-glucosidase activity. This enzyme did have activity against pNP-α-Glc, but no activity against substrates tested could be found, including those composed of α-glucose: maltose, dextran, starch, and laminarin (Supplementary Figures 10-14). A model of Amuc_1870^GH31^ was superimposed on to crystal structures of GH31 enzyme-substrate complexes with α-linked glucose disaccharides: isomaltose, nigerose, and kojibiose (Supplementary Figure 14). Only the isomaltose overlay indicated that this substrate could be accommodated, so we then tried isomaltose and pullulan (maltotriose linked via α1,6-linkages), but also found no activity. Interestingly, however, the closest characterised homologues prefered isomaltose over maltose^31, 32^. Finally, we also tested this enzyme against glucosylceramide (core of gangliosides and globosides) and trehalose, which is an unusual disaccharide produced by some organisms and is two glucose monosaccharides linked via α1,1 bond. There was no activity against either of these substrates and for the glucosylceramide this was confirmed by HPAEC (Supplementary Figure 14). AM did not grow on either maltose, starch, amylose, amylopectin, or pullulan (Supplementary Figure 1). This enzyme is one of the unsolved puzzles (Supplementary Table 1).

#### GH57

Amuc_1868^GH57^ is one of the unsolved puzzles (Supplementary Table 1). We found activity on pNP-α-Glc, but none against the wide variety of carbohydrate substrates tested in this report. Interestingly, the protein sequence for Amuc_1868^GH57^ is one of the most highly conserved across all the *Akkermansia* species considered here (>90 % identity over five species) and is always present in different strains. Adding further intrigue is that the TnSeq data previously reported by Davey *et al.* highlighted this enzyme as important for growth on mucin (Supplementary Table 1)^16^. The GH57 enzyme family has been further categorised in to four groups and the sequence of this enzyme has been previously compared to other GH57 enzymes based on sequence motifs^33^. Intriguingly, this enzyme is classified as an amylase-like protein as it is lacking a catalytic nucleophile.

#### GH63

Amuc_1260^GH63^ is also an unsolved puzzle (Supplementary Table 1) with activity not being identified on any substrate, including pNP-sugars. It is possible that these enzymes are involved in another function besides mucin breakdown. AM has 50 putative glycosytransferase enzymes and it is probable that at least some of these will contribute to synthesising capsular and/or exopolysaccharides. The composition of these carbohydrate structures from microorganisms is a relatively understudied area of glycobiology and the types of structures produced by different microbes are likely highly variable. Therefore, some of the ‘puzzle’ enzymes discussed in this report may target these structures, possibly to recycle the component sugars and facilitate cell growth and division. For example, there was no activity observed for Amuc_1868^GH57^ but it is highly conserved across *Akkermansia* species and was important for growth on mucin^16^.

Amuc_1216^GH177^ and Amuc_0623^GHnc^: These enzymes have both been associated with the breakdown of sialic acid^13, 34^. We could not find activity against any sialylated substrates or pNP-sugars in this work.

---

**Supplementary Figure 1.**
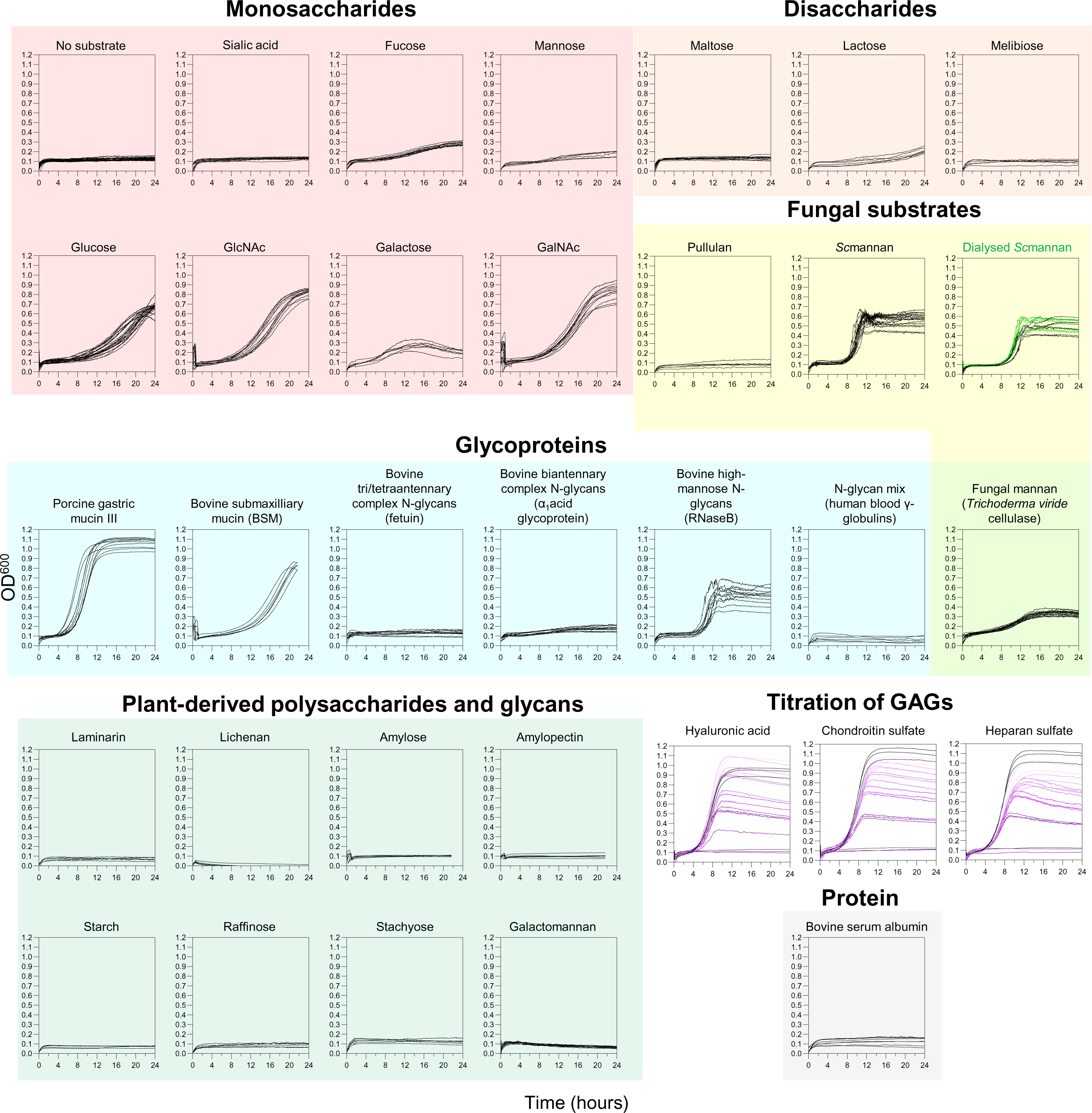
Growth of *A. muciniphila* ATCC BAA-835 on a variety of glycans and polysaccharides. *A. muciniphila* was grown anaerobically in minimal media containing a different potential nutrient sources. Growth was monitored continuously at OD600 using a 96-well plate in a plate reader. Concentrations of substrates are listed in Supplementary Table 2. For the dialysed Scmannan, the black curves are original sample and the green is dialysed. For the GAG substrates, this was completed with varying concentrations of mucin:GAG. The black lines are the mucin growth curves and the darkest purple are the GAGs alone. The lighter the purple gets the more mucin in those growths. Each line is one bacterial culture.

**Supplementary Figure 2.**
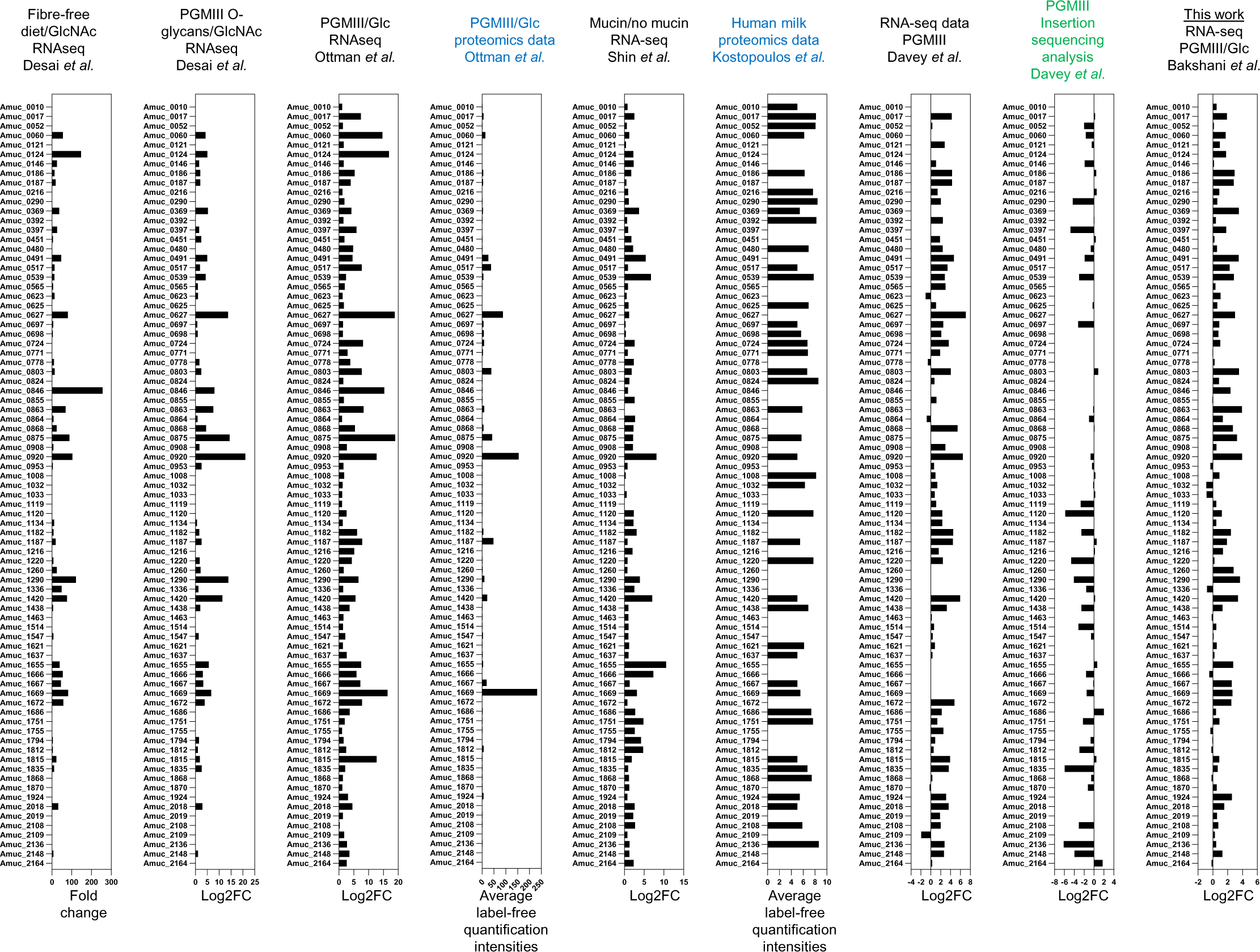
Transcriptomic, proteomic and Tnseq data of AM from this work and previous publications. The bar charts show the data from different labs and techniques that assess gene upregulation, protein expression, or the reduction in growth when a gene is disrupted. Black, blue, and green indicate RNAseq, proteomics, and INSeq data, respectively.

**Supplementary Figure 3.**
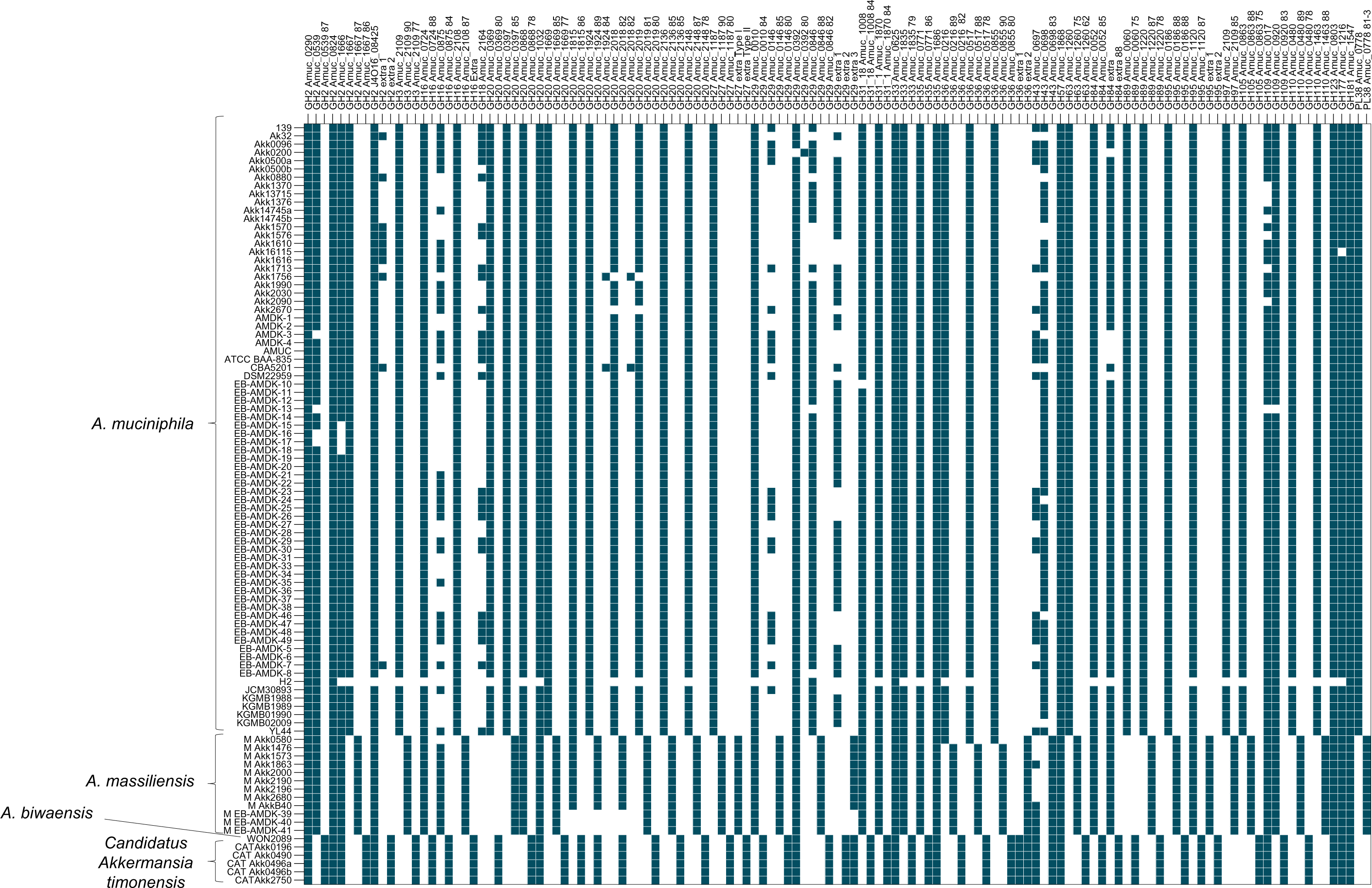
Presence of homologues of the putative CAZymes from ATCC BAA-835 across other *A. muciniphila* strains and *Akkermansia* species. The heat map shows the presence of homologues to the CAZymes in ATCC BAA-835 (dark teal). When the homologue is <90 % identical, <80 % identical, and <70 % identical then these are placed in a separate columns. For instance, the homologues for Amuc_0290^GH2^ are all >90 % across strains and species, whereas the homologues for Amuc_1667^GH2^ all have lower percentage identity across the species but >90 % conservation within strains. The y axis is split up according to species and the x axis is in numerical GH order.

**Supplementary Figure 4.**
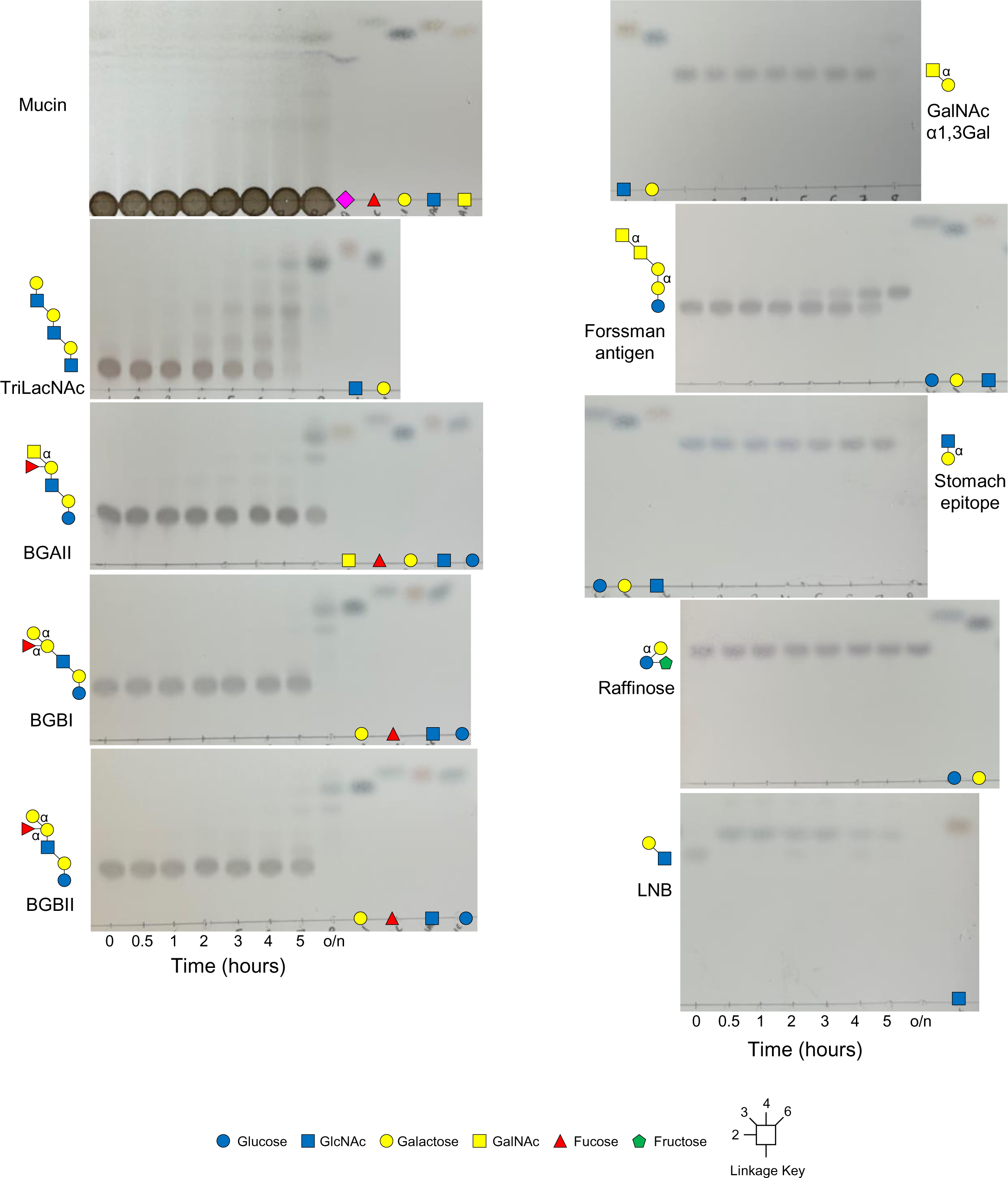
Whole cell assay #1. Thin layer chromatography results of whole cell assays of PGM III-grown cells against different substrates. The reaction was carried out at 37 °C, a sample removed at different time points, and boiled to stop enzyme activities. Monosaccharide standards are shown in the right lanes.

**Supplementary Figure 5.**
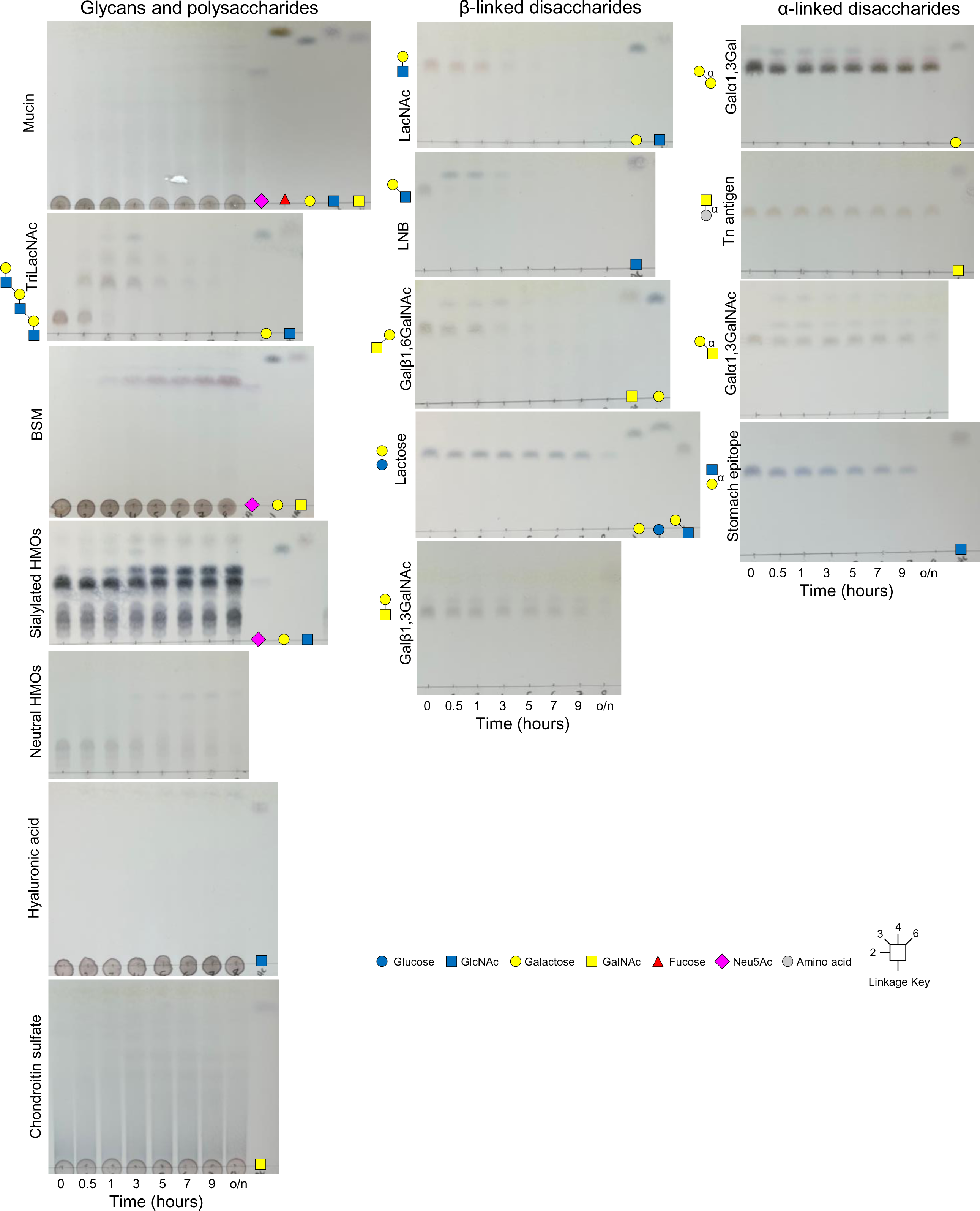
Whole cell assay #2. Thin layer chromatography results of whole cell assays of PGM III-grown cells against different substrates. The reaction was carried out at 37 °C, a sample removed at different time points, and boiled to stop enzyme activities. Monosaccharide standards are shown in the right lanes.

**Supplementary Figure 6.**
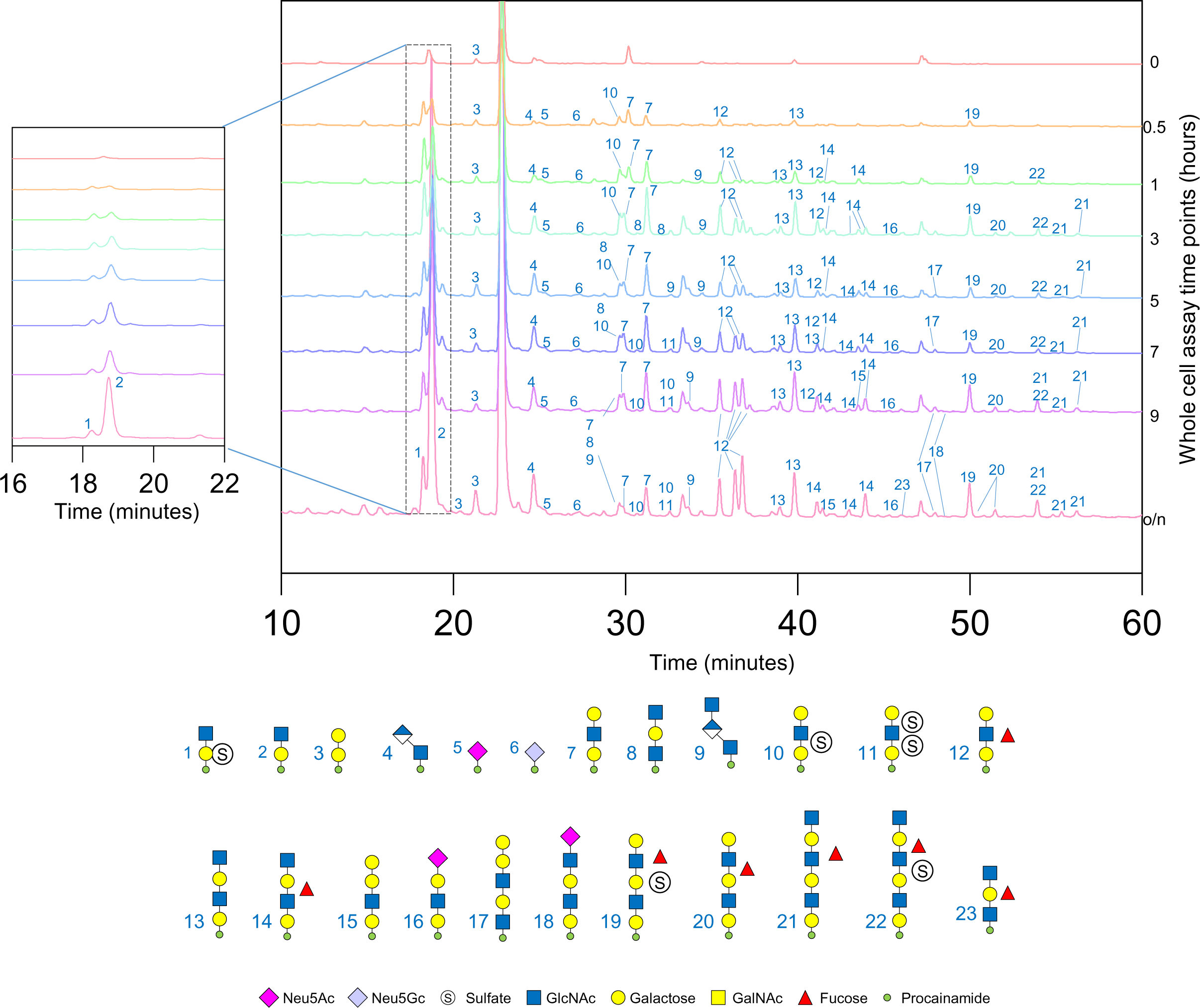
Characterisation of surface enzymes activity of AM using whole cell assays. The glycan products from the different whole cell assay timepoints were labelled with procainamide at the reducing end and analysed by LC-FLD-ESI-MS. The results show a release of a variety of different O-glycan fragments, predominantly with galactose at the reducing end, which is indicative of GH16 endo-O-glycanase activity. The panel on the left emphasises glycans 3 and 4 and the panel on the right emphasises glycans eluting between 25-60 minutes.

**Supplementary Figure 7.**
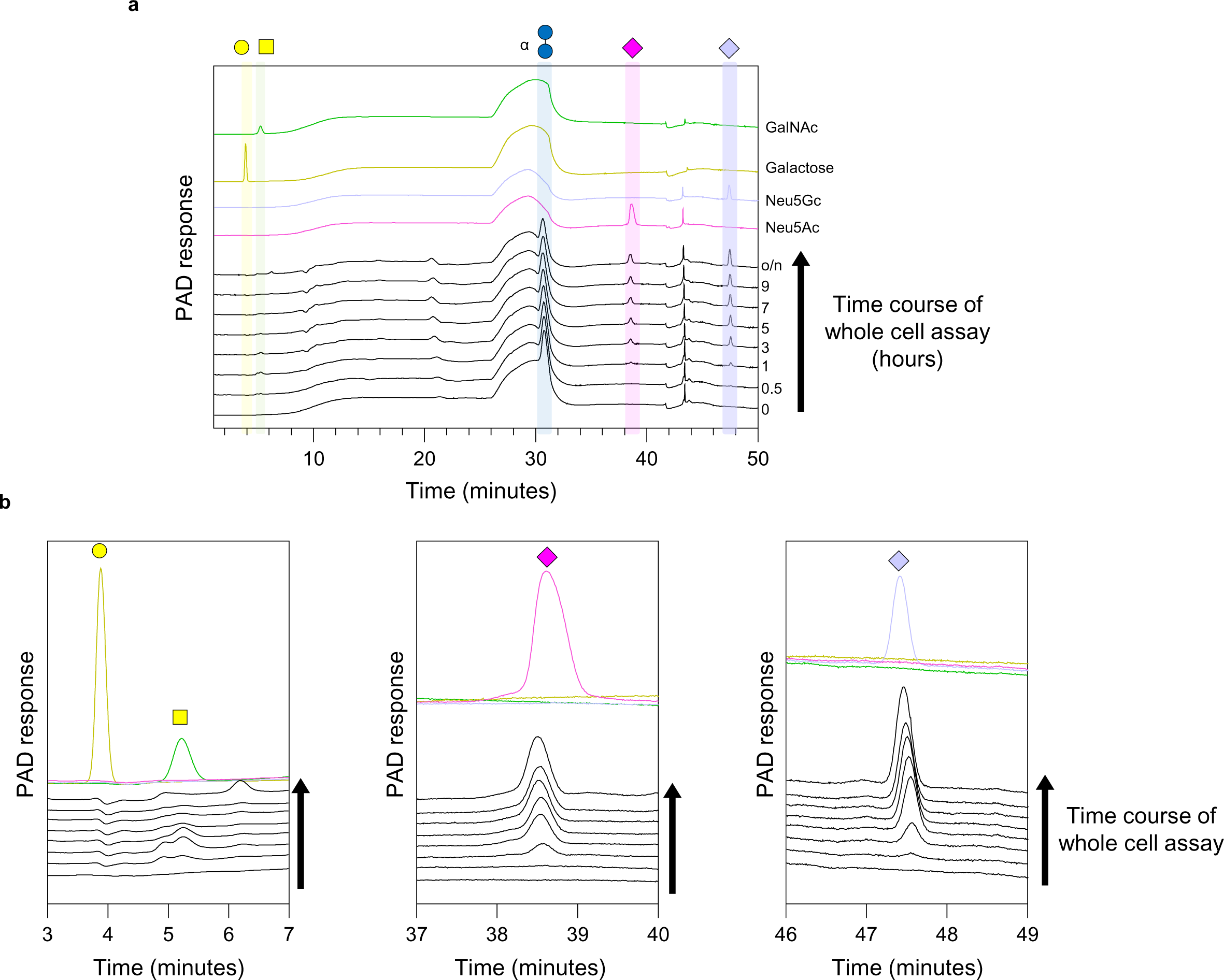
HPAEC-PAD of BSM whole cell assays time points. Samples from Supplementary Figure 9 of PGM III-grown whole cell assays against BSM were analysed by HPAEC-PAD. **a,** The full chromatograms are shown alongside standards of the different expected monosaccharides (colourful lines). **b,** Certain sections of the chromatograms are shown in more detail and stacked to show the increase in sialic acids produced over time.

**Supplementary Figure 8.**
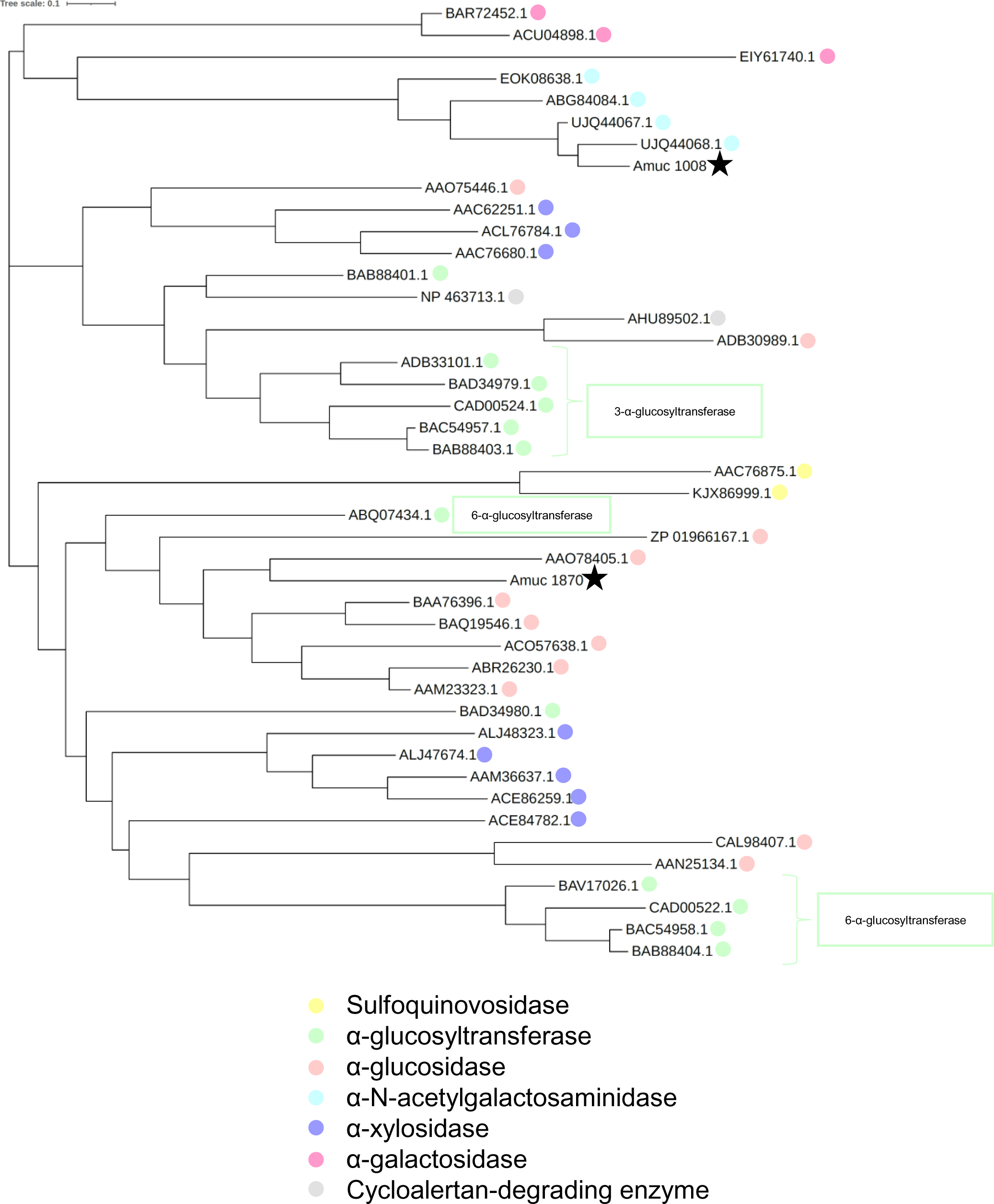
Phylogenetic tree of characterised GH31 family members with those from *A. muciniphila* ATCC BAA-835. The sequences of the GH31 family members with reported activities (CAZy database) and the ones from *A. muciniphila* ATCC BAA-835 were compared as described in the methods. The different specificities are indicated by different colours and the *A. muciniphila* ATCC BAA-835 enzymes are highlighted by black stars. Some more specific information about activity is supplied where possible. The enzymes are represented by their accession numbers of locus tags. Different specificities cluster in this analysis and the observed activity of the two *A. muciniphila* ATCC BAA-835 enzymes correlates to where they cluster on the tree.

**Supplementary Figure 9.**
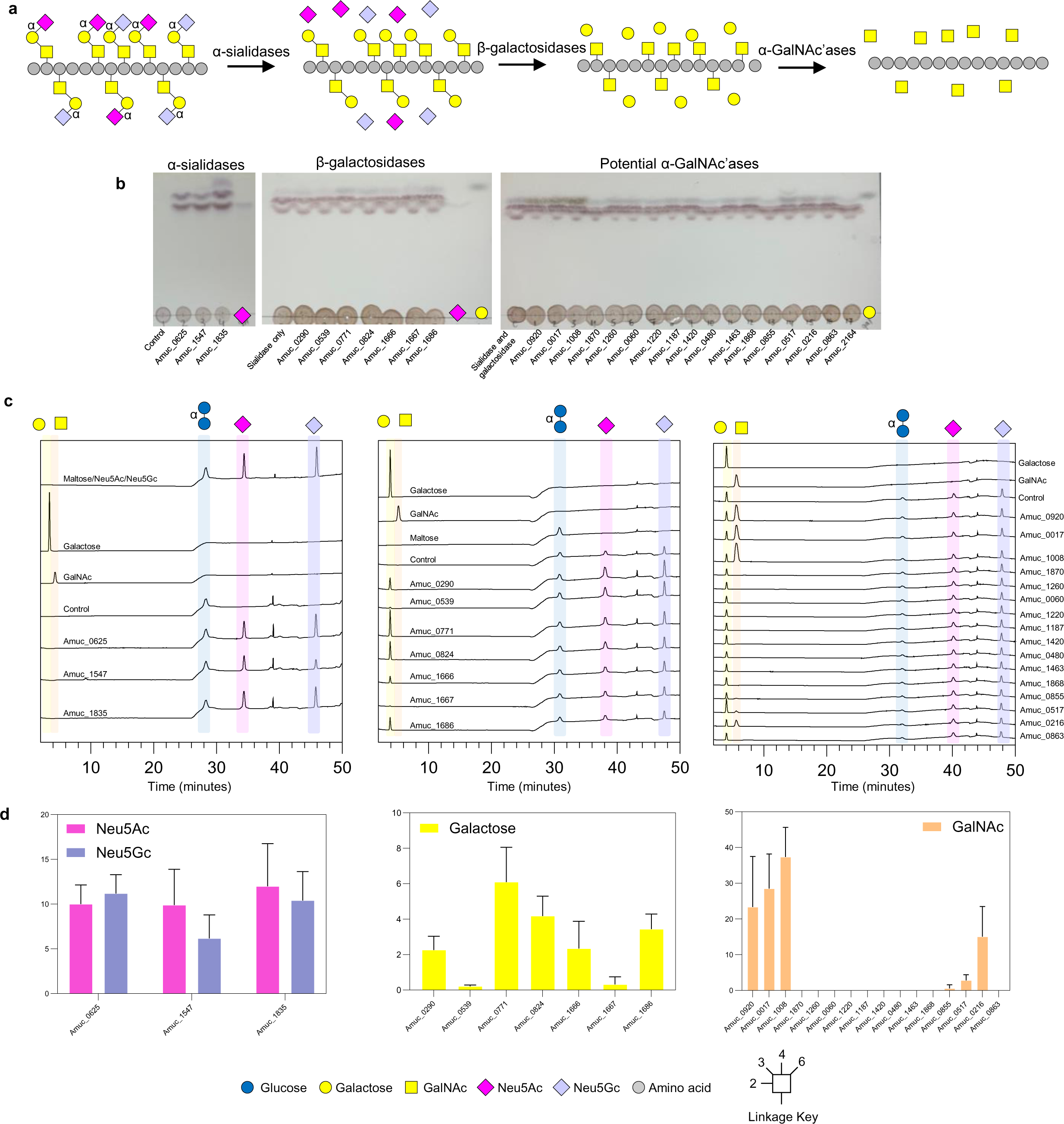
Sequential degradation of O-glycans on BSM with CAZymes from *A. muciniphila* BAA-835. The sialidases, β-galactosidases, and CAZymes associated with α-linked monosaccharide hydrolysis were added sequentially to understand specificity of the CAZymes. **a,** Structural features that are expected in bovine submaxilliary mucin (left) and how it is broken down in this experiment. Only alpha linkages are labelled apart from the core GalNAc monosaccharides which are also alpha linkages. **b**, Thin layer chromatography results of the sequential degradation. Standards have also been included on the left of the TLCs. **c**, The chromatograms of the assays analysed by HPAEC-PAD. Maltose was used as the internal standard and other standards were run separately. **d**, The areas of the peaks were quantified. The data are from three distinct reactions.

**Supplementary Figure 10.**
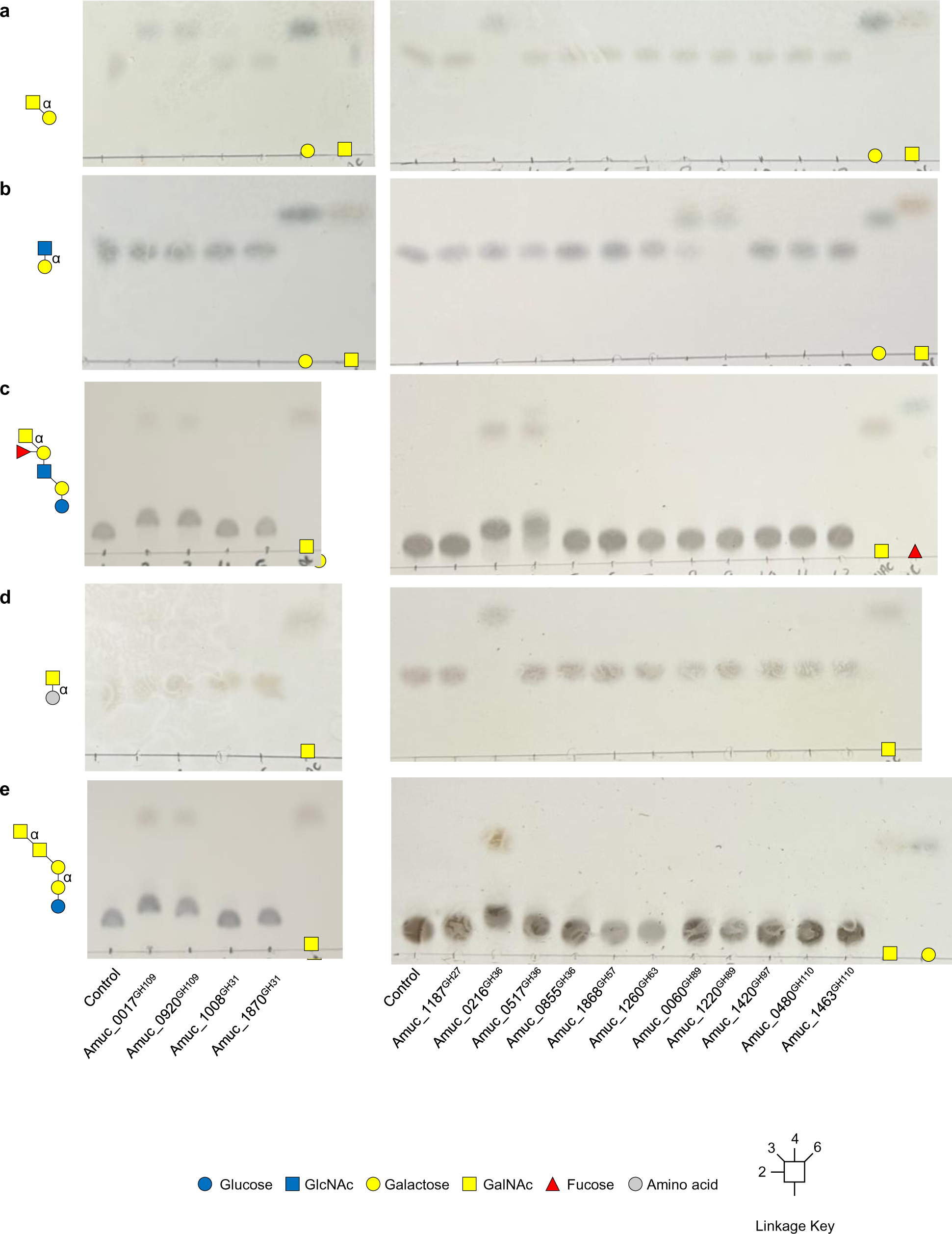
Activity of GH enzymes from *A. muciniphila* BAA-835 against defined oligosaccharides capped with α-GalNAc or α-GlcNAc. **a,** GalNAcα1,3Gal. **b**, GlcNAcα1,4Gal. **c**, Blood group A II. **d**, Tn antigen. **e**, Forssman antigen. Standards have also been included on the TLCs on the right. Enzyme assays were carried out at a final substrate concentration of 1 mM, pH 7, 37 °C, overnight, and with 1 μM enzyme.

**Supplementary Figure 11.**
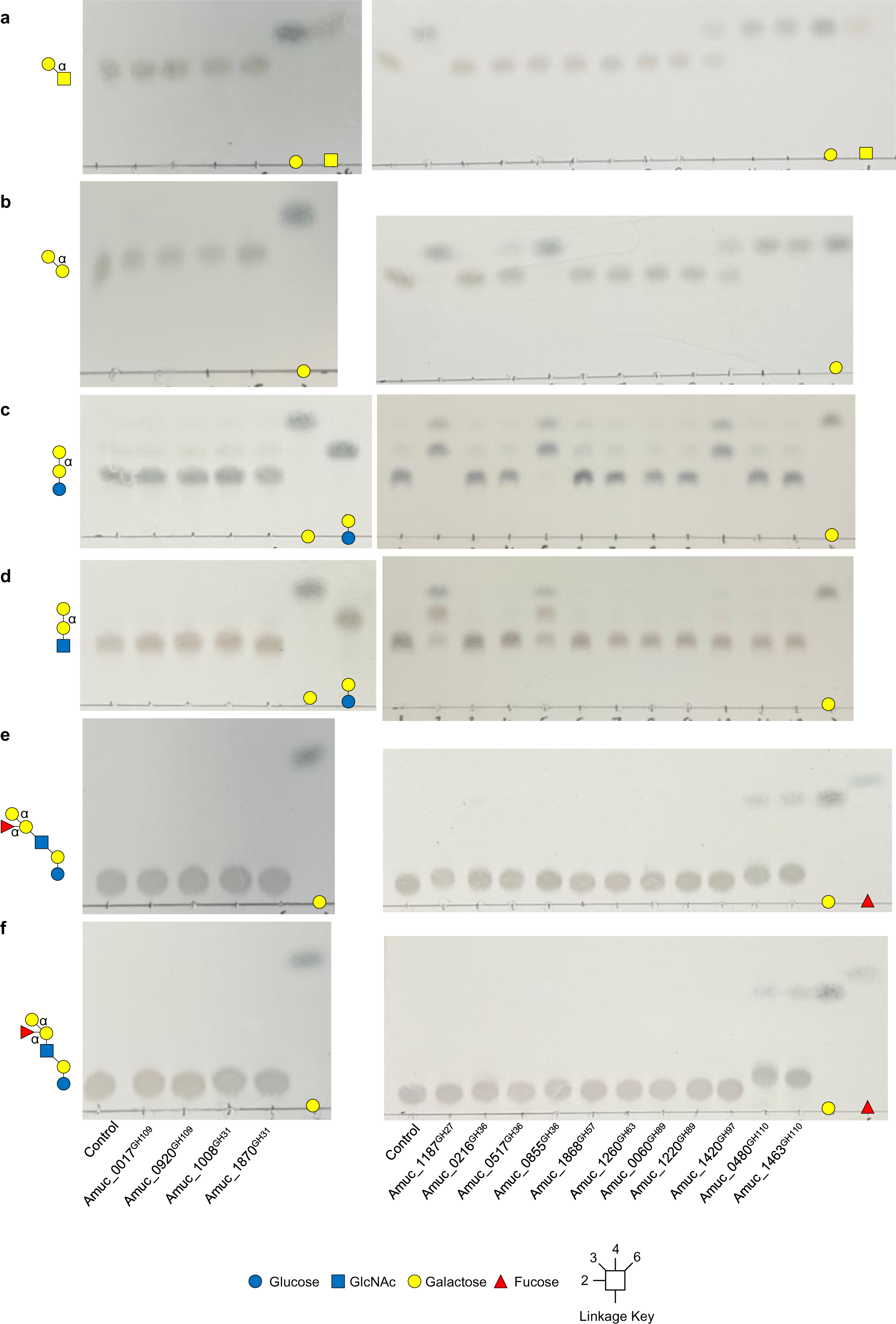
Activity of GH enzymes from *A. muciniphila* BAA-835 against defined oligosaccharides capped with α-GalNAc or α-GlcNAc. **a,** Galα1,3GalNAc. **b**, Galα1,3Gal. **c**, globotriose. **d**, P1 antigen. **e**, Blood group B I. **f**, Blood group B II. Standards have also been included on the TLCs on the right. Enzyme assays were carried out at a final substrate concentration of 1 mM, pH 7, 37 °C, overnight, and with 1 μM enzyme.

**Supplementary Figure 12.**
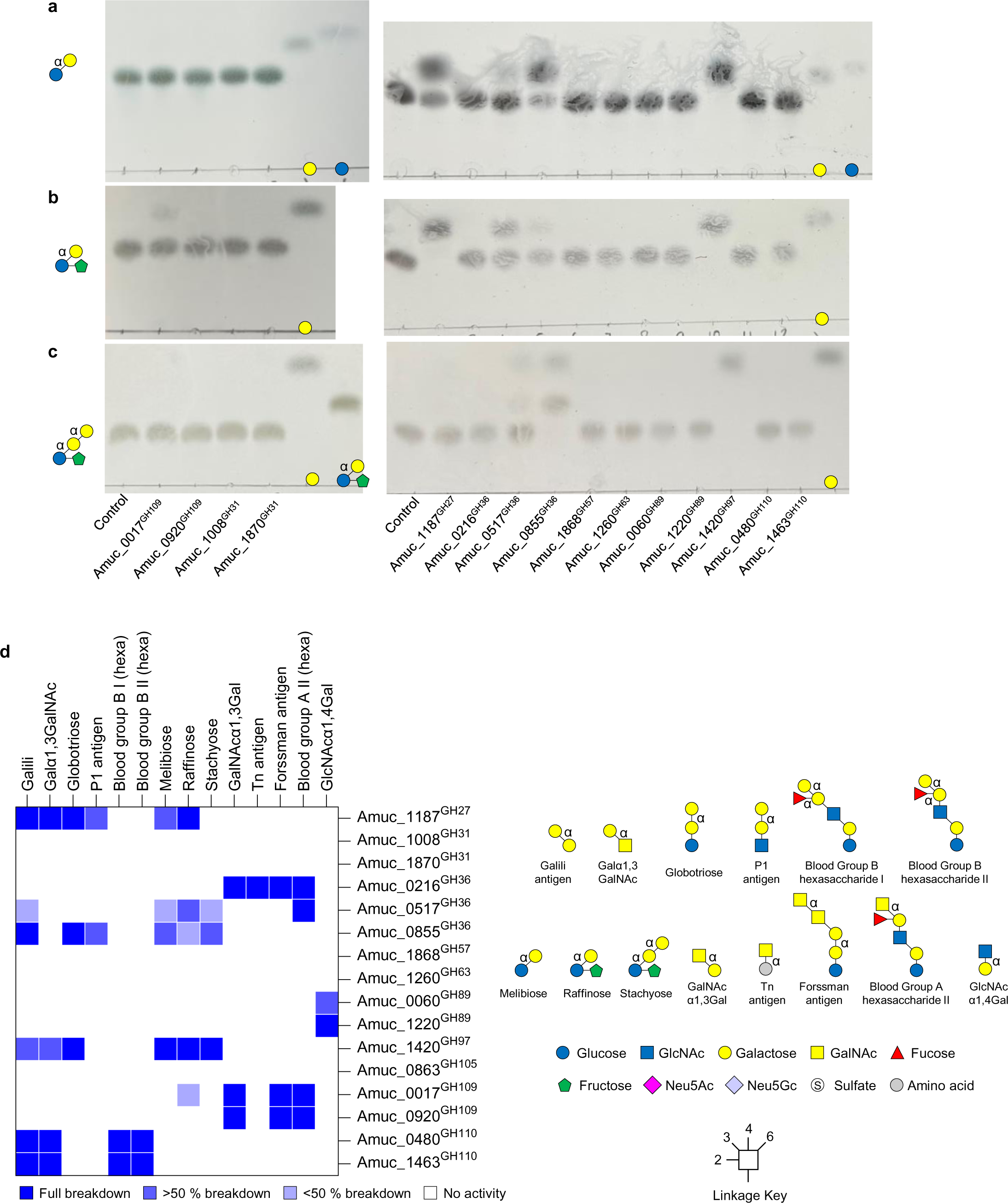
Activity of GH enzymes from *A. muciniphila* BAA-835 against raffinose-type defined oligosaccharides. **a,** melibiose. **b**, raffinose. **c**, stachyose. Standards have also been included on the TLCs on the right. Enzyme assays were carried out at a final substrate concentration of 1 mM, pH 7, 37 °C, overnight, and with 1 μM enzyme. **d**, Heat map of recombinant enzyme activities against defined oligosaccharides. The dark blue and white indicate full and no activity, respectively, and partial activities are represented by the lighter blues.

**Supplementary Figure 13.**
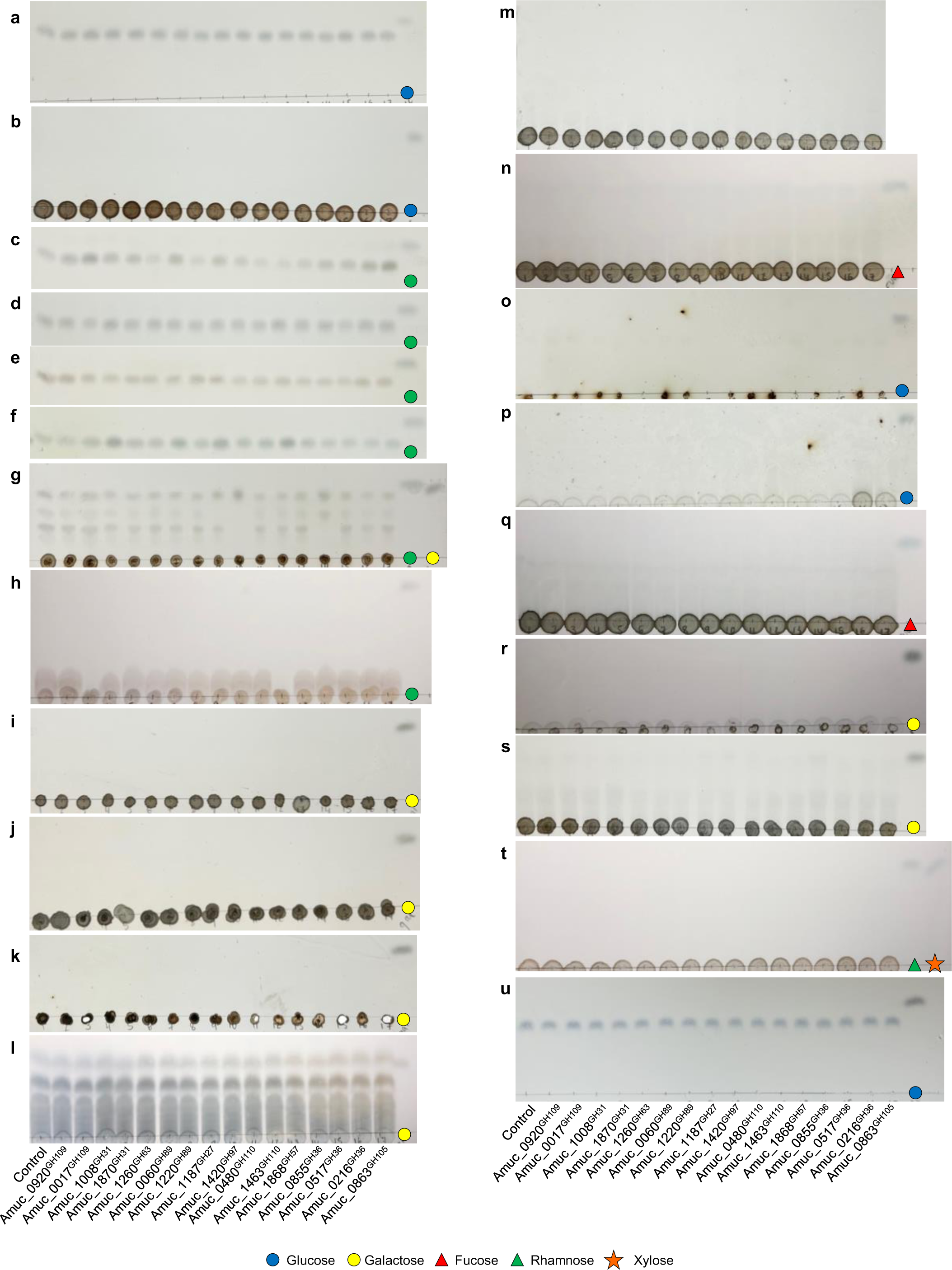
Activity of GH enzymes from *A. muciniphila* BAA-835 against a wide variety of substrates. **a,** maltose. **b**, dextran. **c**, α1,2-mannobiose. **d**, α1,3-mannobiose **e**, α1,4-mannobiose **f**, α1,6-mannobiose. **g,** galactomannan. **h,** RNaseB (high-mannose N-glycans). **i,** L-carrageenan. **j,** K-carrageenan. **k,** I-carrageenan. **l,** porphyran (L). **m,** ulvan. **n,** fucan. **o,** starch. **p,** laminarin. **q,** fucoidan. **r,** agarose. **s,** porphyran (W). **t,** ulvan (entero). **u,** isomaltose. Standards have also been included on the TLCs. Enzyme assays were carried out at a final substrate concentration of 1 mM, pH 7, 37 °C, overnight, and with 1 μM enzyme.

**Supplementary Figure 14.**
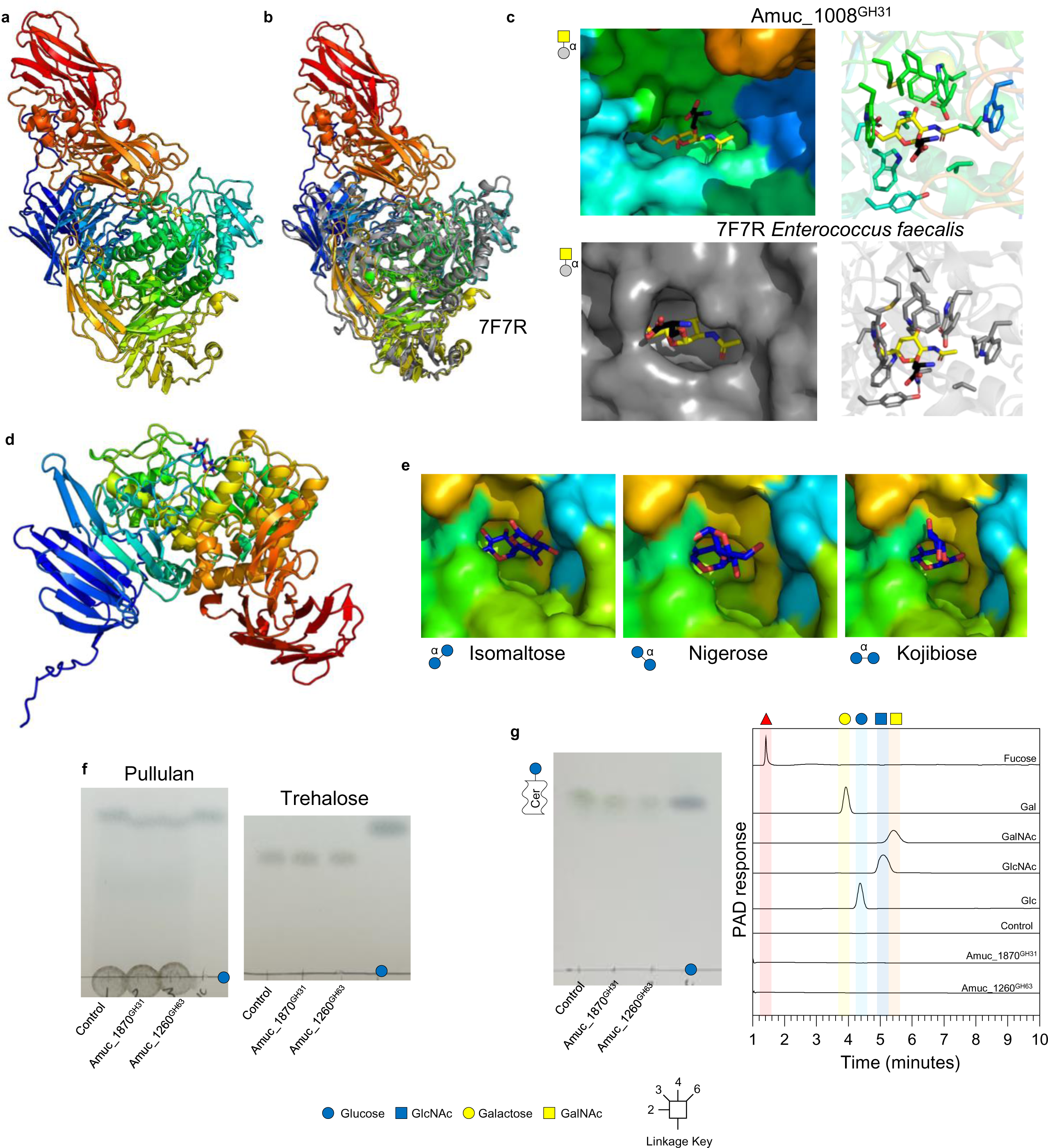
GH31 family members from *A. muciniphila* ATCC BAA-835. **a.** Amuc_1008^GH31^ cartoon model coloured rainbow blue-red from the N- to C-terminus, respectively. GalNAc (yellow) from 7F7Q was overlaid into the active site. **b.** The cartoon of Amuc_1008^GH31^ (rainbow) overlaid with the *E. faecalis* GH31 (grey) with the same specificity showing that they overlay well in terms of secondary structures, but the *A. muciniphila* enzyme has additional modules. **c.** Top: Amuc_1008^GH31^ active site surface overlaid with Tn-antigen from 7F7R *Enterococcus faecalis* GH31_18. Bottom: active site of 7F7R for comparison. Left and right panels show the surface of the enzymes and residues interacting with the sugar in the -1 subsite as sticks, respectively. **d.** Amuc_1187^GH31^ cartoon model coloured rainbow blue-red from the N- to C-terminus, respectively. Isomaltose (blue) from 3MKK was overlaid into the active site. **e.** Amuc_1187^GH31^ active site surface overlaid with α-linked glucose disaccharides from different crystal structures. From left to right: Isomaltose from 3MKK, Nigerose from 7WJC, and Kojibiose from 7WJF. Only the isomaltose was not making obvious steric clashes with the predicted protein structure. **f.** Activity against pullulan and trehalose. Isomaltose reactions are in Supplementary Figure 16. **f.** Activity against glucosylceramide. Left panel is the TLC and right panel are the assays run on the HPAEC alongside standards. Standards have also been included on the TLCs. Enzyme assays were carried out at a final substrate concentration of 1 mM, pH 7, 37 °C, overnight, and with 1 μM enzyme.

**Supplementary Figure 15.**
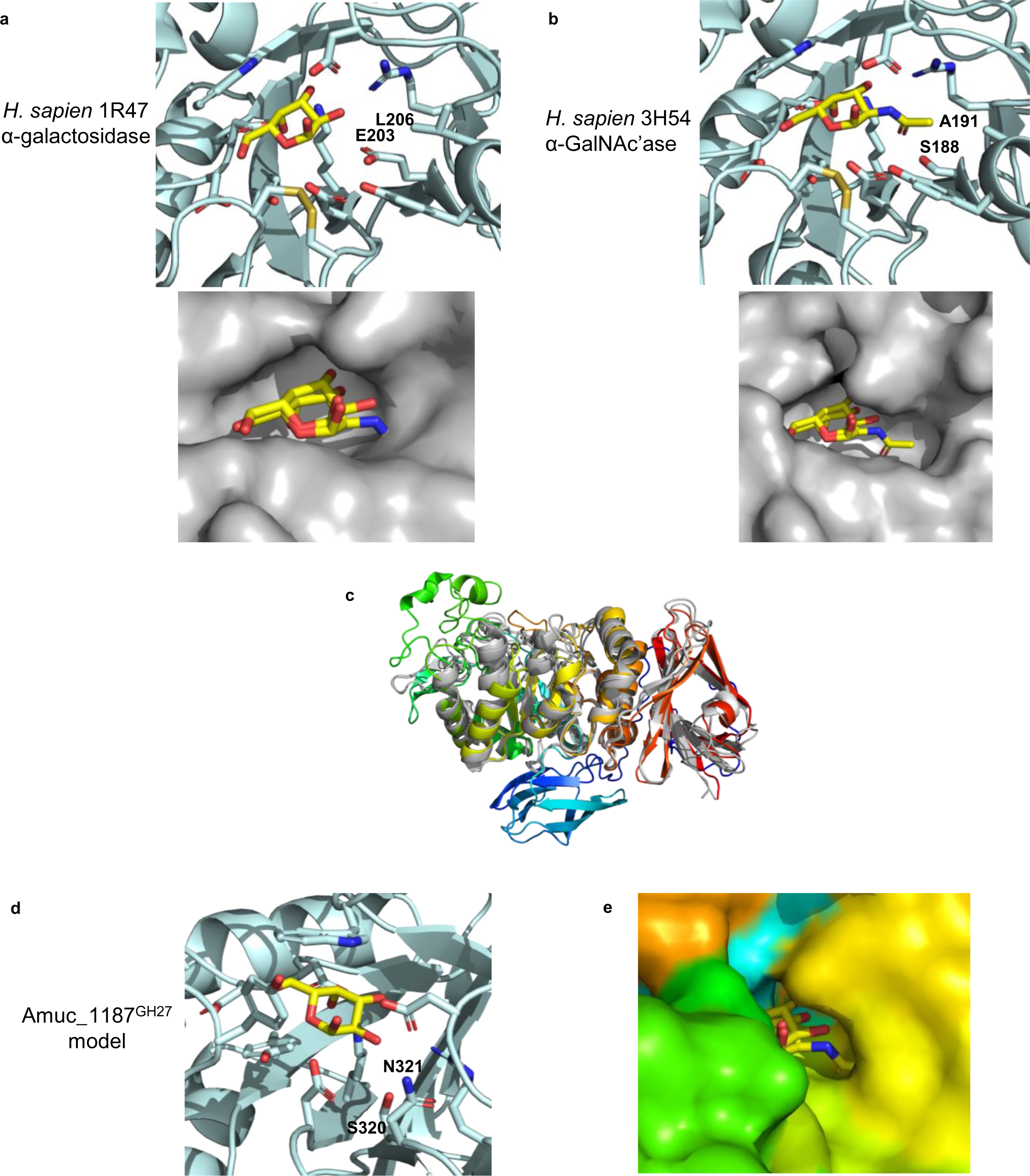
Models of the GH27 family member from *A. muciniphila* ATCC BAA-835. **a.** Human GH27 α-galactosidase with galactose in the -1 subsite and **b.** Human GH27 α-GalNAc’ase with GalNAc in the -1 subsite. The residues coordinating these -1 sugars differ only in the residues that have been labelled. The larger residues coordinate the galactose in 1R47 and the smaller residues allow the accommodation of GalNAc in 3H54. **c.** An overlay of Amuc_1187^GH27^ (rainbow, N- to C-termini are blue-red, respectively) with the two *H. sapien* enzymes (grey) show that the overall fold is largely similar. **d.** A model of Amuc_1187^GH27^ with the ligand from 1R47 and the two equivalent residues. These are relatively large residues, so the enzyme has specificity for galactose rather than GalNAc. **e.** A surface representation of the model of Amuc_1187^GH27^ to show the pocket for galactose more clearly.

**Supplementary Figure 16.**
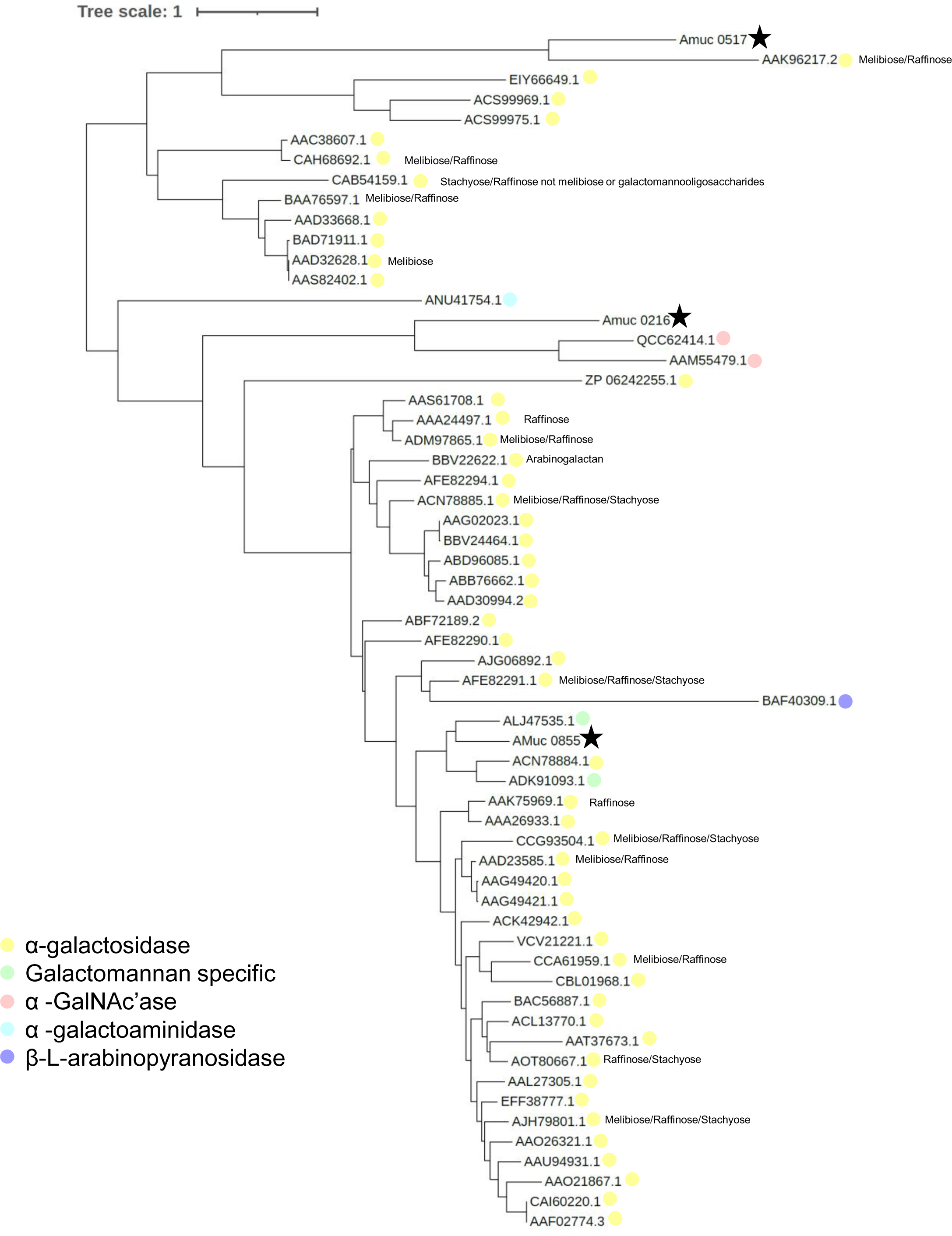
Phylogenetic tree of characterised GH36 family members with those from *A. muciniphila* ATCC BAA-835. The sequences of the GH36 family members with reported activities (CAZy database) and the ones from *A. muciniphila* ATCC BAA-835 were compared as described in the methods. The different specificities are indicated by different colours and the *A. muciniphila* ATCC BAA-835 enzymes are highlighted by black stars. The enzymes are represented by their accession numbers of locus tags.

**Supplementary Figure 17.**
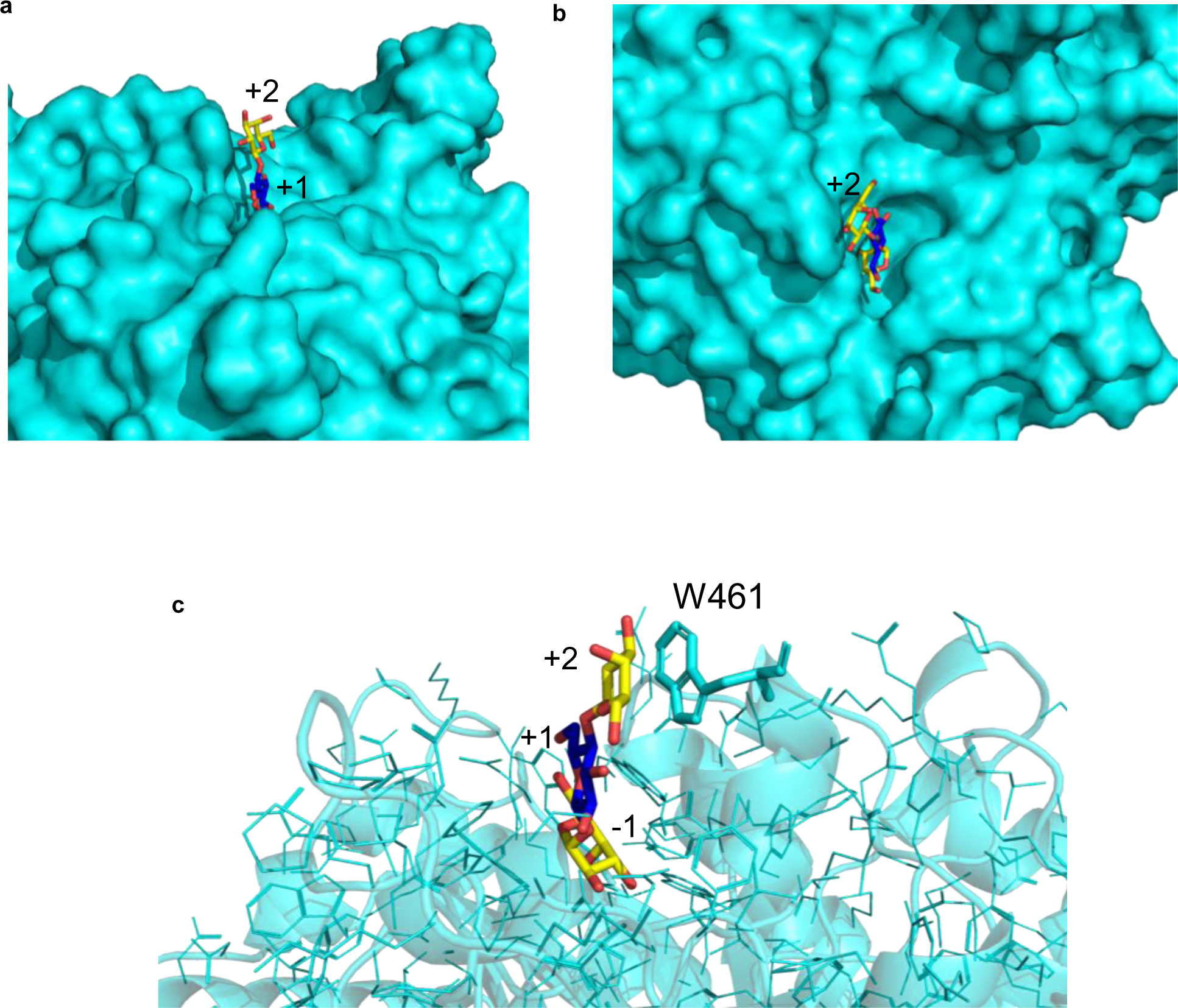
The structural model of the Amuc_1420^GH97^. **a,** The active site of the model is viewed in surface mode from the side. Overlaid into the active site is a trisaccharide (lactose that also has an α-linked galactose to the Glc through a 1,1-linkage) from the *Bacteroides thetaiotaomicron* BT1871 structure. b, same structure looking down into the active site. c, a close up of the active site, showing the secondary structure as cartoon, the amino acid side chains as lines, apart from one tryptophan that may interact with the sugar in the +2 subsite.

**Supplementary Figure 18.**
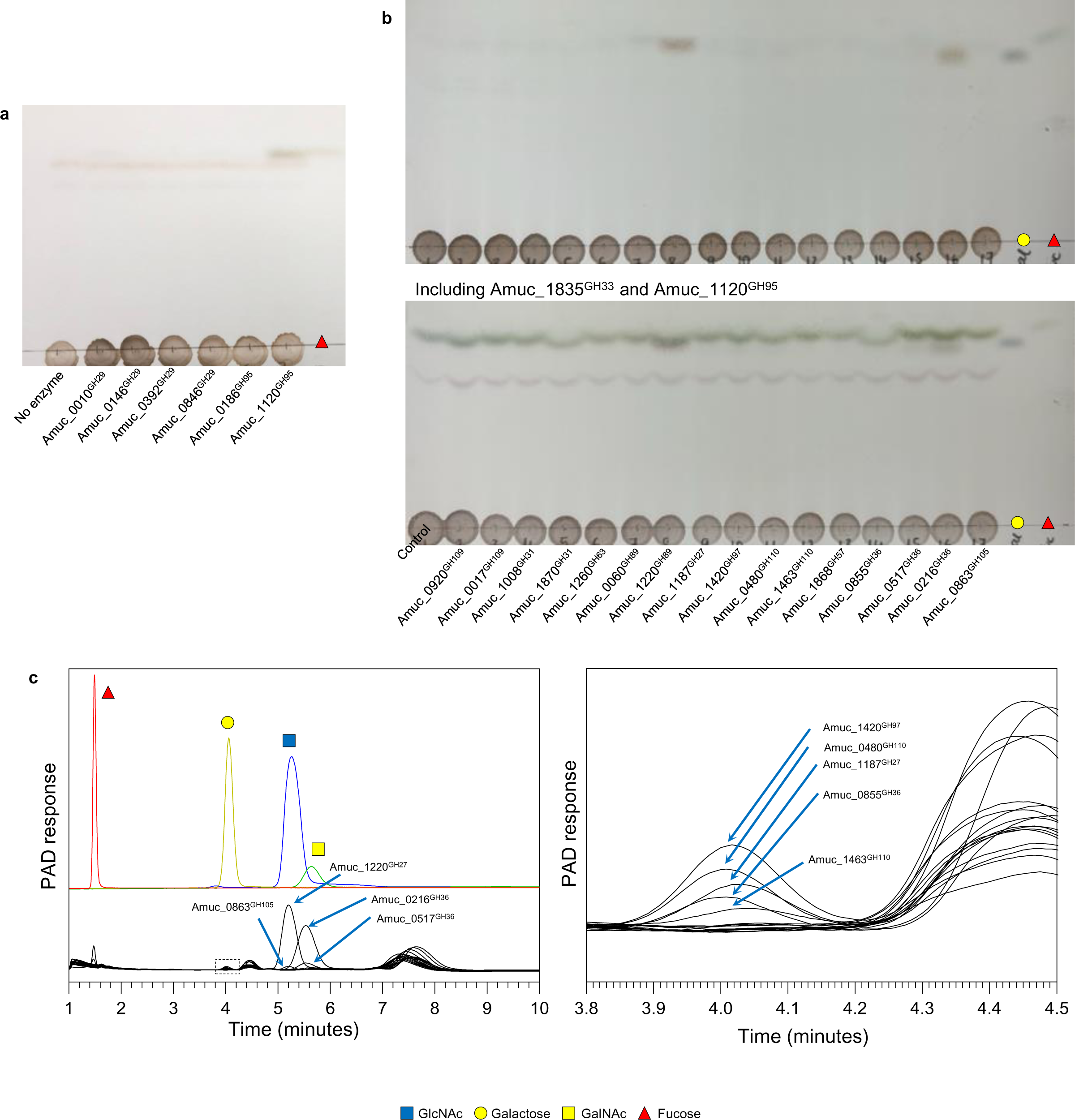
Degradation of PGM III with fucosidases and alpha-linkage CAZymes from *A. muciniphila* BAA-835 and analysis by thin layer chromatography. Enzyme assays were carried out pH 7, 37 °C, overnight, and with 1 μM enzymes. **a,** The activity of GH29 and GH95 family members against PGM III. **b**, The activity of GH enzymes associated with the hydrolysis of alpha-linkage removal. Top panel is without enzyme pre-treatment included and the HPAEC-PAD for these samples is presented in ‘c’. The bottom panel are the same assays with Amuc_1835^GH33^ and Amuc_1120^GH95^ included and the HPAEC-PAD for these samples is presented in Figure 2.

**Supplementary Figure 19.**
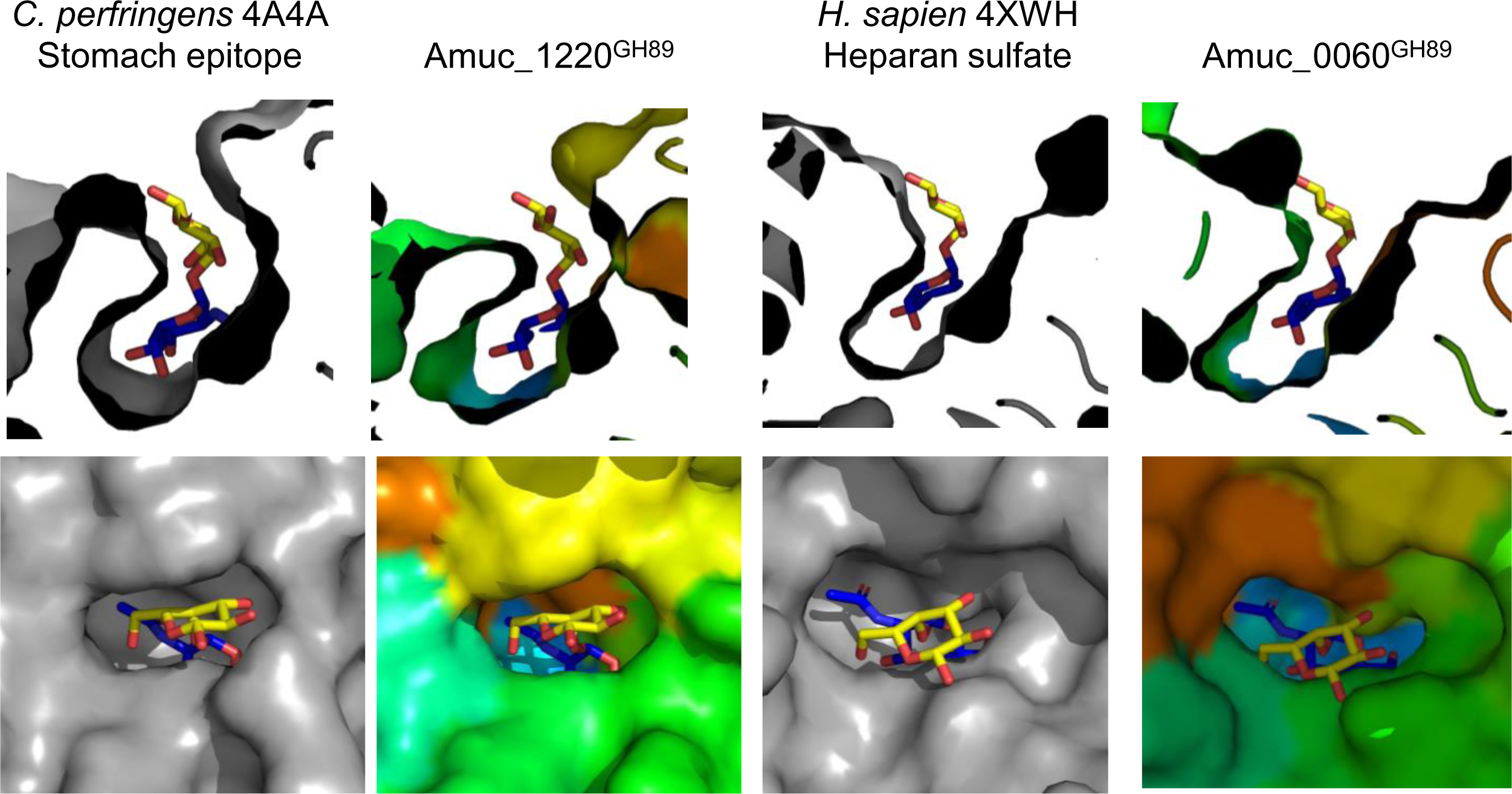
Structural insights into GH89 enzymes from *A. muciniphila* ATCC BAA-835. Two crystal structures (grey) with different substrate preferences from the GH89 family were compared to models of the two GH89 enzymes analysed in this work (rainbow). The *C. perfringens* 4A4A structure was crystallised with GlcNAcα1,4Gal in its active site and this was overlaid into the structure from *H. sapien* 4XWH and the two models. GlcNAc and Gal are blue and yellow, respectively. The top panels are cross sections through the active site to show how this disaccharide sits in there and the bottom panels are looking down at the entrance of the active site. The active site of the *C. perfringens* 4A4A structure packs tightly around the sugars in the -1 and +1 subsites, which has a considerable curve in its structure. The conformation of the Amuc_1220^GH89^ active site pocket is predicted to be very similar to this and accommodates this disaccharide well. The *H. sapien* 4XWH and Amuc_0060^GH89^model do not look like they can accommodate the angle of this substrate very well.

**Supplementary Figure 20.**
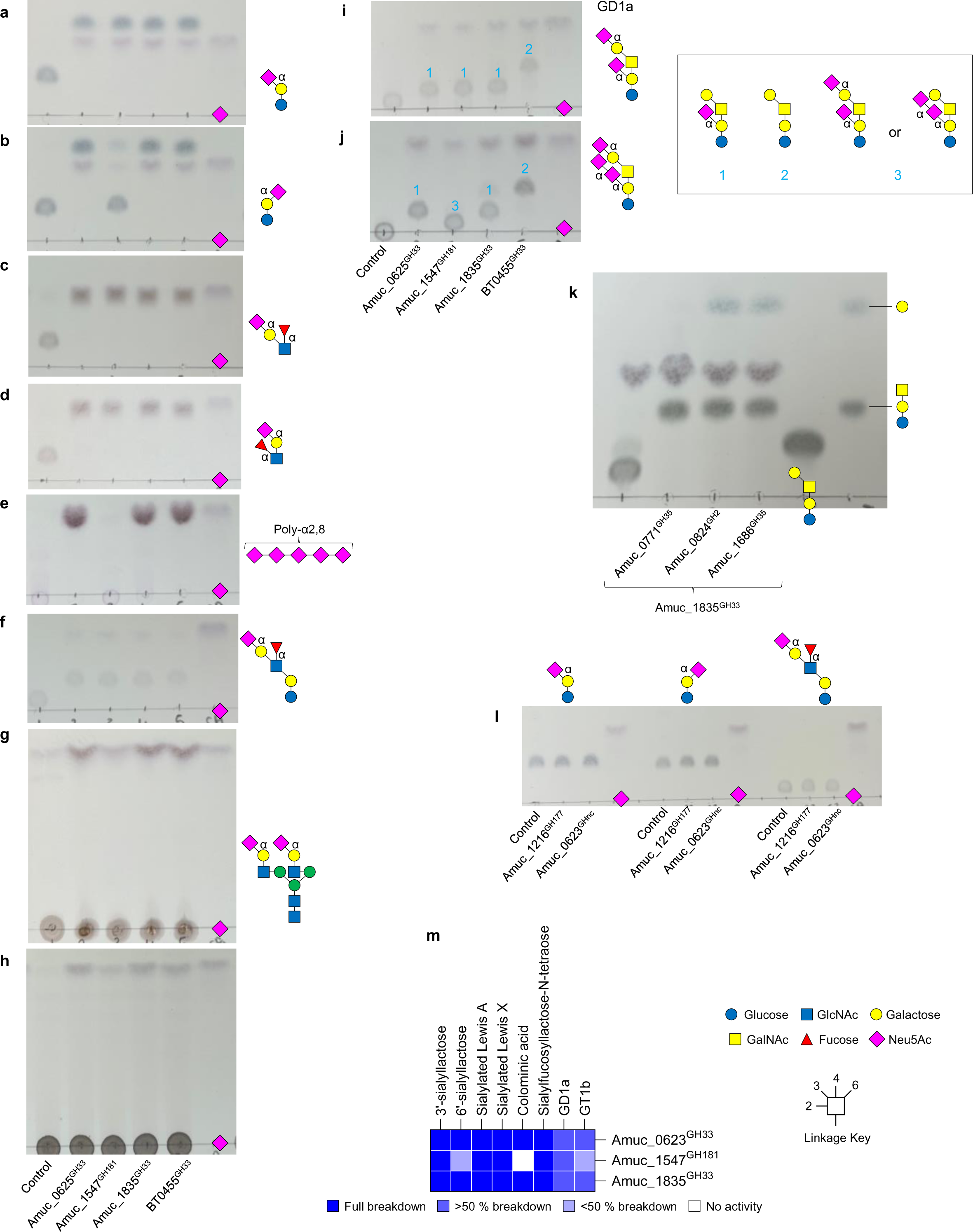
Activity of the GH33 and GH181 family members from *A. muciniphila* BAA-835 against defined oligosaccharides, polysaccharides, and mucin substrates. **a,** 3’-sialyllactose. **b**, 6’-sialyllactose. **c**, Sialylated Lewis A. **d**, Sialylated Lewis X. **e**, colominic acid. **f**, sialylfucosyllacto-N-tetraose. **g**, biantennary complex N-glycans (α_1_acid glycoprotein). **h**, PGM III. **i,** GD1a **j,** and GT1b. **k,** Exploring if removing the galactose also allows the release of the final sialic acid. **l,** Testing Amuc_1216^GH177^ and Amuc_0623^GHnc^ against three different sialylated substrates. **m,** Heat map of recombinant enzyme activities against defined oligosaccharides. The dark blue and white indicate full and no activity, respectively, and partial activities are represented by the lighter blues. Standards have also been included on the TLCs and substrates have been included on the right where possible. For the gangliosides (I and j) the structures of the different products are in the key to the right. Enzyme assays were carried out at a final substrate concentration of 1 mM (except N-glycans at 10 mg/ml and colominic acid at 25 mg/ml), pH 7, 37 °C, overnight, and with 1 μM enzymes. BT0455 from *Bacteroides thetaiotaomicron* has been used as a positive control a-i.

**Supplementary Figure 21.**
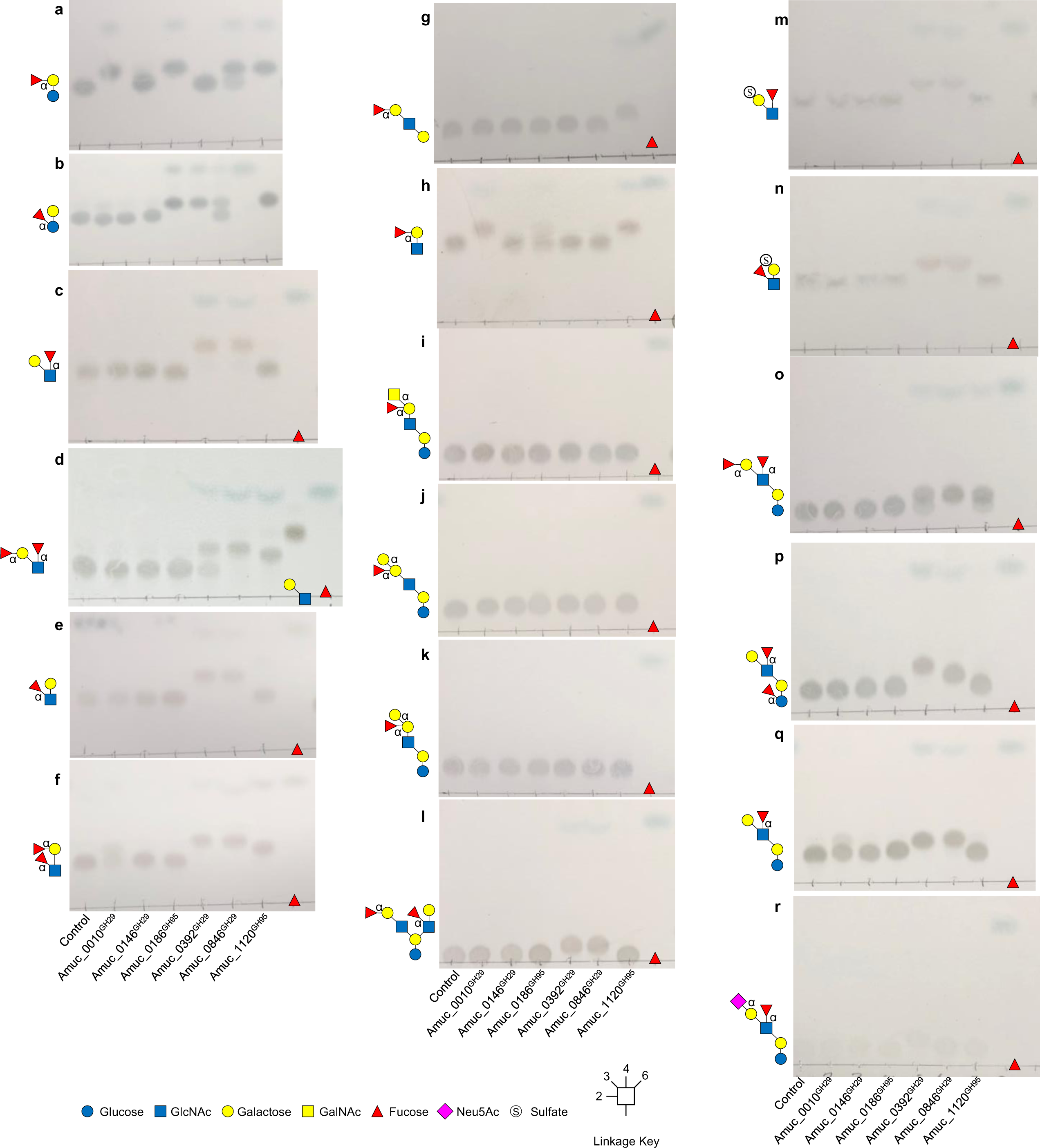
Activity of the GH29 and GH95 family members from *A. muciniphila* BAA-835 against defined oligosaccharides, Part 1. **a,** 2’-fucosyllactose. **b**, 3-fucosyllactose. **c**, Lewis A. **d**, Lewis B. **e**, Lewis X. **f**, Lewis Y. **g**, Blood group H I. **h**, Blood group H II. **i**, Blood group A II. **j**, Blood group B I. **k**, Blood group B II. **l**, difucosyllacto-N-hexaose. **m**, 3’-sulphated Lewis A. **n**, 3’-sulphated Lewis X. **o**, Lacto-N-difucohexaose I. **p**, Lacto-N-difucohexaose I. **q**, Lacto-N-fucopentaose II. **r**, Sialylfucosyllacto-N-tetraose. Standards have also been included on the TLCs. Enzyme assays were carried out at a final substrate concentration of 1 mM, pH 7, 37 °C, overnight, and with 1 μM enzyme.

**Supplementary Figure 22.**
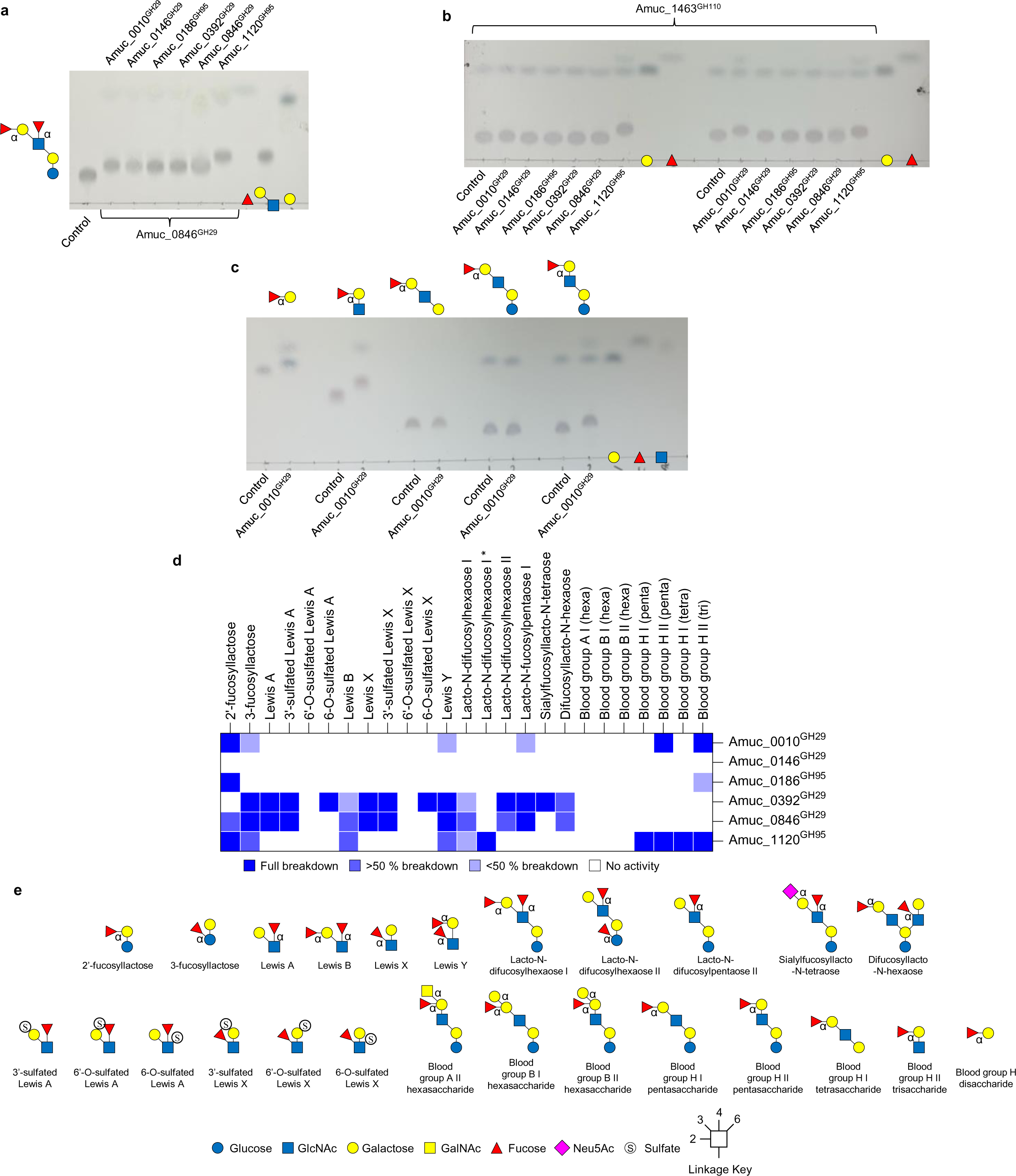
Activity of the GH29 and GH95 family members from *A. muciniphila* BAA-835 against defined oligosaccharides, Part 2. **a,** Exploring the full degradation of LNDFH I by pre-treating with Amuc_0846^GH29^ and then adding the other fucosidases. **b**, Fucosidase activity against different types of blood group B structures where the α-linked galactose has been removed. **c**, Amuc_0010^GH29^ activity against a range of blood group H structures. For all assays, standards have been included on the TLCs. Enzyme assays were carried out at a final substrate concentration of 1 mM, pH 7, 37 °C, overnight, and with 1 μM enzyme. **d**, A. heat map of recombinant enzyme activities against defined oligosaccharides. The dark blue and white indicate full and no activity, respectively, and partial activities are represented by the lighter blues. The asterisk indicates that one fucose has been remove by Amuc_0846^GH29^. **e**, Structures of the defined substrates used to characterise the fucosidases.

**Supplementary Figure 23.**
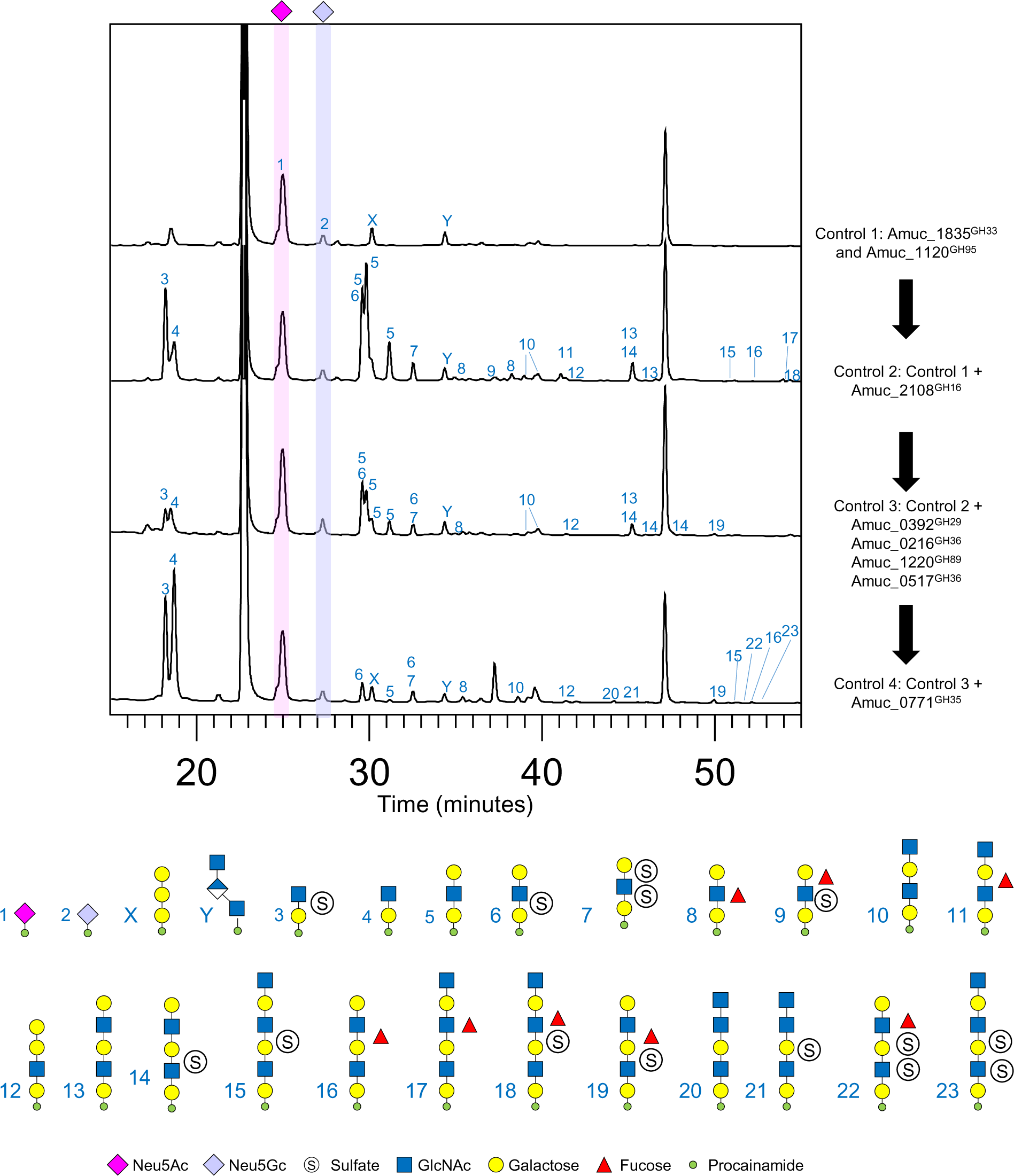
Sequential degradation of porcine gastric mucin by CAZymes from AM. Mucin was incubated with different enzymes sequentially and the reaction was stopped in between each step. The glycan products were labelled with procainamide at the reducing end and analysed by LC-FLD-ESI-MS. The results show a release of sialic acid in 1 (other monosaccharides are not resolved under these conditions) and then a release of O-glycan fragments with the addition of a GH16 endo-O-glycanase in 2. 3: The addition of a Amuc_0392^GH29^, Amuc_1220^GH89^, Amuc_0517^GH36^ and Amuc_0216^GH36^. 4: The addition of a broad-acting galactosidase (Amuc_0771^GH35^).

**Supplementary Figure 24.**
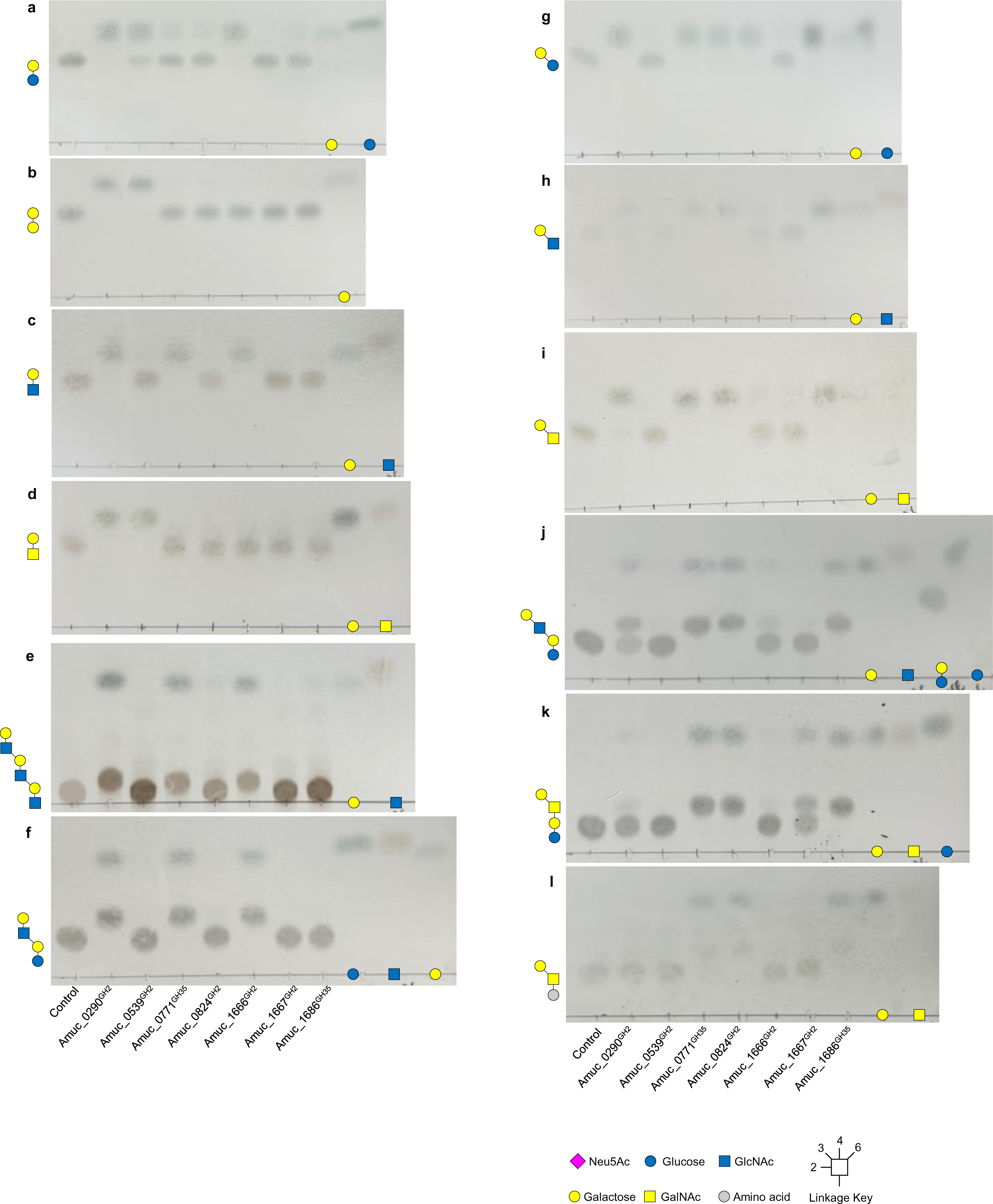
Activity of GH2 and GH35 enzymes from *A. muciniphila* BAA-835 against a wide variety of substrates. **a,** lactose. **b**, Galβ1,4Gal. **c**, LacNAc. **d**, Galβ1,4GalNAc. **e**, TriLacNAc. **f,** Lacto-N-*neo*tetraose. **g,** Galβ1,3Glc. **h,** LNB. **i,** Galβ1,3GalNAc. **j,** Lacto-N-tetrasoe. **k,** GA1. **l,** Tn antigen. Standards have also been included on the TLCs. Enzyme assays were carried out at a final substrate concentration of 1 mM, pH 7, 37 °C, overnight, and with 1 μM enzyme.

**Supplementary Figure 25.**
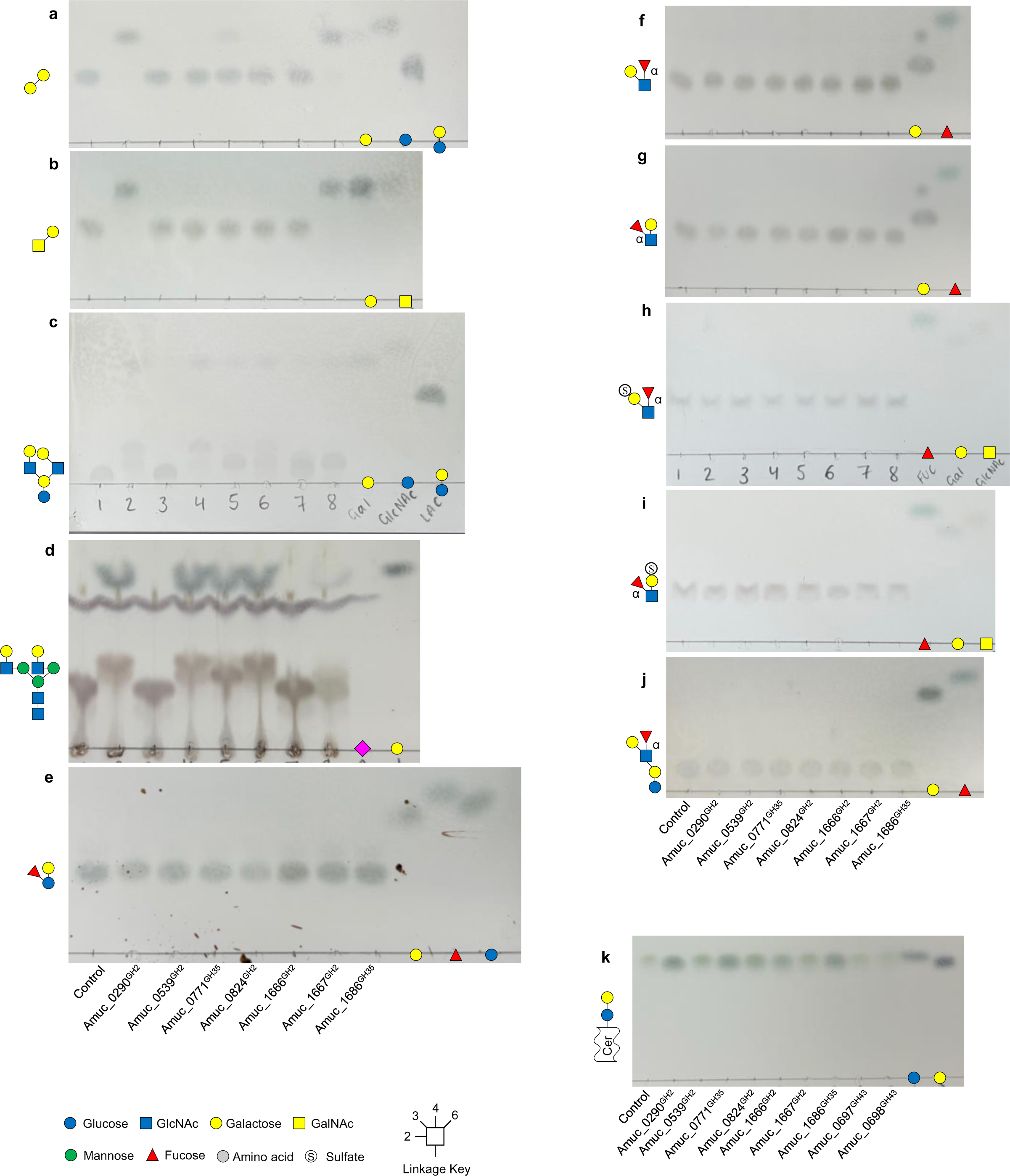
Activity of GH2 and GH35 enzymes from *A. muciniphila* BAA-835 against a wide variety of substrates. **a,** Galβ1,6Gal. **b**, Galβ1,6GalNAc. **c**, Lacto-N-*neo*hexaose. **d**, biantennary complex N-glycans. **e**, 2’FL. **f**, Lewis A. **g,** Lewis X. **h,** 3S Lewis A. **i,** 3S Lewis X. **j,** LNFP II. **k,** Lactosylceramide. Standards have also been included on the TLCs. Enzyme assays were carried out at a final substrate concentration of 1 mM (or 10mg/ml for glycoproteins), pH 7, 37 °C, overnight, and with 1 μM enzyme.

**Supplementary Figure 26.**
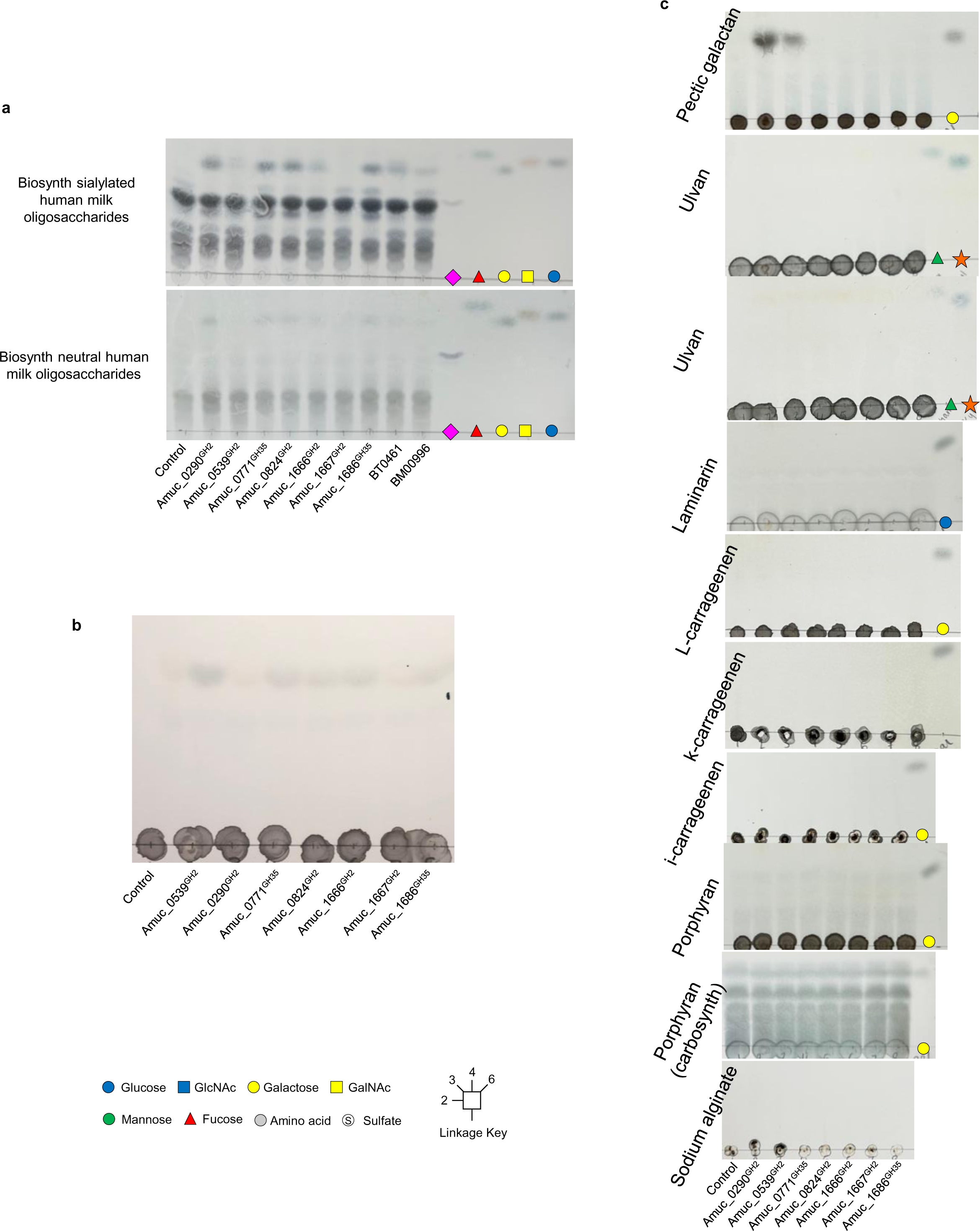
Activity of GH2 and GH35 enzymes from *A. muciniphila* BAA-835 against a wide variety of substrates. **a,** Human milk oligosaccharides. **b**, PGM III. **c**, Range of plant polysaccharides, Lactosylceramide. Standards have also been included on the TLCs.

**Supplementary Figure 27.**
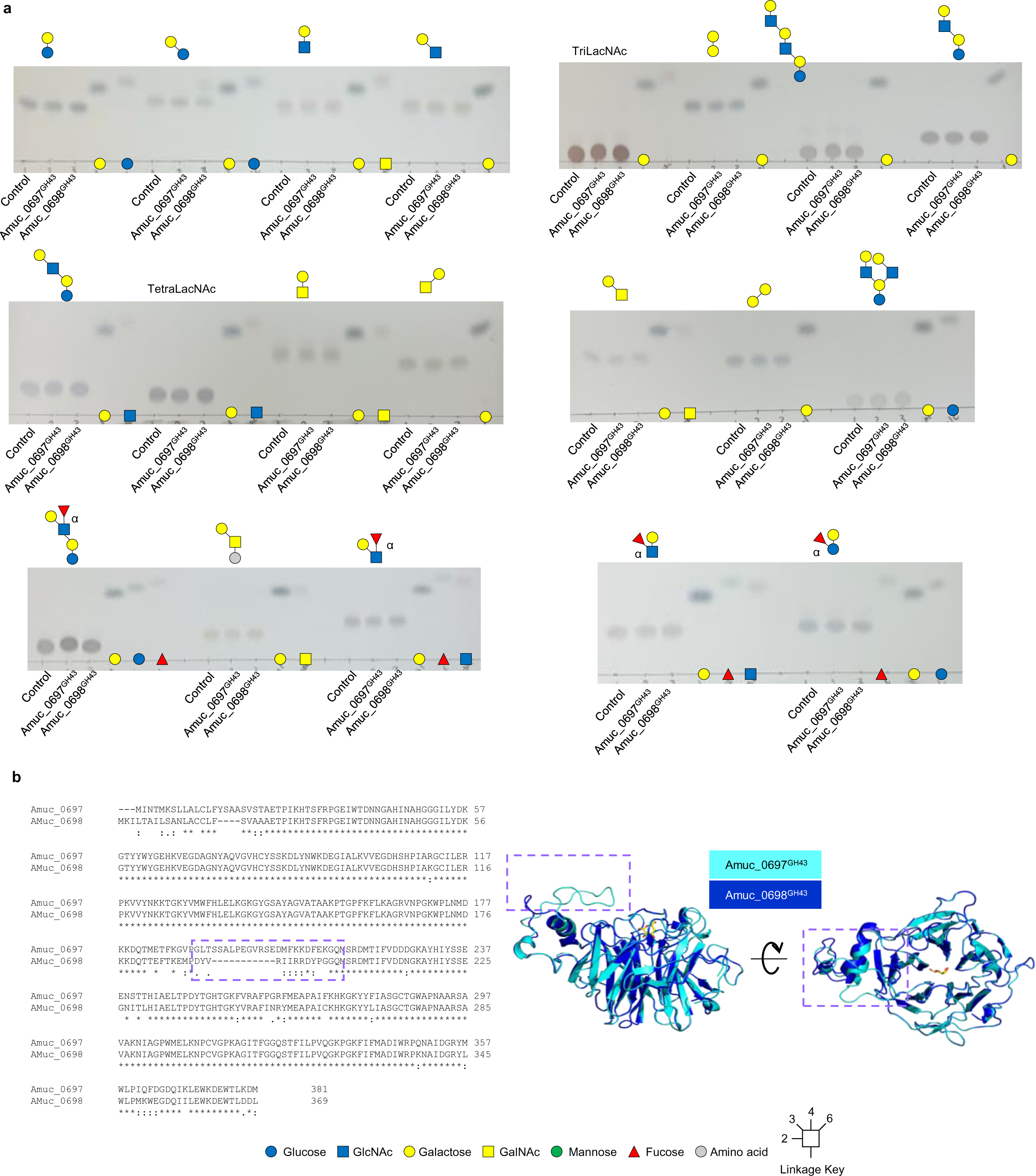
The GH43 enzymes of *A. muciniphila* BAA-835. **a,** Testing activity against a variety of defined substrates. Standards have also been included on the TLCs on the right. Enzyme assays were carried out at a final substrate concentration of 1 mM, pH 7, 37 °C, overnight, and with 1 μM enzyme. **b,** Sequence alignment (left) showing that the sequences are conserved apart from a section in the middle (purple dashed box). Right: Structural models of the two GH43 enzymes superimposed with a galactose in the active site (from 6EUI, which is BT3683 from *Bacteroides thetaiotaomicron*). The only difference between the models is one of the loops on the side of the active site and corresponds to the sequence difference seen with the alignment purple dashed box).

**Supplementary Figure 28.**
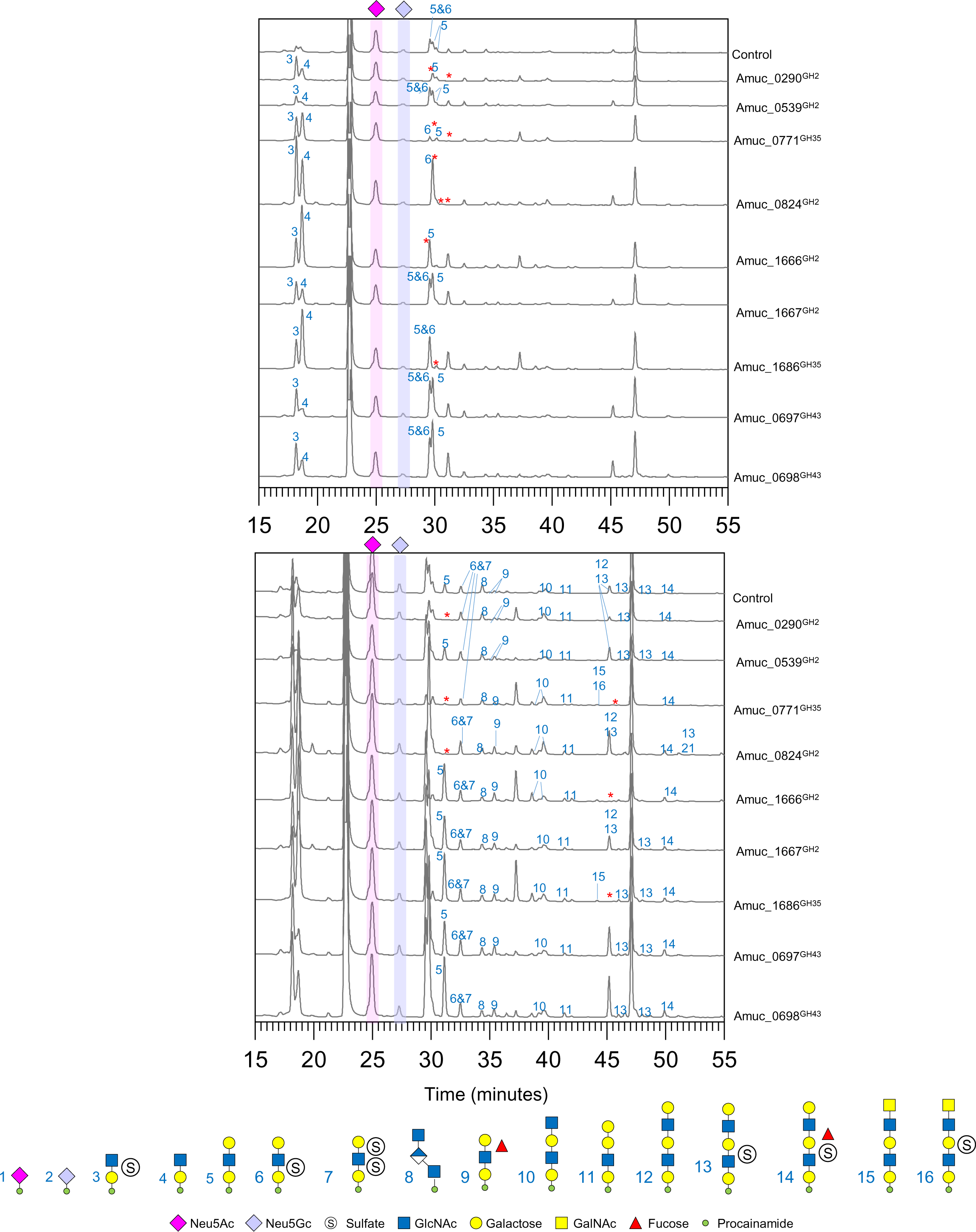
Activity of β-galactosidases against O-glycans from PGMIII analysed by LC-FLD-ESI-MS. PGMIII was pretreated sequentially with 1) sialidase and fucosidase, 2) GH16 endo-O-glycanase, and then 3) another fucosidase and three of the CAZymes that remove α-linked capping monosaccharides. The reaction was stopped in between each step and the putative β-galactosidases were added individually. The resulting O-glycans were labelled with procainamide at the reducing end and analysed by LC-FLD-ESI-MS. The chromatograms are presented in two different ways to emphasise the large peaks (top panel) and smaller peaks (bottom panel). The red asterisks indicate where a glycan is no longer present relative to the control and the β-galactosidase has broken this down.

**Supplementary Figure 29.**
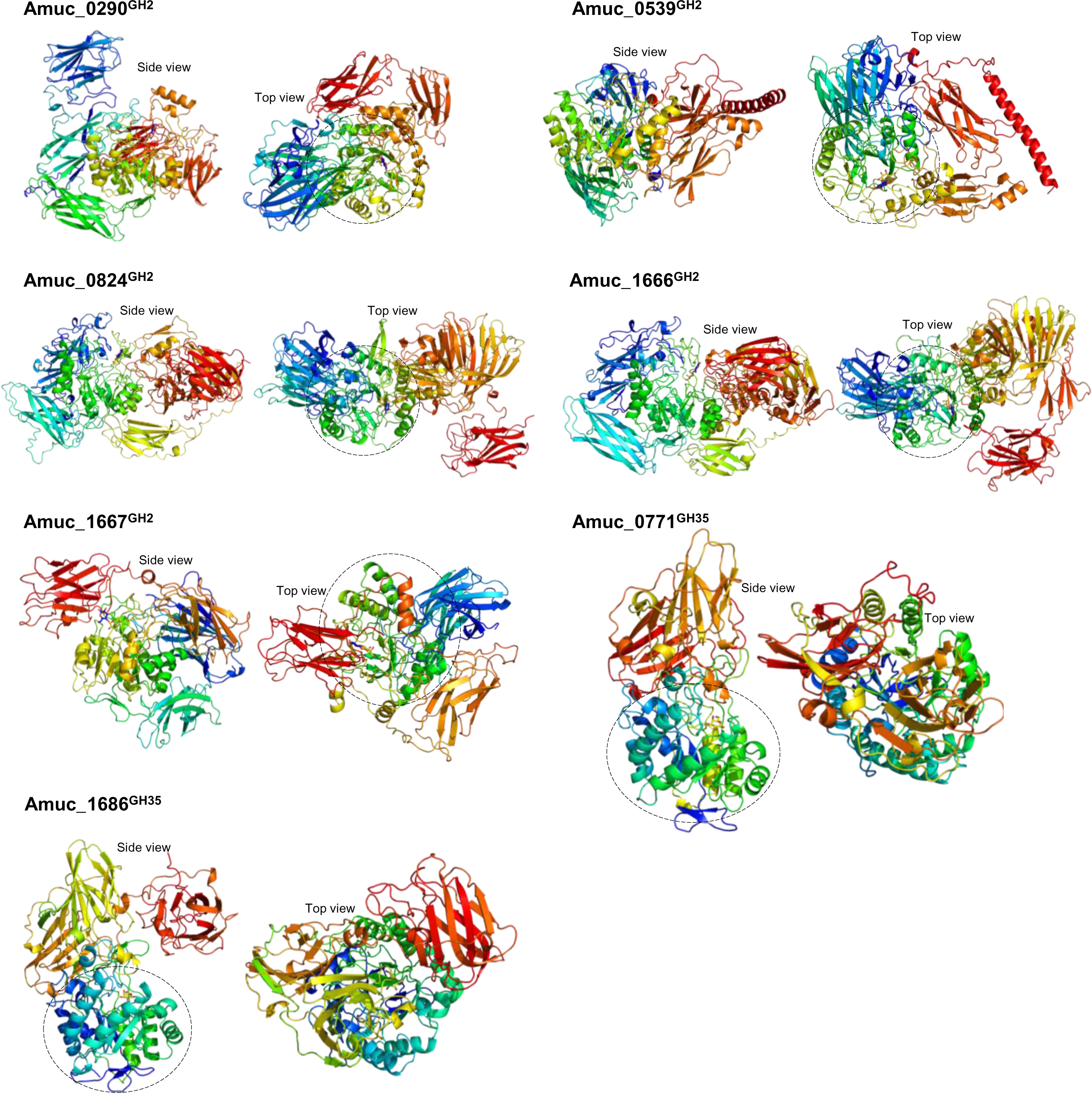
Models of the GH2 and GH35 enzymes from *A. muciniphila* BAA-835. Models were built using AlphaFold2 Colab as described in the materials and methods. Lactose from 1JYN was overlaid into the active site of the GH2 enzymes and galactose from 4E8C was overlaid into the active sites of the GH35 enzymes and are shown as sticks. The enzymes are coloured blue to red from the N- to the C-termini, respectively. The catalytic modules are circled with a dotted line to highlight the variety of different accessory modules.

**Supplementary Figure 30.**
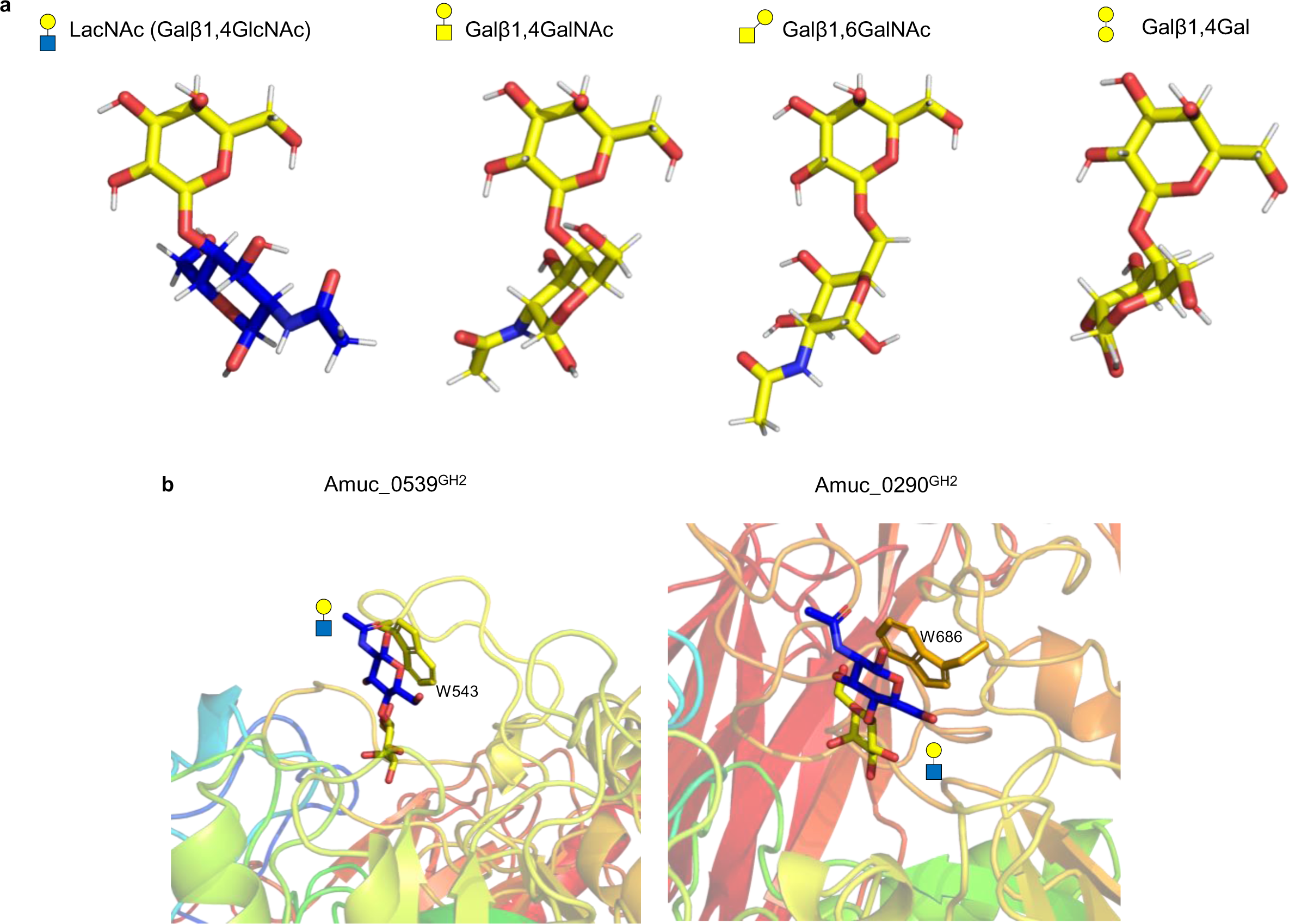
Analysis of models of the GH2 enzymes from *A. muciniphila* BAA-835. **a,** Models of different disaccharides. **b,** Models of two GH2 enzymes with LacNAc overlaid into the active site (from a crystal structure of a *Streptococcus pneumoniae* GH2, 4CUC). Aromatic residues contact the +1 sugar to likely play a role in selectivity and substrate binding.

**Supplementary Figure 31.**
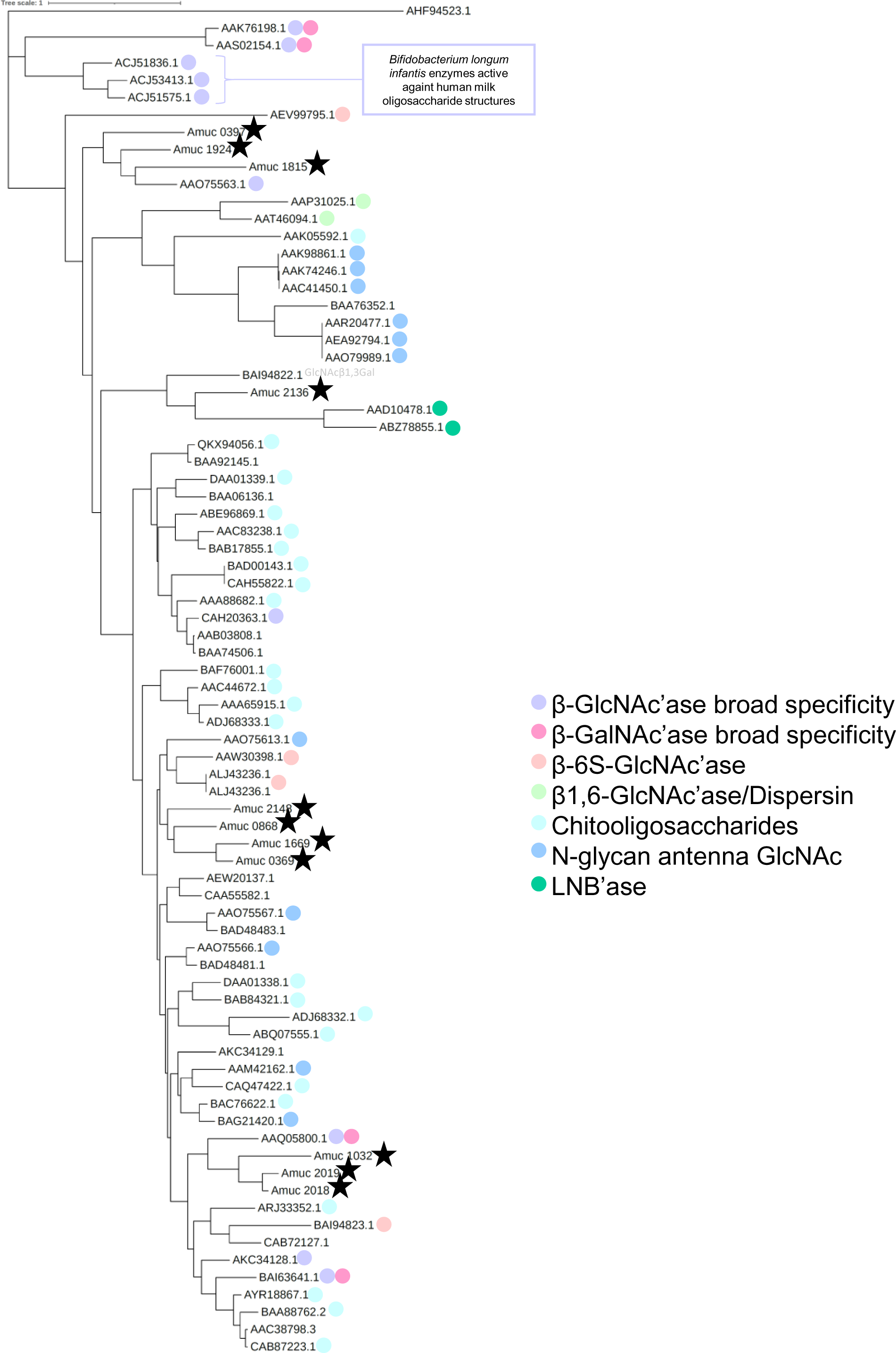
Phylogenetic tree of characterised GH20 family members with those from *A. muciniphila* ATCC BAA-835. The sequences of the GH20 family members with reported activities (CAZy database) and the ones from *A. muciniphila* ATCC BAA-835 were compared as described in the methods. The different specificities are indicated by different colours and the *A. muciniphila* ATCC BAA-835 enzymes are highlighted by black stars. Some more specific information about activity is supplied were possible. The enzymes are represented by their accession numbers of locus tags. Different specificities cluster in this analysis and the observed activity of the three *A. muciniphila* ATCC BAA-835 enzymes correlates to where they cluster on the tree.

**Supplementary Figure 32.**
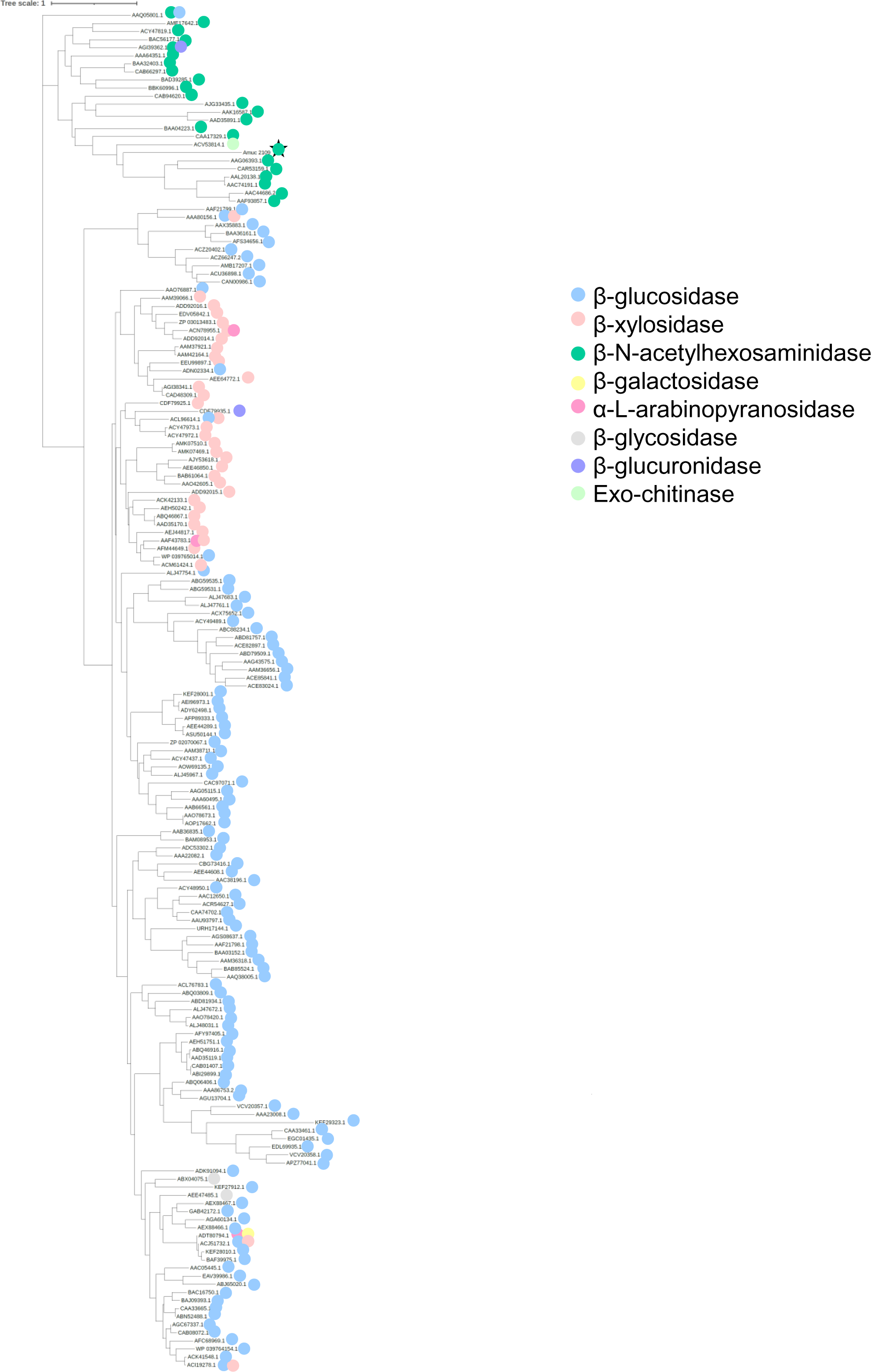
Phylogenetic tree of characterised GH3 family members with those from *A. muciniphila* ATCC BAA-835. The sequences of the GH3 family members with reported activities (CAZy database) and the ones from *A. muciniphila* ATCC BAA-835 were compared as described in the methods. The different specificities are indicated by different colours and the *A. muciniphila* ATCC BAA-835 enzymes are highlighted by black stars. The enzymes are represented by their accession numbers of locus tags. Different specificities cluster in this analysis and the observed activity of the *A. muciniphila* ATCC BAA-835 enzyme correlates to where they cluster on the tree.

**Supplementary Figure 33.**
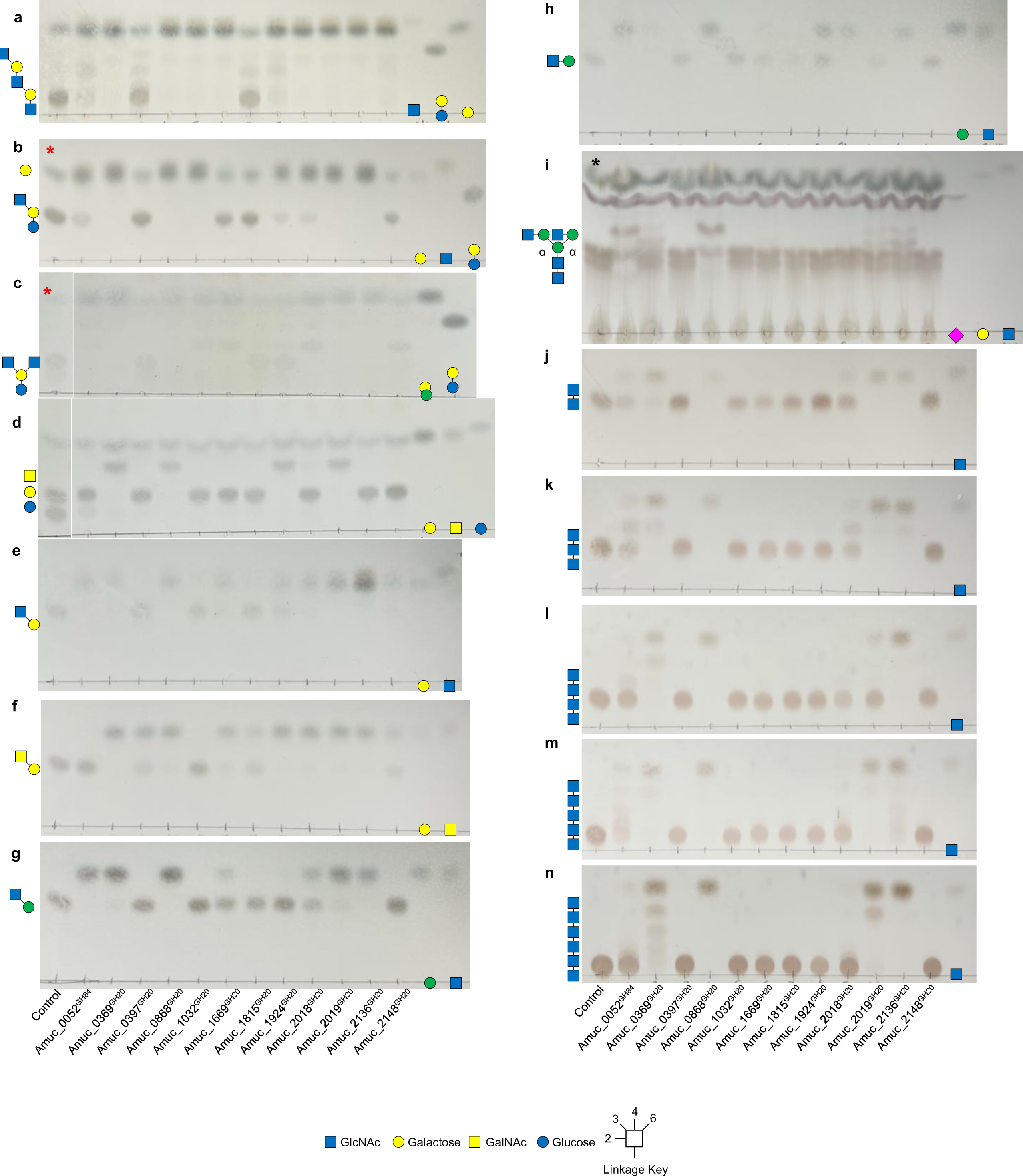
Activity of the GH20 and GH84 family members from *A. muciniphila* BAA-835 against defined oligosaccharides. Enzyme assays were carried out at a final substrate concentration of 1 mM, pH 7, 37 °C, overnight, and with 1 μM enzymes. **a,** TriLacNAc. **b**, Lacto-N-neotetraose. **c**, Lacto-N-neohexaose. **d**, Galβ1,3GalNAcβ1,4Galβ1,4Glc. **e**, GlcNAcβ1,3Gal. **f**, GalNAcβ1,3Gal. **g**, GlcNAcβ1,3Man. **h**, GlcNAcβ1,2Man. **i**, α_1_acidglycoprotein. **j**, Chitobiose. **k**, Chitotriose. **l**, Chitotetraose. **m**, Chitopentaose. **n**, Chitohexaose. Some pre-treatments were required to generate the substrates: Red asterisk - pre-treated with BT0461^GH2^ to remove the non-reducing end β1,4galactose, cyan asterisk - pre-treated with B035DRAFT_00996^GH2^ to remove the non-reducing end β1,3galactose, black asterisk – pre-treated with BT0455^GH33^, BT0461^GH2^, and PNGaseL, which are a broad-acting sialidase, β1,4-galactosidase, and remove N-glycans from the protein, respectively. The glycan that the GH20 enzymes are tested on are shown to the right of each TLC and all linkages are beta. Standards have also been included on the left of the TLCs.

**Supplementary Figure 34.**
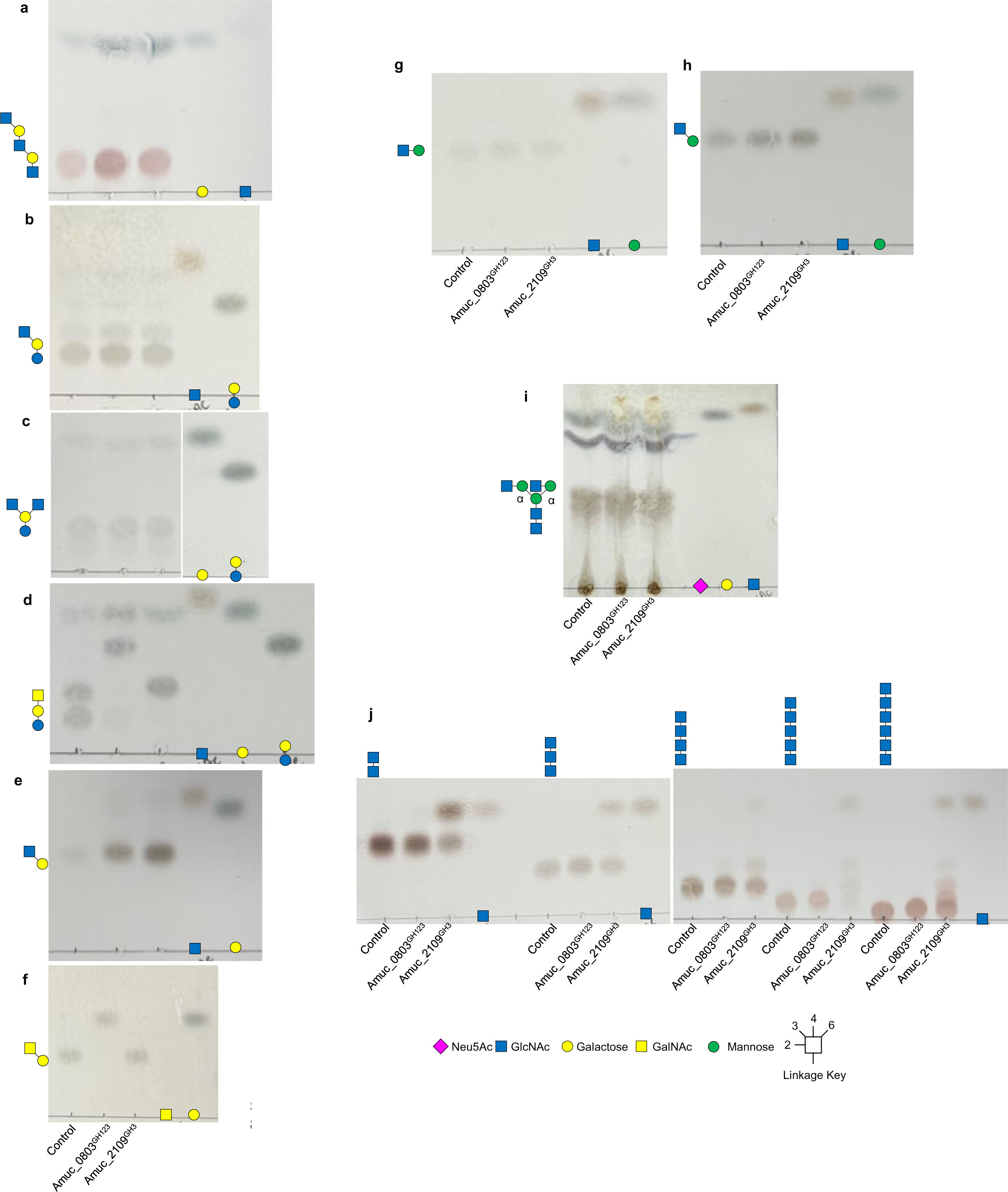
Activity of the GH123 and GH3 family members from *A. muciniphila* BAA-835 against defined oligosaccharides. Enzyme assays were carried out at a final substrate concentration of 1 mM, pH 7, 37 °C, overnight, and with 1 μM enzymes. **a,** TriLacNAc. **b**, Lacto-N-neotetraose. **c**, Lacto-N-neohexaose. **d**, Galβ1,3GalNAcβ1,4Galβ1,4Glc. **e**, GlcNAcβ1,3Gal. **f**, GalNAcβ1,3Gal. **g**, GlcNAcβ1,3Man. **h**, GlcNAcβ1,2Man. **i**, α_1_acidglycoprotein. **j**, Chitobiose-Chitohexaose, left to right. Some pre-treatments were required to generate the substrates: Red asterisk - pre-treated with BT0461^GH2^ to remove the non-reducing end β1,4galactose, cyan asterisk - pre-treated with B035DRAFT_00996^GH2^ to remove the non-reducing end β1,3galactose, black asterisk – pre-treated with BT0455^GH33^, BT0461^GH2^, and PNGaseL, which are a broad-acting sialidase, β1,4-galactosidase, and remove N-glycans from the protein, respectively. The glycan that the GH20 enzymes are tested on are shown to the right of each TLC and all linkages are beta. Standards have also been included on the left of the TLCs.

**Supplementary Figure 35.**
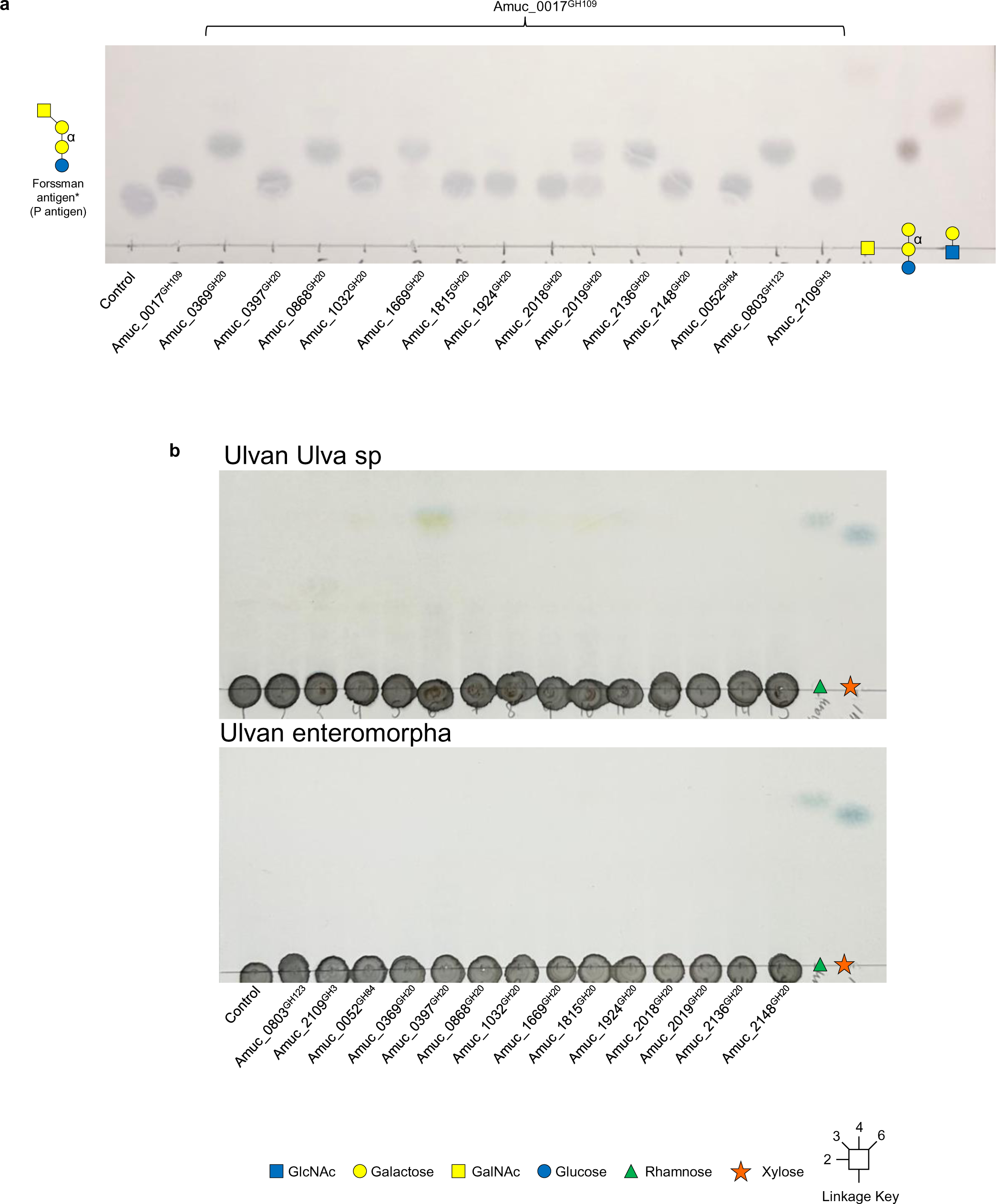
Activity of the β-HexNAc’ases from *A. muciniphila* BAA-835 against P antigen and ulvan substrates. Enzyme assays were carried out at pH 7, 37 °C, overnight, and with 1 μM enzymes. Standards have also been included on the left of the TLCs. The Forssman antigen was treated with Amuc_0017^GH109^

**Supplementary Figure 36.**
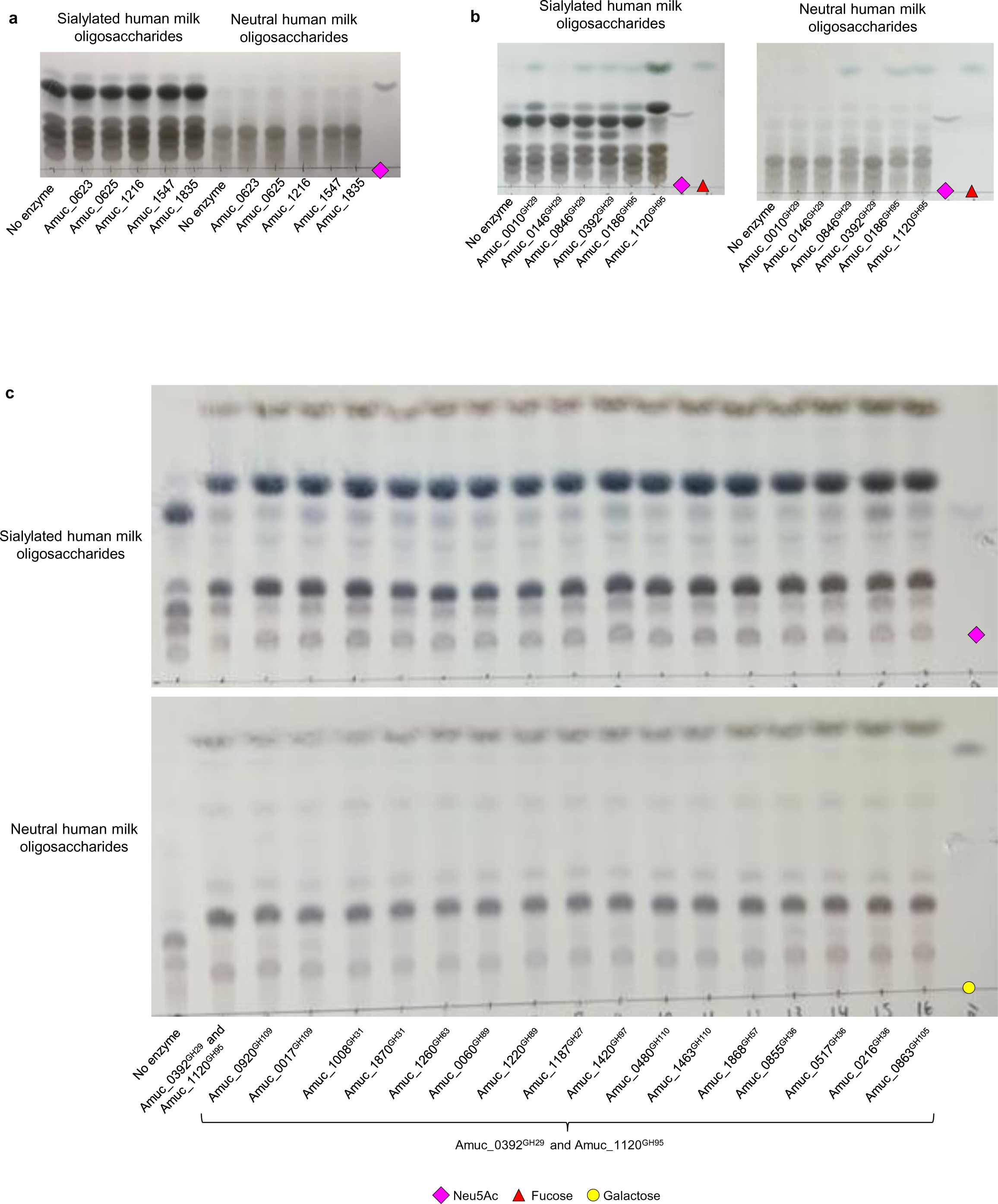
Activity of sialidases, fucosidases and other α-linkage acting GH enzymes from AM against human milk oligosaccharides. **a,** Sialidases and potential sialidases. **b**, Fucosidases. **c**, CAZymes to predicted to act on α-linked monosaccharides and these samples were also pre-treated with two fucosidases. Standards have also been included on the TLCs. Enzyme assays were carried out at a final substrate concentration of 1 mM, pH 7, 37 °C, overnight, and with 1 μM enzyme.

**Supplementary Figure 37.**
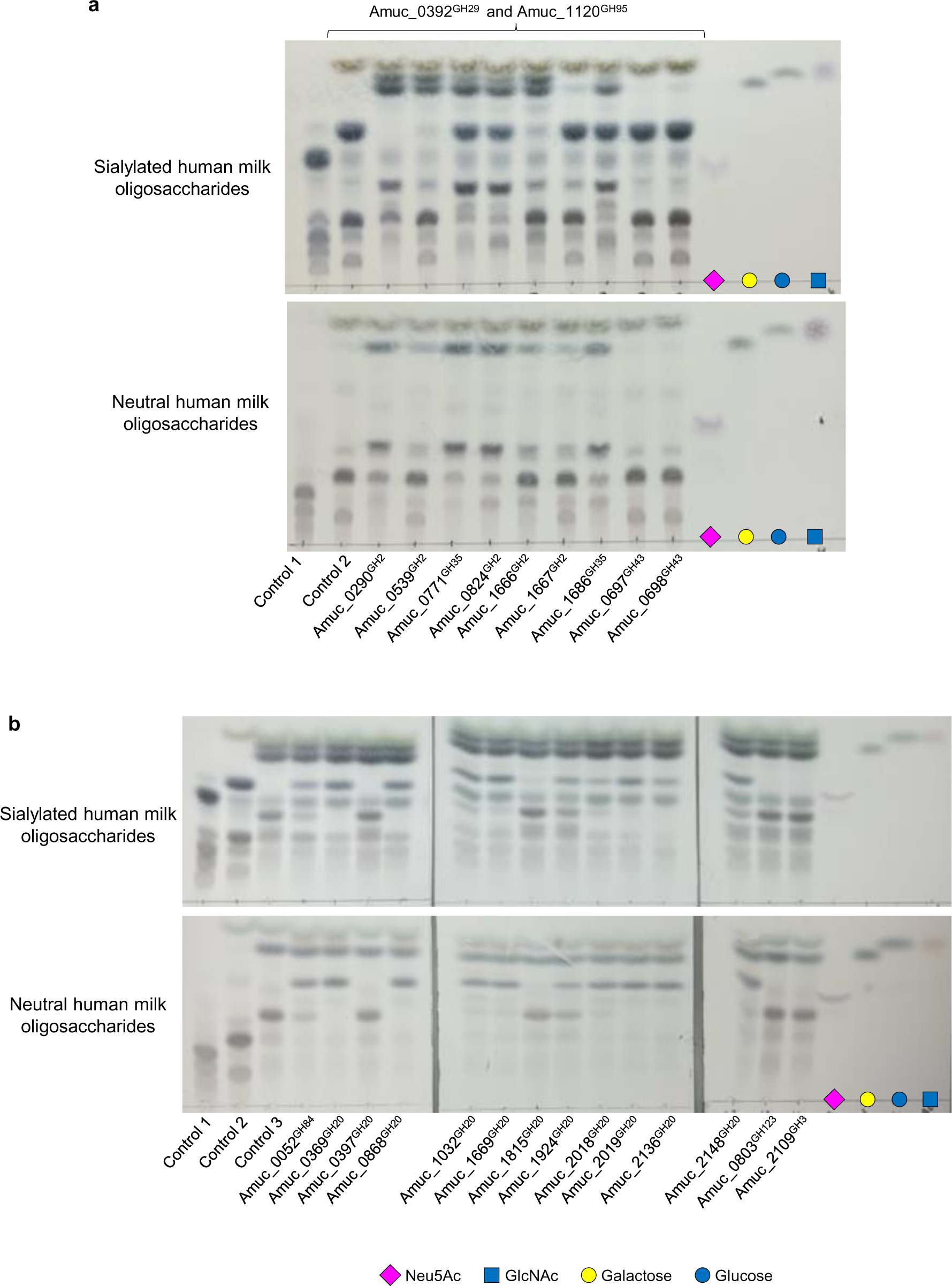
Activity of β-galactosidase and β-HexNAc’ase GH enzymes from AM against human milk oligosaccharides. **a,** β-galactosidases. Top and bottom panels are sialylated and neutral HMO preparations (Biosynth). Control 1 – no enzymes added and Control 2 - Amuc_0392^GH29^ and Amuc_1120^GH95^. The rest of the lanes are the individual enzymes with the fucosidases are also included in these reactions. **b**, β-HexNAc’ase. Top and bottom panels are sialylated and neutral HMO preparations (Biosynth). Control 1 – no enzymes added and Control 2 - Amuc_0392^GH29^ and Amuc_1120^GH95^. Control 3 – Top panel: Amuc_0517^GH36^ and Amuc_0290^GH2^. Bottom panel: Amuc_0216^GH36^ and Amuc_0771^GH35^. The controls were boiled in between all stages to produce a sequential degradation. The rest of the lanes are the individual β-HexNAc’ases. Standards have also been included on the TLCs. Enzyme assays were carried out at a final substrate concentration of 1 mM, pH 7, 37 °C, overnight, and with 1 μM enzyme.

**Supplementary Figure 38.**
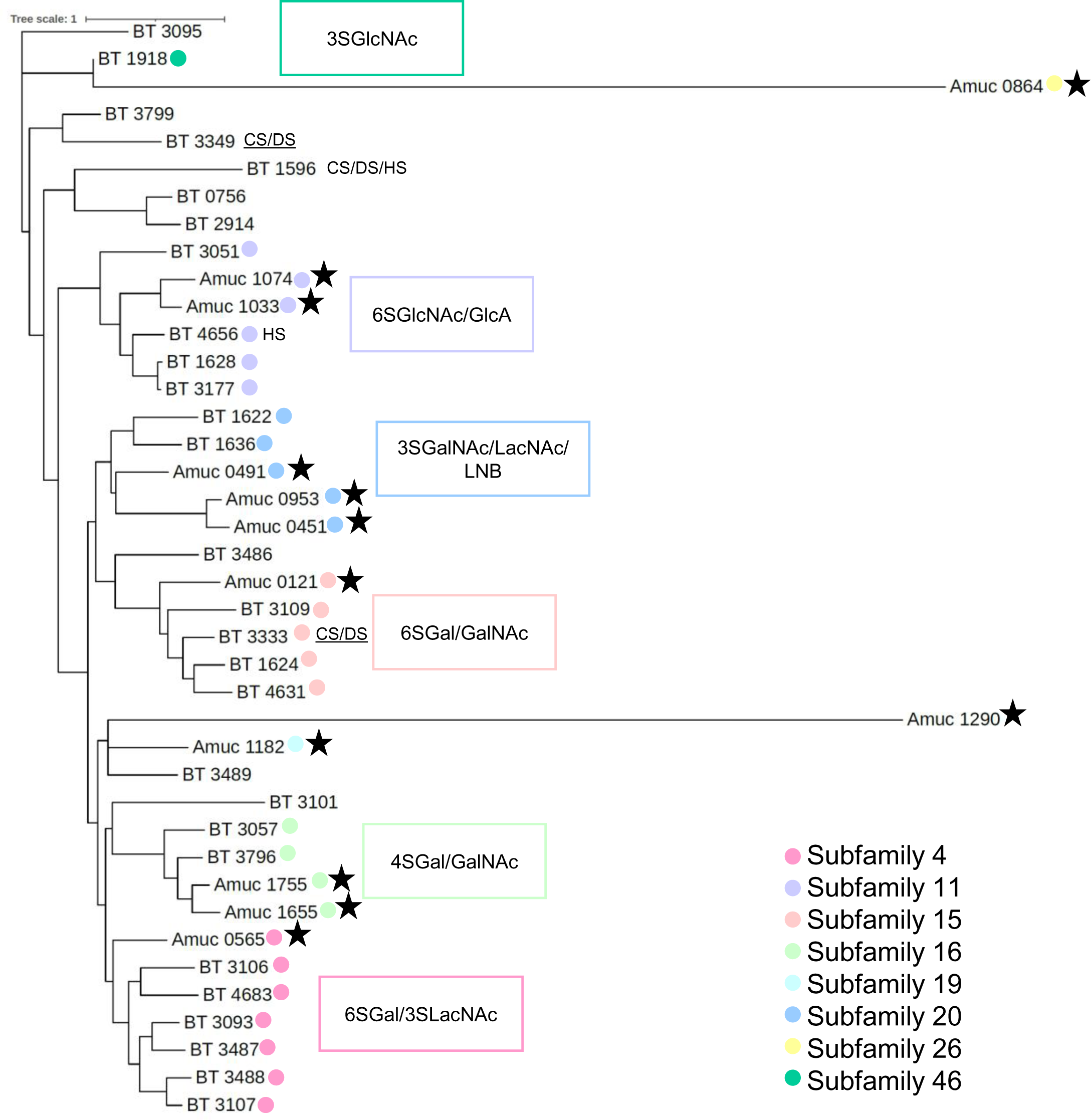
Phylogenetic tree of characterised sulfatase family members from *Bacteroides thetaiotaomicron* with those from *A. muciniphila* ATCC BAA-835. The sequences of the sulfatases with reported activities and the ones from *A. muciniphila* ATCC BAA-835 were compared as described in the methods. The different subfamilies are indicated by different colours and the *A. muciniphila* ATCC BAA-835 enzymes are highlighted by black stars. Some more specific information about activity is supplied were possible. When a sulfatase is known to act on a particular GAG, then this is noted and underlined. The enzymes are represented by their locus tags.

**Supplementary Figure 39.**
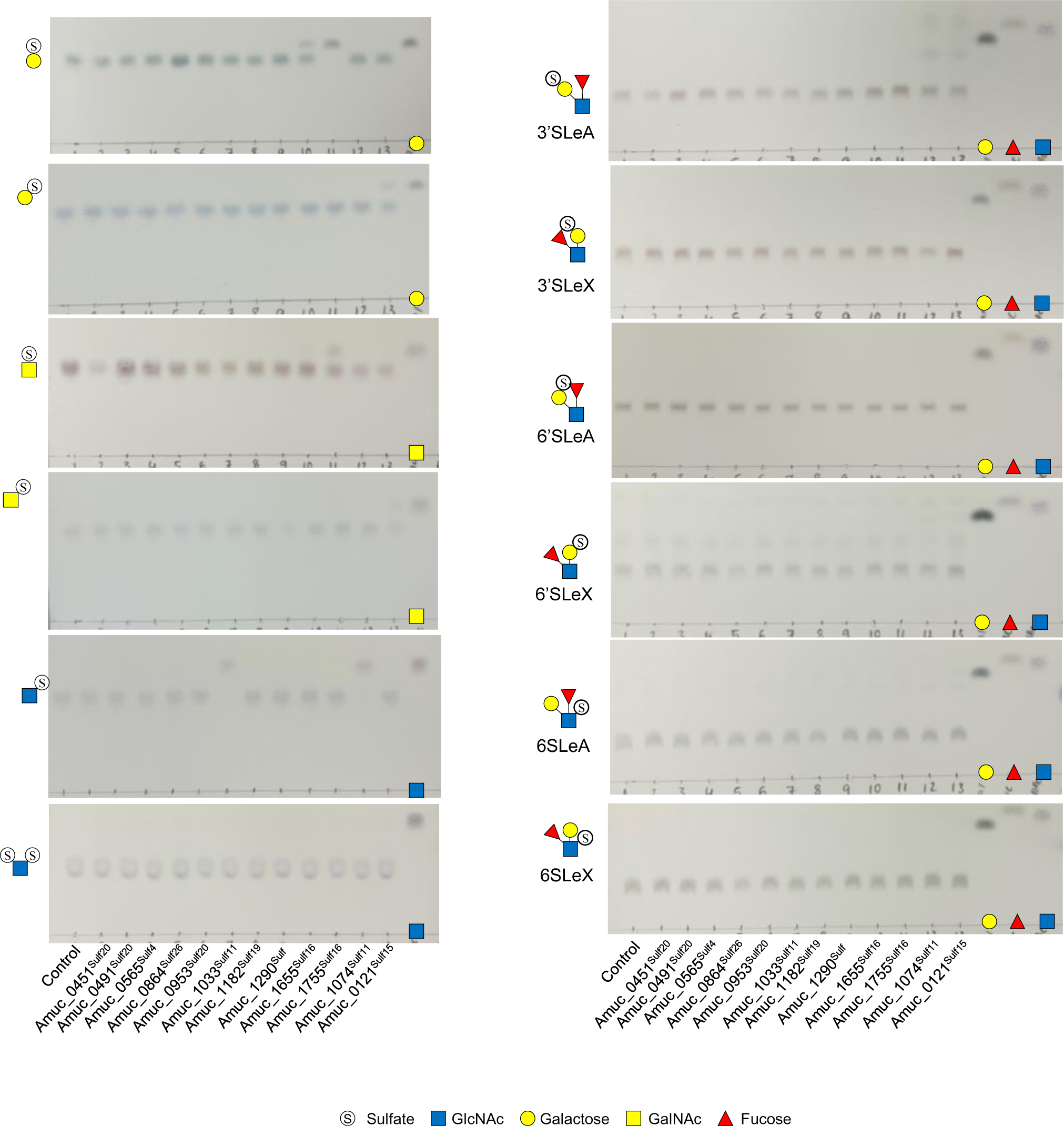
Activity of sulfatases from *A. muciniphila* BAA-835 against defined sulfated monosaccharides and Lewis Structures. Standards have also been included on the TLCs. Enzyme assays were carried out at a final substrate concentration of 1 mM, pH 7, 37 °C, overnight, and with 1 μM enzyme.

**Supplementary Figure 40.**
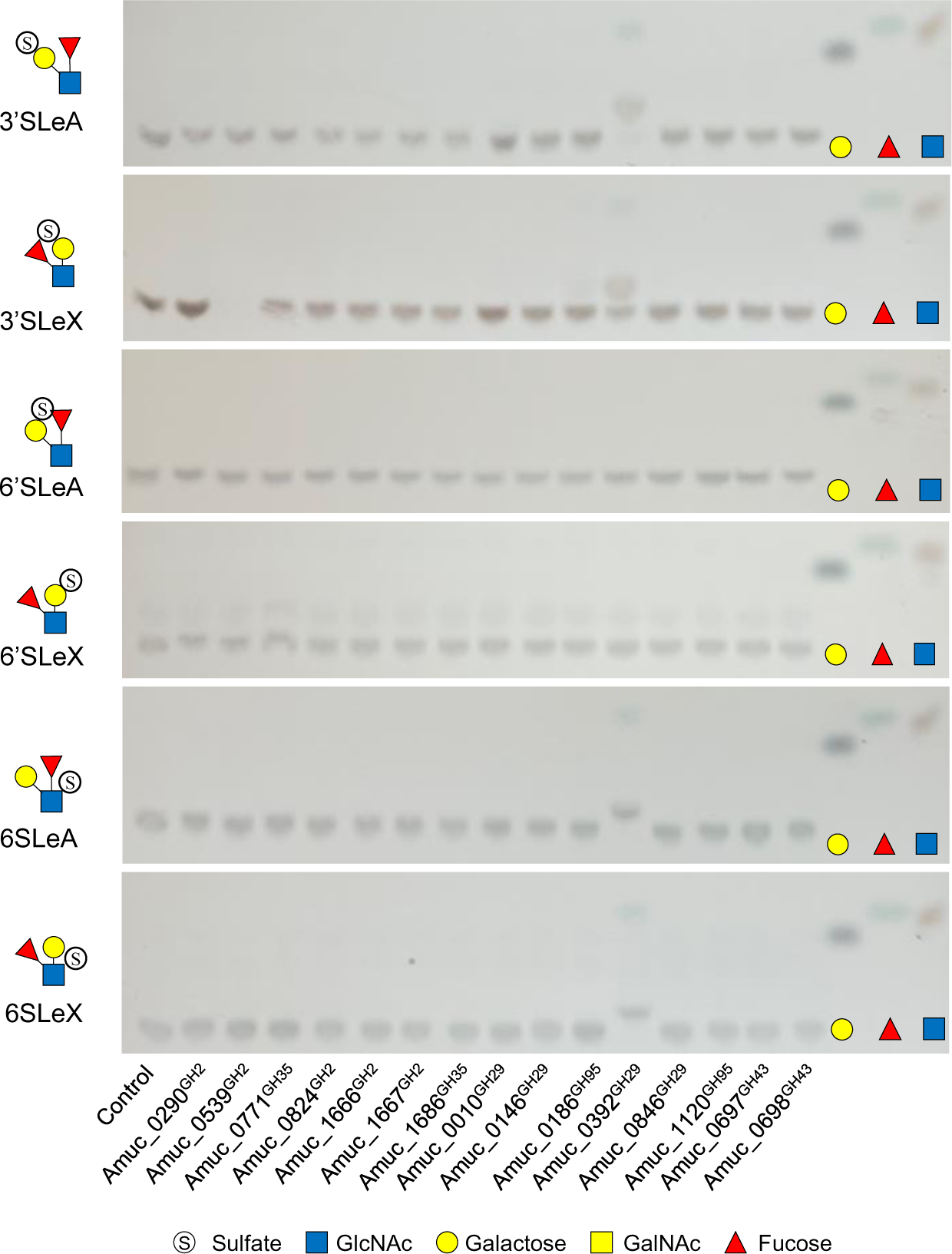
Activity of β-galactosidases and α-fucosidases from *A. muciniphila* BAA-835 against sulfated Lewis Structures. Standards have also been included on the TLCs. Enzyme assays were carried out at a final substrate concentration of 1 mM, pH 7, 37 °C, overnight, and with 1 μM enzyme.

**Supplementary Figure 41.**
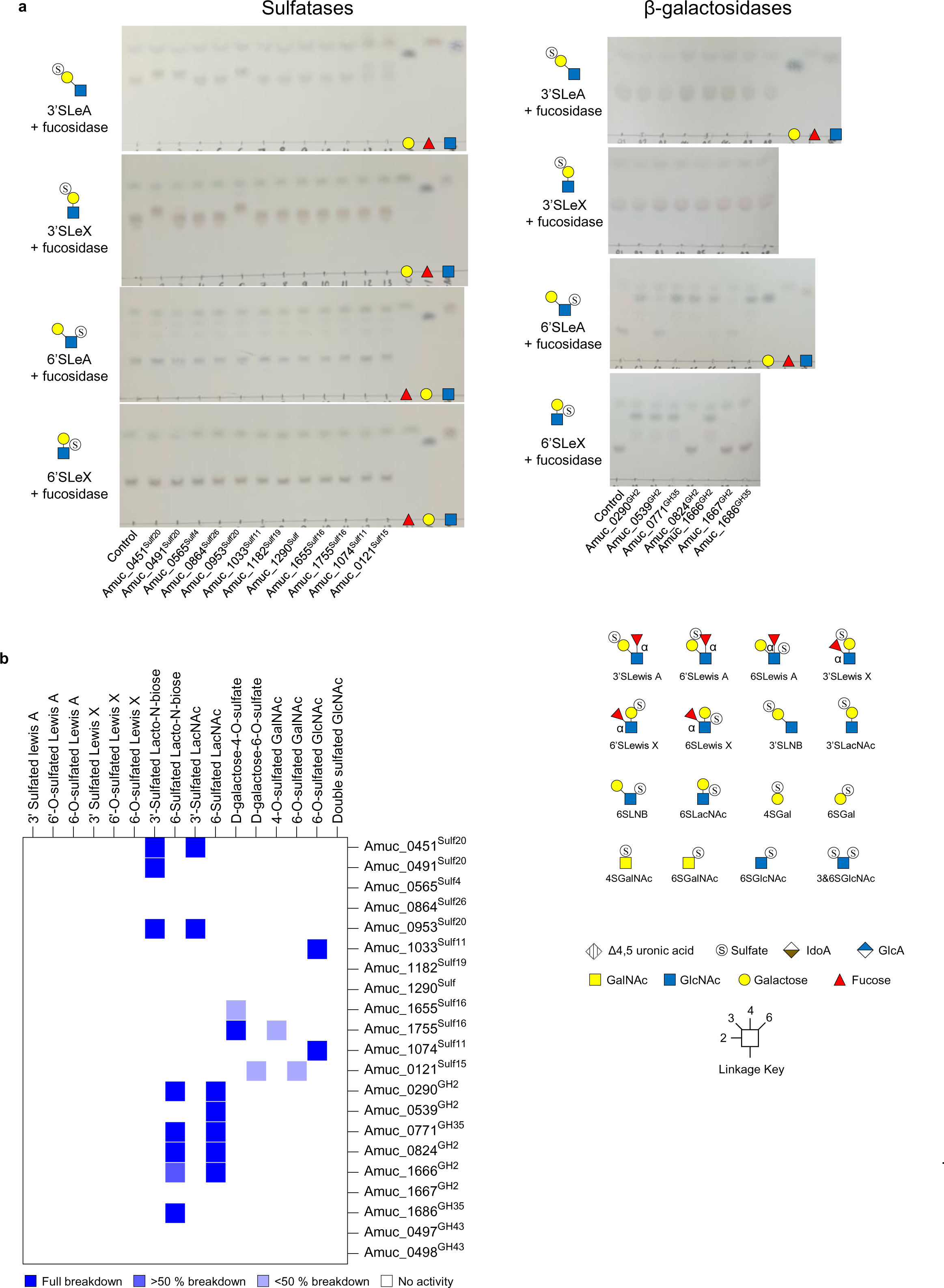
Activity of fucosidases and β-galactosidases from *A. muciniphila* BAA-835 against sulfated Lewis Structures. **a,** Standards have also been included on the TLCs. Enzyme assays were carried out at a final substrate concentration of 1 mM, pH 7, 37 °C, overnight, and with 1 μM enzyme. **a**, Heat map of recombinant enzyme activities against defined oligosaccharides. The dark blue and white indicate full and no activity, respectively, and partial activities are represented by the lighter blues.

**Supplementary Figure 42.**
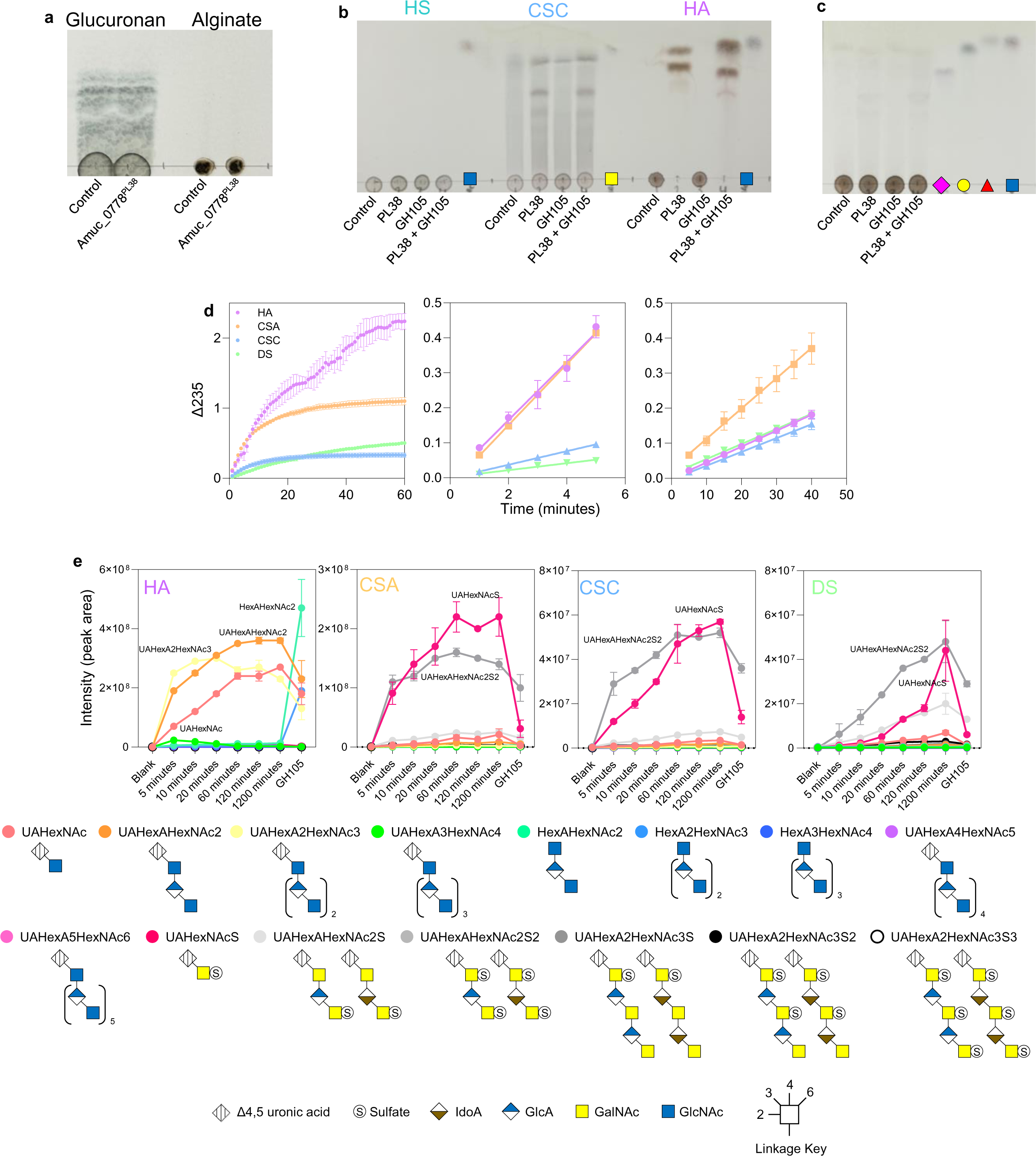
Activity of the GAG-active enzymes and breakdown of sulfated host glycans by AM. **a,** Activity of Amuc_0778^PL38^ against known PL38 substrates glucuronan (left) and alginate (right). Enzyme assays were carried out at pH 7, 37 °C, overnight, and with 1 μM enzyme. **b**, Activities of the Amuc_0771^PL38^ and Amuc_0863^GH105.^ **c**, Testing Amuc_0771^PL38^ and Amuc_0863^GH105^ against PGMIII. **d**, Left: substrate depletion of different GAGs (2 g/L) by Amuc_0771^PL38^. This wavelength allows the monitoring of the generation of unsaturated products (double bonds) of the PL38. Middle: Highest initial rates of Amuc_0771^PL38^. Right: Initial rates of the Amuc_0863^GH105^. The change in absorbance monitors the GH105 activity against the Amuc_0771^PL38^ product. **e**, Quantification of the Amuc_0771^PL38^ activity over time from four different GAG substrates by LC-MS. The area under the peaks from the chromatograms are shown and each glycan has been given a different colour. For sulfated glycans, the position of the sulfation is unknown. CSA and DS typically has 4S on GalNAc, whereas CSC has relatively high levels of 6S on GalNAc, but also low levels of 4S. The GlcA in CSC can also have 2S sulfation. HA has no sulfation.

**Supplementary Figure 43.**
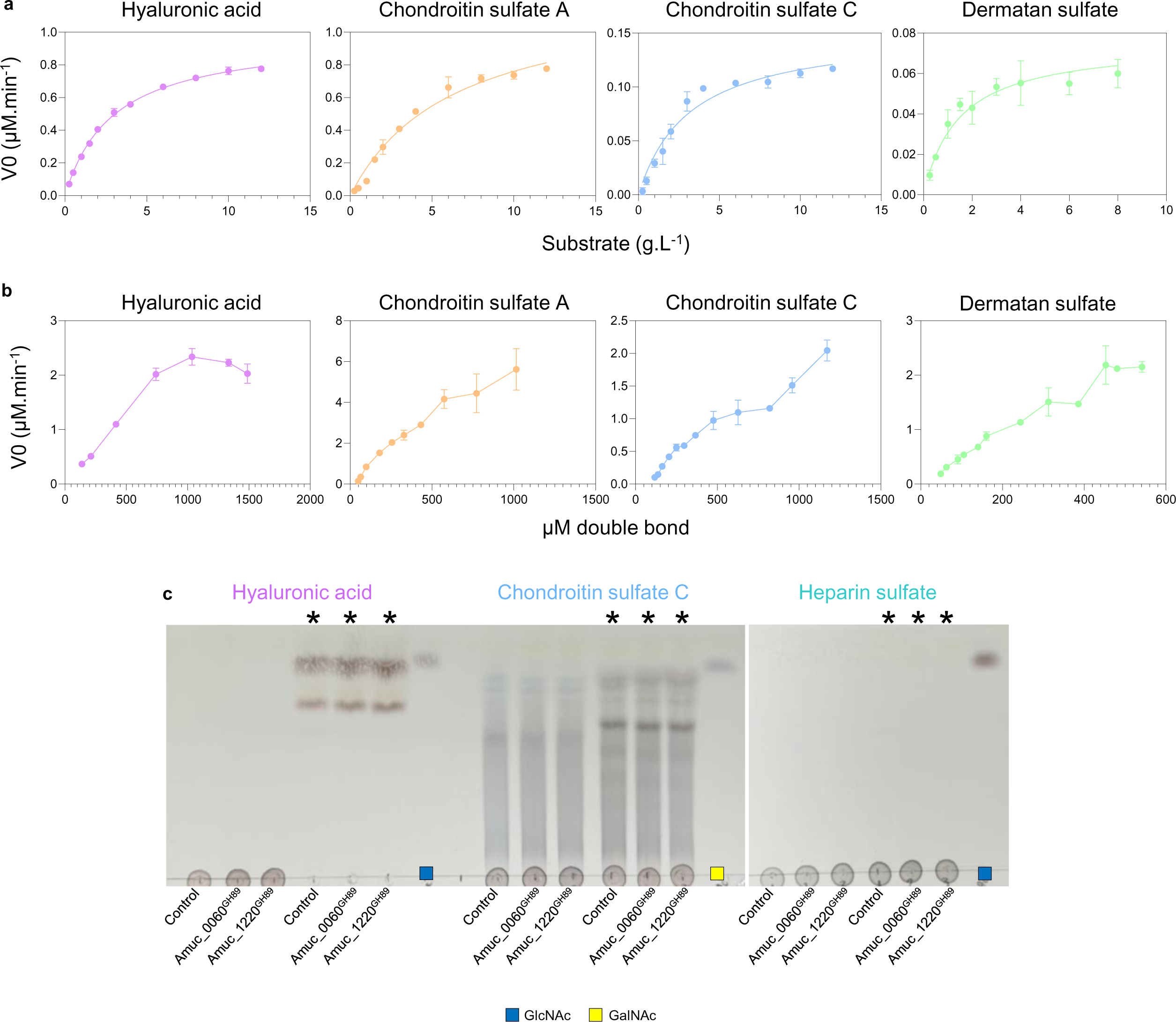
Enzyme kinetics for Amuc_0778^PL38^ and Amuc_0863^GH105^ against GAG substrates. **a,** Initial rates of Amuc_0778^PL38^. with increasing substrate concentrations. The Michaelis-Menten model was fitted in Graph-Prism. Error-bars represents standard deviations from triplicate experiments. **b,** Initial rates of Amuc_0863^GH105^ for increasing substrate concentrations. The substrate concentrations were obtained directly from the AmPL38 kinetic experiments, and the initial rate defined as loss/converted double-bond per minute. In cases were the initial absorbance of substrate exceeded 2.7 (A235) the data were omitted. No kinetic models could be fitted reliably to any of the datasets. Error-bars represent standard deviations from triplicate experiments. **c,** Testing if the GH89 enzymes have activity against GAGs. Asterisks indicate the presence of Amuc_0778^PL38^ and Amuc_0863^GH105^. Standards have also been included on the TLCs. Enzyme assays were carried out at pH 7, 37 °C, overnight, and with 1 μM enzyme.

**Supplementary Figure 44.**
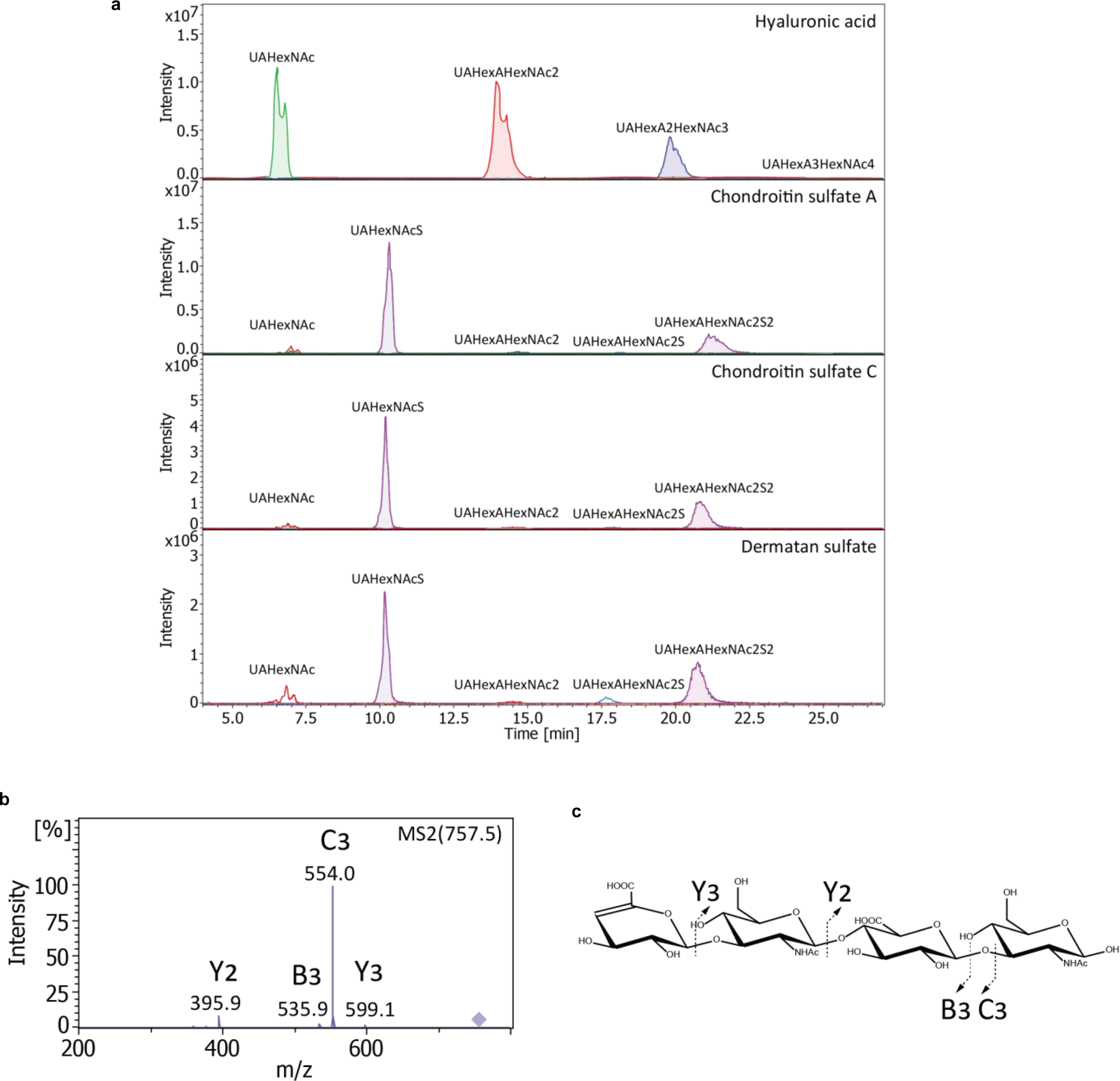
Examples of the LC-MS data of of Amuc_0778^PL38^ against GAG substrates. **a,** Extracted ion chromatograms corresponding to the most abundant compounds during Amuc_0778^PL38^ degradation of hyaluronic acid, chondroitin sulfate A, chondroitin sulfate C, and dermatan sulfate. Extracted masses: UAHexNAc2: [M-H]-378 Da, [2M-H]-757 Da; UAHexAHexNac2: [M-H]2-378 Da, [M-H]-757 Da; UAHexA2HexNac3: [M-H]2-567.5 Da, [M-H]-1136.1 Da, UAHexA3HexNac4: [M-H]2-757 Da, UAHexNAc2S: [M-H]-458 Da; UAHexAHexNac2S: [M-H]2-418 Da, [M-H]-837 Da; UAHexAHexNac2S2: [M-H]2-458 Da. **b,** Negative mode ESI MS^2^ fragmentation of UAHexAHexNAc2, a product of hyaluronic acid degradation by Amuc_0778^PL38^. Diamond indicates precursor ion at 757.5 Da. Fragments are following the nomenclature of Domon and Costello are indicated on both MS spectrum and structure. **c,** A structure of the glycan corresponding to the fragmentation pattern in ‘b’.

**Supplementary Figure 45.**
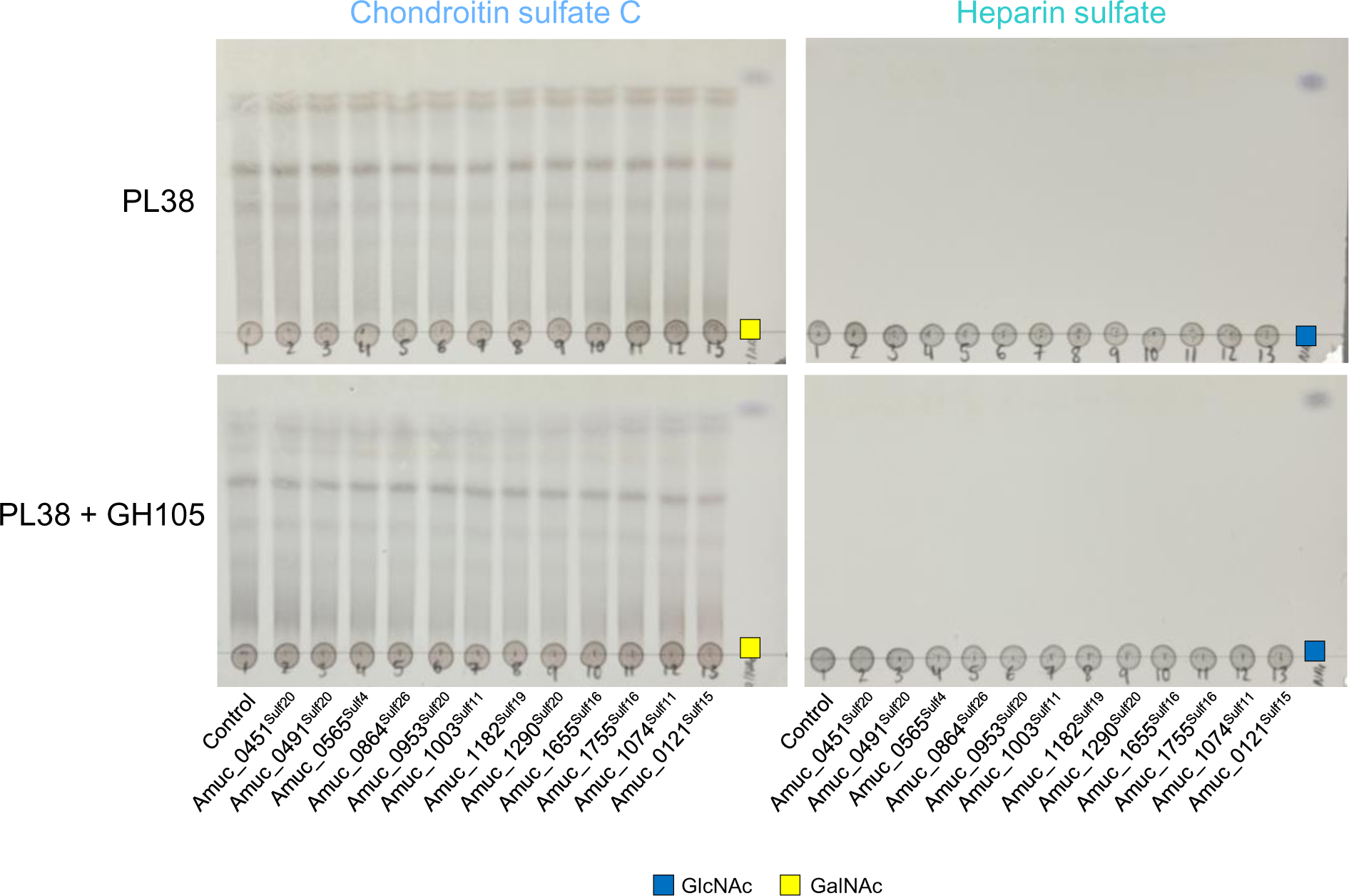
Activity of sulfatases from *A. muciniphila* BAA-835 against GAGs. The putative sulfatases were tested against two types of sulfated GAGs in the presence of Amuc_0778^PL38^ and Amuc_0863^GH105^. Standards have also been included on the TLCs. Enzyme assays were carried out at pH 7, 37 °C, overnight, and with 1 μM enzyme.

**Supplementary Figure 46.**
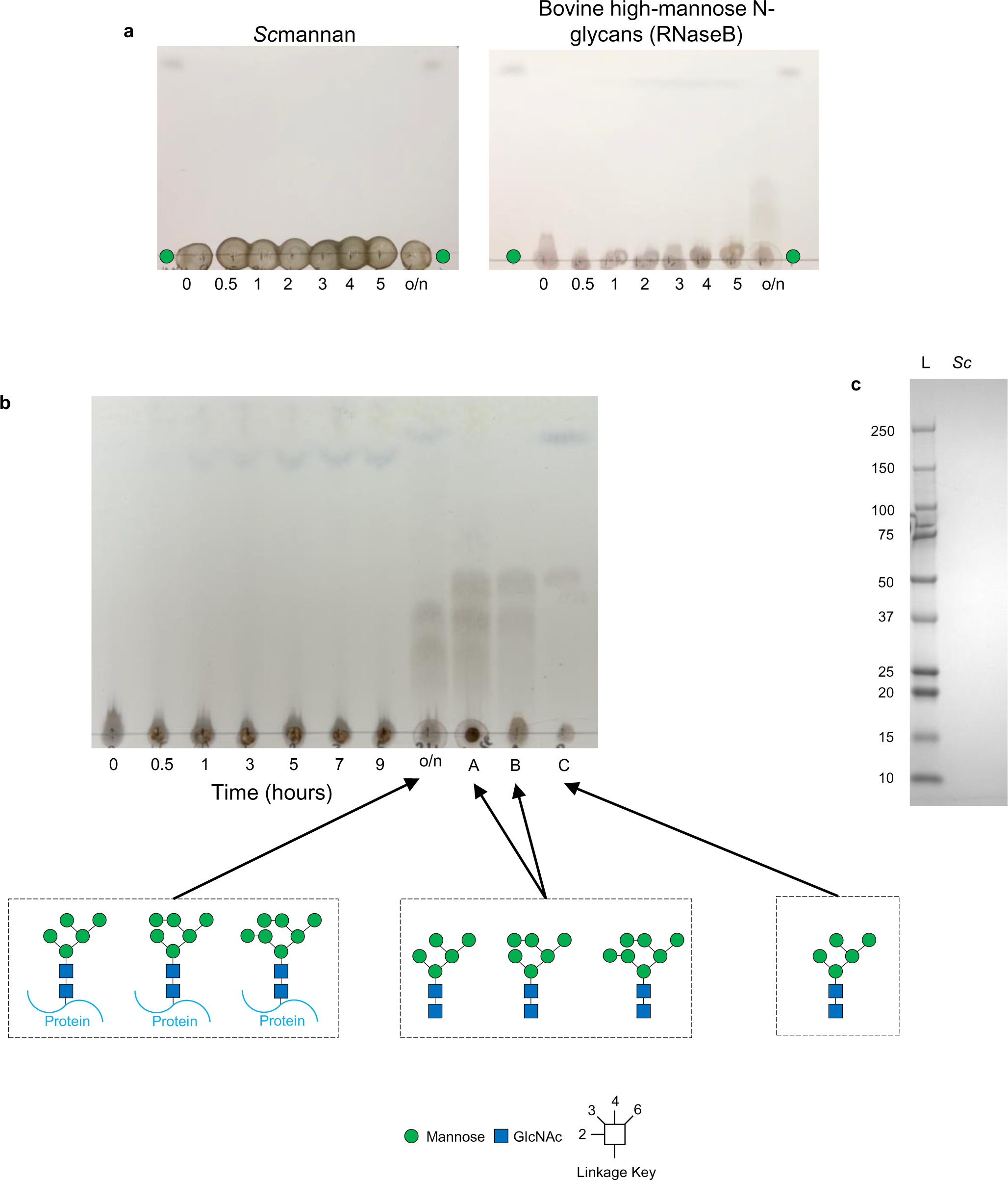
Whole cell assay with high-mannose N-glycoprotein and *Sc*mannan. **a,** The results from the whole cell assay showing that no obvious degradation of the mannan is occurring (left) and a smear appears overnight for the high-mannose N-glycoprotein (right). **b,** TLC is repeated with controls. A: the overnight whole cell assay sample treated with PNGaseL, which is a broad-acting PNGase. B: RNaseB treated with PNGaseL. The smear seen here is a mix of high-mannose N-glycans of between 5 and 7 mannose sugars. C: RNaseB treated with PNGaseL and BT3990, which is an α1,2-mannosidase that trims all the high-mannose N-glycans down to a five-mannose structure. The reaction was carried out at 37 °C, a sample removed at different time points, and boiled to stop enzyme activities. Monosaccharide standards are shown in the right lanes. All mannose linkages are alpha. **c,** SDS-PAGE gel stained for protein using Coomassie Brilliant Blue. L – ladder and *Sc* – *Sc*mannan.

**Supplementary Figure 47.**
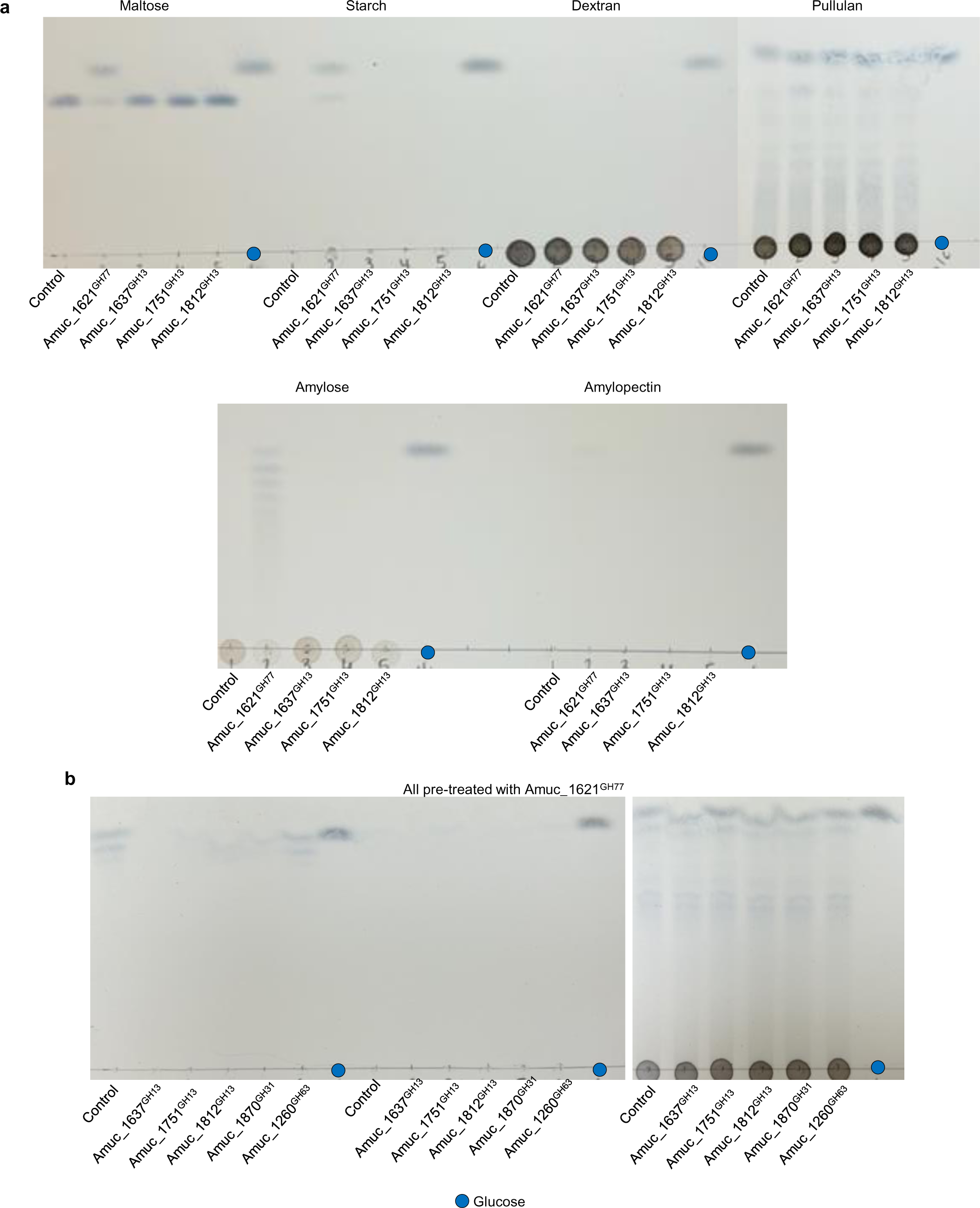
Activities of the GH13 and GH77 family members from *A. muciniphila* ATCC BAA-835 against different α-linked glucose polysaccharides. Enzyme assays were carried out at pH 7, 37 °C, overnight, and with 1 μM enzymes. **a,** Assays of the individual enzymes against possible substrates. **b**, Assays where the substrates were treated with Amuc_1621^GH77^ as well as the other enzymes. Standards have also been included on the left of the TLCs.

